# Iron Responsive Element (IRE)-mediated responses to iron dyshomeostasis in Alzheimer’s disease

**DOI:** 10.1101/2020.05.01.071498

**Authors:** Nhi Hin, Morgan Newman, Stephen Pederson, Michael Lardelli

## Abstract

**Background:** Iron trafficking and accumulation is associated with Alzheimer’s disease (AD) pathogenesis. However, the role of iron dyshomeostasis in early disease stages is uncertain. Currently, gene expression changes indicative of iron dyshomeostasis are not well characterized, making it difficult to explore these in existing datasets.

**Objective:** To identify sets of genes predicted to contain Iron Responsive Elements (IREs) and use these to explore possible iron dyshomeostasis-associated gene expression responses in AD.

**Methods:** Comprehensive sets of genes containing predicted IRE or IRE-like motifs in their 3’ or 5’ untranslated regions (UTRs) were identified in human, mouse, and zebrafish reference transcriptomes. Further analyses focusing on these genes were applied to a range of cultured cell, human, mouse, and zebrafish gene expression datasets.

**Results:** IRE gene sets are sufficiently sensitive to distinguish not only between iron overload and deficiency in cultured cells, but also between AD and other pathological brain conditions. Notably, changes in IRE transcript abundance are amongst the earliest observable changes in zebrafish familial AD (fAD)-like brains, preceding other AD-typical pathologies such as inflammatory changes. Unexpectedly, while some IREs in the 3’ untranslated regions of transcripts show significantly increased stability under iron deficiency in line with current assumptions, many such transcripts instead display decreased stability, indicating that this is not a generalizable paradigm.

**Conclusion:** Our results reveal IRE gene expression changes as early markers of the pathogenic process in fAD and are consistent with iron dyshomeostasis as an important driver of this disease. Our work demonstrates how differences in the stability of IRE-containing transcripts can be used to explore and compare iron dyshomeostasis-associated gene expression responses across different species, tissues, and conditions.

## Introduction

The pathological processes underlying Alzheimer’s Disease (AD) commence decades before symptoms become evident [1, 2]. Since the early days of AD research, iron trafficking and accumulation has been observed to be altered in AD [3–7]. However, it is still unclear whether iron dyshomeostasis represents a late symptom or an early pathological driver of AD. Iron homeostasis is closely linked to many critical biological processes including cellular respiration, hypoxia, and immune responses, all of which are disrupted in AD. Consistent with this, discoveries over the past decade have shown that disruptions to iron homeostasis can drive feedback loops that worsen AD pathology [8, 9]. However, evidence for iron dyshomeostasis as an early driver of the disease is only just emerging. While iron accumulation associated with amyloid plaque formation has been observed in various transgenic mouse models of AD [10–13], the age at which iron dyshomeostasis first occurs is uncertain [14].

Currently, gene expression patterns representing responses to iron dyshomeostasis are not well-characterized. Cellular responses to iron dyshomeostasis are complex and involve several systems and layers of regulation. The stability of the transcription factor HIF1*α*, (a component of HIF1, a master regulator of responses to hypoxia) is regulated by an iron-dependent mechanism so that transcriptional responses to iron deficiency can resemble hypoxia responses [15]. However, cellular iron homeostasis is also regulated at the post-transcriptional level by the IRP/IRE system [16–18]. In this system, altered levels of available ferrous iron (Fe^2+^) cause Iron Regulatory Proteins (IRP1 or IRP2) to change conformation or stability respectively [19, 20]. This alters their ability to bind *cis*-regulatory Iron Responsive Element (IRE) stem-loop motifs in the 3’ or 5’ untranslated regions (UTRs) of genes encoding products related to iron metabolism. Only a few IRE-containing genes have been characterized in detail, including transferrin receptor 1 (*TFR1*; 3’ UTR IRE), divalent metal transporter 1 (*DMT1* or *SLC11A2*; 3’ UTR IRE), ferritin heavy chain and ferritin light chain (*FTH1* and *FTL*; both 5’ UTR IRE), and ferroportin (*SLC40A1*; 5’ UTR IRE) [18]. In general, these previously-characterized IRE-containing genes suggest that IRPs binding to 3’ IREs tend to stabilize transcripts to increase protein translation, while binding to 5’ IREs suppresses translation [17]. It is important to note that while the ancestral ferritin IRE is present even in lower metazoans (including sponges), IRE elements in other genes have likely arisen more recently through convergent evolution [21]. Perhaps because of this, the properties of IRE genes can vary in different ways. For example, *SLC11A2* is stabilized by IRP1 binding to a greater extent than by IRP2 binding (as its 3’ IRE has a greater affinity for IRP1) [22]. Similarly, while *CDC42BPA* and *TFRC* both possess 3’ IREs, *CDC42BPA* is stabilized to a greater extent than *TFRC* under iron deficiency [23]. There are also IRE-containing genes that bind only IRP1 or IRP2 exclusively, at least in mice [24]. However, global gene expression changes mediated by genes with IREs have not yet been well-defined, and the overall expression patterns of IRE-containing genes have not been explored in the context of AD. In addition, it is unclear how expression of these genes might differ between AD and other neurodegenerative diseases, or how AD risk factors such as aging and hypoxia might contribute.

In this study, we utilized the SIREs (Searching for Iron Responsive Elements) tool [25] to predict and identify sets of IRE-containing genes in human, mouse, and zebrafish. We then applied these gene sets to explore overall IRP/IRE-mediated iron dyshomeostasis- associated responses in datasets involving: (1) a cultured cell line subjected to iron overload and deficiency treatments, (2) a cohort of AD patients, healthy controls, and two other pathological conditions affecting the brain, (3) 5XFAD mice used to model the amyloid and tau pathology seen in AD, and (4) a zebrafish knock-in model possessing a familial AD (fAD)-like mutation.

Our IRE gene sets displayed significant enrichment in AD, the 5XFAD mouse model, and an fAD-like zebrafish model, demonstrating for the first time the early and extensive involvement of IRP/IRE-mediated iron dyshomeostasis-associated gene expression responses in the context of AD. IRE gene sets were sufficiently sensitive to distinguish not only between iron overload and deficiency in a cultured cell line dataset, but also between AD and other pathological conditions affecting the brain (pathological aging and progressive supranuclear palsy), implying that the dysregulation of IRE-containing genes in AD may differ from other conditions. Changes in IRE gene expression were already evident in young adult brains of animal models of AD (3-month-old 5XFAD mice, 6-month-old fAD-like zebrafish). Overall, our observations do not support the current assumption that IRP binding to 3’ IREs generally stabilizes transcripts as, under most conditions, we observed both increases and decreases in the abundances of transcripts with either 3’ or 5’ IREs.

## Methods

### Defining IRE gene sets for human, mouse, and zebrafish

We extracted all 3’ and 5’ UTR sequences from the human, mouse and zebrafish genome assemblies using the Bioconductor packages BSgenome.Hsapiens.UCSC.hg38, BSgenome.Mmusculus.UCSC.mm10 and BSgenome.Drerio.UCSC.danRer11, and gene definitions from the Ensembl 94 release. Note that potential IREs have also been identified within protein coding sequences [24, 26], but were not included in our analyses under the initial assumption that the classic IRE paradigm was valid, as their possible effects on transcript stability are unknown. Each set of UTR sequences was then submitted into the SIREs web server (v.2.0 [25]). The SIREs algorithm assigns quality scores to predicted IREs taking into account whether the sequence is canonical, whether it contains any mismatches and bulges, and the free energy of the secondary structure. Canonical sequences are tagged by SIREs as being high-quality, while IRE-like sequences are tagged as low or medium quality. Given that high-quality IRE predictions miss the majority of true IREs, this enables a more comprehensive sampling of IRE motifs. For human, mouse, and zebrafish, we separately defined the following four gene sets: ***HQ 3’ IREs*** and ***HQ 5’ IREs*** (representing genes with high-quality predicted IREs), along with ***all 3’ IREs*** and ***all 5’ IREs***, which included genes containing any predicted IRE in the respective UTR. Comparisons between gene sets were performed using the UpSetR package (v.1.4.0 [27]) with gene ID mappings between species obtained by BioMart [28].

### Over-representation of IRE gene sets in existing MSigDB gene sets

We downloaded the following gene set collections from MSigDB (v.6.0 [29]): Hallmark, C2 (gene sets from online pathway databases and biomedical literature, including KEGG and the REACTOME databases), C3 (motif gene sets based on regulatory targets), and C5 (gene sets defined by Gene Ontology terms). We excluded the following collections from analysis: C1 (specific to human chromosomes and cytogenetic bands, while our analysis involves different species), C4 (computationally-defined cancer-focused gene sets), C6 (oncogenic signatures) and C7 (immunologic signatures). C4, C6, and C7 were not included as the level of detail in the gene sets in these specific collections is more domain-specific rather than broad-level. We used Fisher’s exact test to determine whether any IRE gene set was significantly over-represented in each MSigDB gene set. Gene sets were defined as having significant enrichment for IRE gene sets if the FDR-adjusted *p-*value from Fisher’s exact test was below 0.05. UpSet plots were produced using the UpSetR package (v.1.4.0 [27]) while network representations were produced in Gephi (v.0.9.3 [30]). To produce network visualizations, we first exported node and edge tables from R. The nodes table contained the following gene sets: top 15 gene sets (ranked by Fisher’s exact test *p*-value), Hallmark Heme Metabolism gene set, ***all 3’ IREs***, ***all 5’ IREs***, and all genes contained within these gene sets. The edges table contained gene – gene set edges which indicated the gene set(s) that genes belonged to. To create the network plots in Gephi, we used “Force Atlas 2” as the initial layout algorithm, followed by the “Yifan Hu” [31] layout algorithm to improve visual separation between distinct groupings of genes.

### Over-representation analysis of transcription factor motifs in IRE gene promoters

Defining promoter regions as being 1500 bp upstream and 200 bp downstream of the transcription start site for each gene, we used the findMotifs.pl script from HOMER (v.4.11) [32, 33] to search for known transcription factor binding site (TFBS) motifs in the promoters of each IRE gene set. The HOMER Motif database contains 363 vertebrate transcription factor binding motifs based on analysis of high-quality public ChIP-seq datasets (http://homer.ucsd.edu/homer/motif/HomerMotifDB/homerResults.html). We considered TFBS motifs as being significantly enriched in a gene set if the FDR-adjusted *p*-value (q-value as provided in HOMER output) was less than 0.05.

### Gene set enrichment testing

We performed all gene set enrichment tests in R v3.6.1 [34] using *fry* [35, 36], *camera* [37], and *fgsea* [38, 39]. For *fry*, and *camera,* we used model fits obtained using *limma* [40, 41], whilst for *fgsea*, a ranked list was obtained using *moderated t-*statistics taken from *limma*. All genes were used in gene set enrichment tests (i.e. not just DE genes). We combined the raw *p*-values from *fry, camera*, and *fgsea* using Wilkinson’s method [42] with default parameters, followed by FDR-adjustment. When performing gene set enrichment testing on the MSigDB Hallmark gene sets, we applied FDR-adjustment to combined *p*-values and defined significant enrichment as gene sets having an adjusted *p*-value < 0.05. When performing gene set enrichment on the four IRE gene sets (***all 3’ IREs***, ***all 5’ IREs***, ***HQ 3’ IREs***, ***HQ 5’ IREs***), we applied Bonferroni-adjustment to combined *p*-values to further protect against Type I errors and defined significant enrichment as gene sets having an adjusted *p*-value < 0.05. Depending on the species in the dataset being analyzed, we used the respective IRE gene sets defined for human, mouse, or zebrafish.

### Analysis of the Caco-2 cultured cell line dataset

We downloaded processed microarray data from GEO (accession number: GSE3573). This study investigated gene expression responses to iron treatments, including iron deficiency (cells in iron-free medium vs. cells cultivated in ferric ammonium nitrate, and iron overload (cells cultivated in DMEM-FBS medium with hemin vs. cells in DMEM-FBS medium) [43]. We performed differential gene expression analysis using the “lmFit” and eBayes” functions in *limma* [40]. Genes were defined as differentially expressed when their FDR-adjusted *p*- value < 0.05.

### Analysis of the Mayo Clinic RNA-seq dataset

No experiments of human subjects were performed in this study as all human data was accessed from public databases and was anonymous. We downloaded processed CPM count data from Synapse (https://www.synapse.org/#!Synapse:syn5550404). We matched cerebellum and temporal cortex samples by their patient ID, and only retained genes which were present across all samples and patients for which there were both cerebellum and temporal cortex samples (n=236 patients with measurements for cerebellum and temporal cortex, 472 samples in total). We performed analysis using *limma* [40, 41] and determined differentially expressed genes between conditions. In addition, we used the “duplicateCorrelation” function in *limma*, setting the “block” parameter to the patient ID. Genes were considered differentially expressed if their FDR-adjusted *p*-value < 0.05.

### Zebrafish husbandry and animal ethics

All zebrafish work was conducted under the auspices of the Animal Ethics Committee (permit numbers S-2017-089 and S-2017-073) and the Institutional Biosafety Committee of the University of Adelaide. Tübingen strain zebrafish were maintained in a recirculated water system.

### fAD-like *psen1*^Q96_K97del/+^ zebrafish

The isolation of the *psen1^Q96_K97del^* mutation has previously been described [44]. Zebrafish mutations were only analyzed in the heterozygous state in this study.

### Acute hypoxia treatment of zebrafish

*psen1*^Q96_K97del/+^ mutants and their wild-type siblings were treated in low oxygen levels by placing zebrafish in oxygen-depleted water for 3 hours (oxygen concentration of 6.6 ± 0.2 mg/L in normoxia and 0.6 ± 0.2 mg/L in hypoxia.

### Whole brain removal from adult zebrafish

After normoxia or hypoxia treatment adult fish were euthanized by sudden immersion in an ice water slurry for at least ∼30 seconds before decapitation and removal of the entire brain for immediate RNA or protein extraction. All fish brains were removed during late morning/noon to minimize any influence of circadian rhythms.

### RNA extraction from whole brain

Total RNA was isolated from heterozygous mutant and WT siblings using the *mir*Vana miRNA isolation kit (Thermo Fisher). RNA isolation was performed according to the manufacturer’s protocol. First a brain was lysed in a denaturing lysis solution. The lysate was then extracted once with acid-phenol:chloroform leaving a semi-pure RNA sample. The sample was then purified further over a glass-fiber filter to yield total RNA. Total RNA was DNase treated using the DNA-free™ Kit from Ambion, Life Technologies according to the manufacturer’s instructions. Total RNA was then sent to the Genomics Facility at the South Australian Health and Medical Research Institute (Adelaide, Australia) to assess RNA quality and for subsequent RNA sequencing *(using poly-A enriched RNA-seq technology, and estimated gene expression from the resulting single-end 75 bp reads using the reference GRCz11 zebrafish assembly transcriptome)*

### Pre-processing of the fAD-like *psen1*^Q96_K97del/+^ zebrafish dataset

RNA-seq libraries contained single-end 75bp Illumina NextSeq reads. We performed quality trimming with *AdapterRemoval* using default parameters. Trimmed reads were pseudo-aligned to the reference zebrafish transcriptome using *Kallisto* (v.0.45) [45] and transcript descriptions from Ensembl release 94. The “catchKallisto” function from *edgeR* [46] was used to import and summarise counts from transcript-level to gene-level, with all subsequent analyses performed at the gene-level.

### Differential gene expression analysis for fAD-like zebrafish dataset

For differential gene expression analysis, we retained all genes with expression of at least 1 cpm in 4 or more samples, and used voomWithQualityWeights to downweight lower quality samples [41]. Contrasts were defined to include all relevant pairwise comparisons between conditions, and genes were considered as differentially expressed using an FDR-adjusted *p*- value < 0.05. The *limma* model used to create the design matrix for this dataset included biological sex as a variable (∼0 + biologicalSex + group, where biologicalSex is either male or female and group refers to the combination of age, hypoxia treatment, and genotype of the samples) to minimize any sex-specific effects in the gene expression analysis. More details regarding the modest effects of biological sex in this dataset are provided in **Supplementary Text 1**.

Estimation of spliced and unspliced gene expression in fAD-like zebrafish dataset For spliced transcripts in Ensembl release 94, we additionally defined unspliced genes including intronic regions. Unspliced transcripts were appended to the end of the reference transcriptome and used to build a new Kallisto [45] index. Estimated counts for spliced transcripts and unspliced genes were imported into R using the “catchKallisto” function from *edgeR* [46].

### Gene set enrichment tests for other zebrafish datasets

Please refer to ref. [44] for details on RNA-seq data processing and analysis of the non-fAD-like dataset, and ref. [47] (GEO accession: GSE148631) for details on RNA-seq data processing and analysis of the 7-day-old Q96_K97del/+ dataset. In the current work, we used the gene expression counts matrix with *limma*. The voom, design, and contrasts objects produced as part of the *limma* analysis were used for gene set enrichment analysis with the zebrafish IRE gene sets we defined as well as the MSigDB Hallmark gene sets. Significantly enriched Hallmark gene sets had FDR-adjusted *p*-value < 0.05 while IRE gene sets were considered significantly enriched if the Bonferroni-adjusted *p*-value < 0.05.

### Differential transcript stability analysis

The estimated spliced and unspliced transcript count estimates from *kallisto* [45] were imported into R using the catchKallisto function from *edgeR* [46]. We used *limma* [40] to determine the logFC of spliced transcripts and unspliced transcripts for each comparison. To test for whether there was a significant difference in the logFC of the spliced and unspliced transcripts, we used Welch’s *t*-test with the *s2*.*prior* values from *limma* as the variances of the spliced and unspliced transcripts. We defined the null (no stabilization of transcript) and alternate (stabilization of transcript) hypotheses for each gene as follows, where s and u refer to the spliced and unspliced versions of a particular gene:

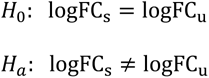

We defined genes with FDR-adjusted *p*-values < 0.05 as having differential stability.

### Checks that observed gene expression differences are not artifacts of differences in proportions of cell types

The Mayo Clinic RNA-seq study, 5XFAD mice, and fAD-like zebrafish datasets are bulk RNA-seq datasets. To confirm that any gene expression changes were likely due to altered transcriptional programs rather than changes in cell type proportions, we compared expression of marker genes for four common neural cell types (astrocytes, neurons, oligodendrocytes, microglia) in conditions within each dataset. The marker genes for astrocytes, neurons, and oligodendrocytes were obtained from MSigDB gene sets from [48] which were based on studies in mice. The marker genes for microglia were derived from [49] which was based on studies in human and mouse. All gene IDs were converted to human, mouse, or zebrafish Ensembl IDs using BioMart [28] for each dataset. Please refer to **Supplementary Text 2** for details.

### Expression of previously validated IRE genes in cell line datasets

Gene symbols from Figure 2 and Supplementary Tables S1 and S2 of Sanchez et al. [50] were matched to corresponding Ensembl gene identifiers using the *biomaRt* R package [28], and filtered to include only genes present in ***HQ 3’ IRE*** and ***HQ 5’ IRE*** gene sets for human and/or mouse. Processed cell line microarray datasets were downloaded from GEO and included a human Caco-2 cell line dataset (GEO accession GSE3573), and mouse splenic B cell dataset (GEO accession GSE77306). The datasets were filtered for validated IRE genes and boxplots were used to visualize expression between groups. *t*-tests were used to test for differences in expression between groups, with *p-*values ≤ 0.05 considered statistically significant. The Sankey plot was produced using the *networkD3* R package [51].

## Results

### Defining sets of genes containing Iron Responsive Elements (IREs)

We utilized the SIREs (Searching for Iron Responsive Elements) tool, to define species-specific IRE gene sets by searching for IRE and IRE-like motifs in the 3’ and 5’ untranslated regions (UTRs) of the reference transcriptomes of human (hg38), mouse (mm10), and zebrafish (z11). *SIREs* assigns a quality-score to all detected IREs, with “high-quality” scores corresponding to canonical IREs and “medium-quality” or “low-quality” scores reflecting deviations from the canonical IRE (alternative nucleotide composition in the apical loop, one bulge at the 3’ or one mismatch in the upper stem) that would still produce an IRE-like motif with potential to be functional [25,52–55]. **Figure 1** summarizes the IRP/IRE interaction effects on transcripts and gives examples of canonical and non-canonical IREs.

**Figure 1.**
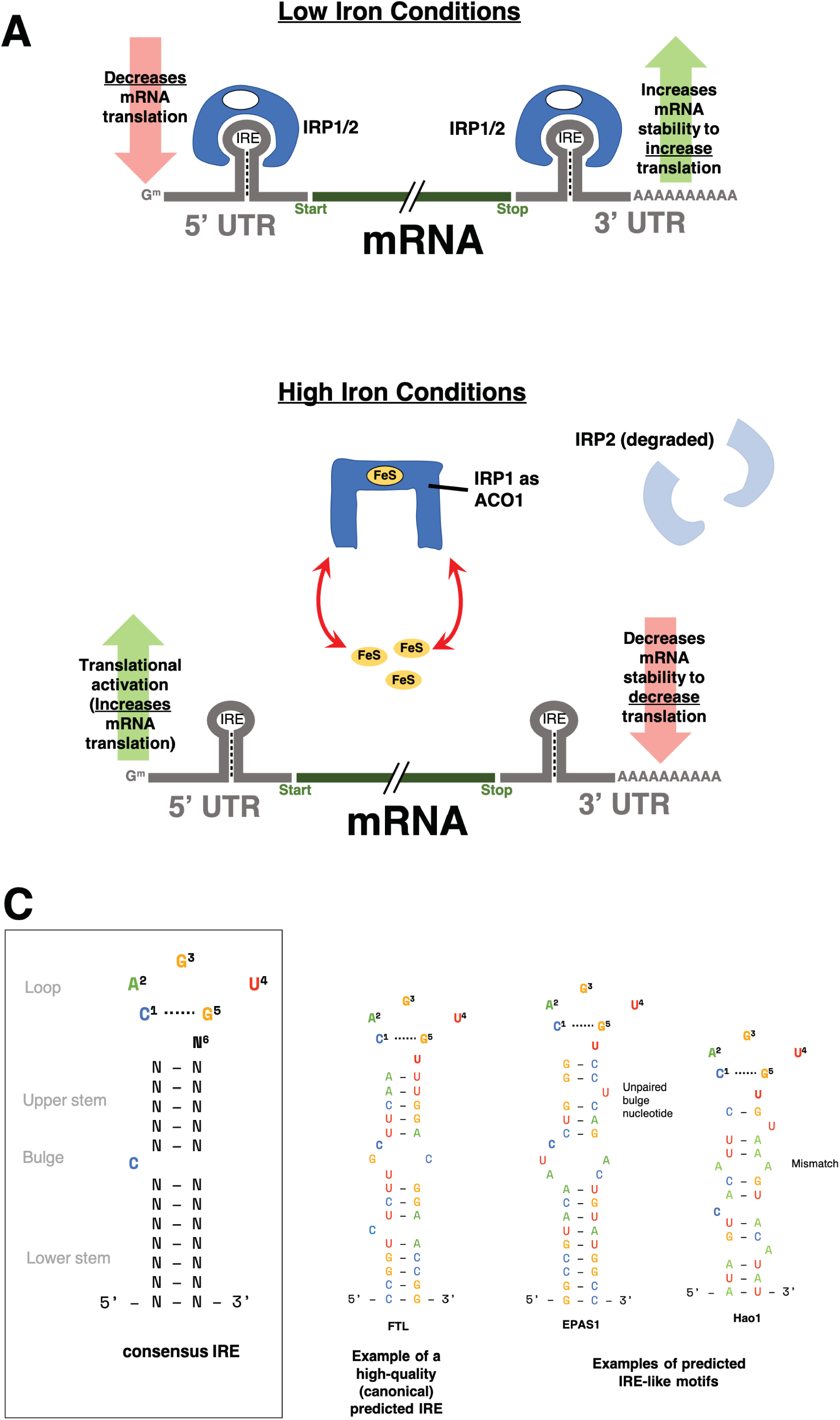
Iron Responsive Element (IRE) background. **A. The classical paradigm of altered transcript stability due to the IRP/IRE system under iron dyshomeostasis.** When cellular iron levels (through FeS) are low, Iron Regulatory Proteins (IRP1 or IRP2) bind IREs in the 5’ or 3’ untranslated regions (UTRs) of genes involved in iron homeostasis. In general, genes with IREs in their 3’ UTRs will be stabilised and hence increased in expression while genes with IREs in their 5’ UTR will have repressed translation from IRP binding. However, when cellular iron levels are high, IRP1 undergoes a conformation change and is now unable to bind IREs. IRP1 now acts as a cytosolic aconitase (ACO1). IRP2 is degraded in high iron conditions and is also unable to bind IREs. Without IRPs binding to IREs, genes with 5’ IREs will have increased translation, while those with 3’ IREs will have decreased mRNA stability, causing their expression to be decreased. **B. Consensus IRE secondary structure and examples of high-quality and IRE-like motifs predicted by SIREs.** IRE-like motifs with non-canonical structure can be detected by SIREs if they have up to one mismatch pair in the upper stem (e.g. *Hao1*) or 1 unpaired bulge nucleotide on the 3’ strand of the upper stem (e.g. *EPAS1*). For more details on the prediction of non-canonical IRE motifs, please refer to Figure 1 of Campillos et al. [25].

We defined four gene sets for each species as follows: ***HQ 3’ IREs*** (high-quality predicted 3’ IRE genes), ***HQ 5’ IREs*** (high-quality predicted 5’ IRE genes), ***all 3’ IREs*** (including all low, medium, and high-quality predicted 3’ IRE genes), and ***all 5’ IREs*** (including all low, medium, and high-quality predicted 5’ IRE genes). The size of these gene sets for each species and the overlap present is shown in **Figure 2** and the gene sets are provided in **Supplementary Table 1**. Overall, searching through human UTRs uncovered the largest number of predicted IRE genes, followed by mouse, and then zebrafish. Overlap between homologous genes from the IRE gene sets of different species was generally poor. The largest sets of genes shared between species were consistently found between human and mouse, which is reflective of their closer evolutionary divergence.

**Figure 2.**
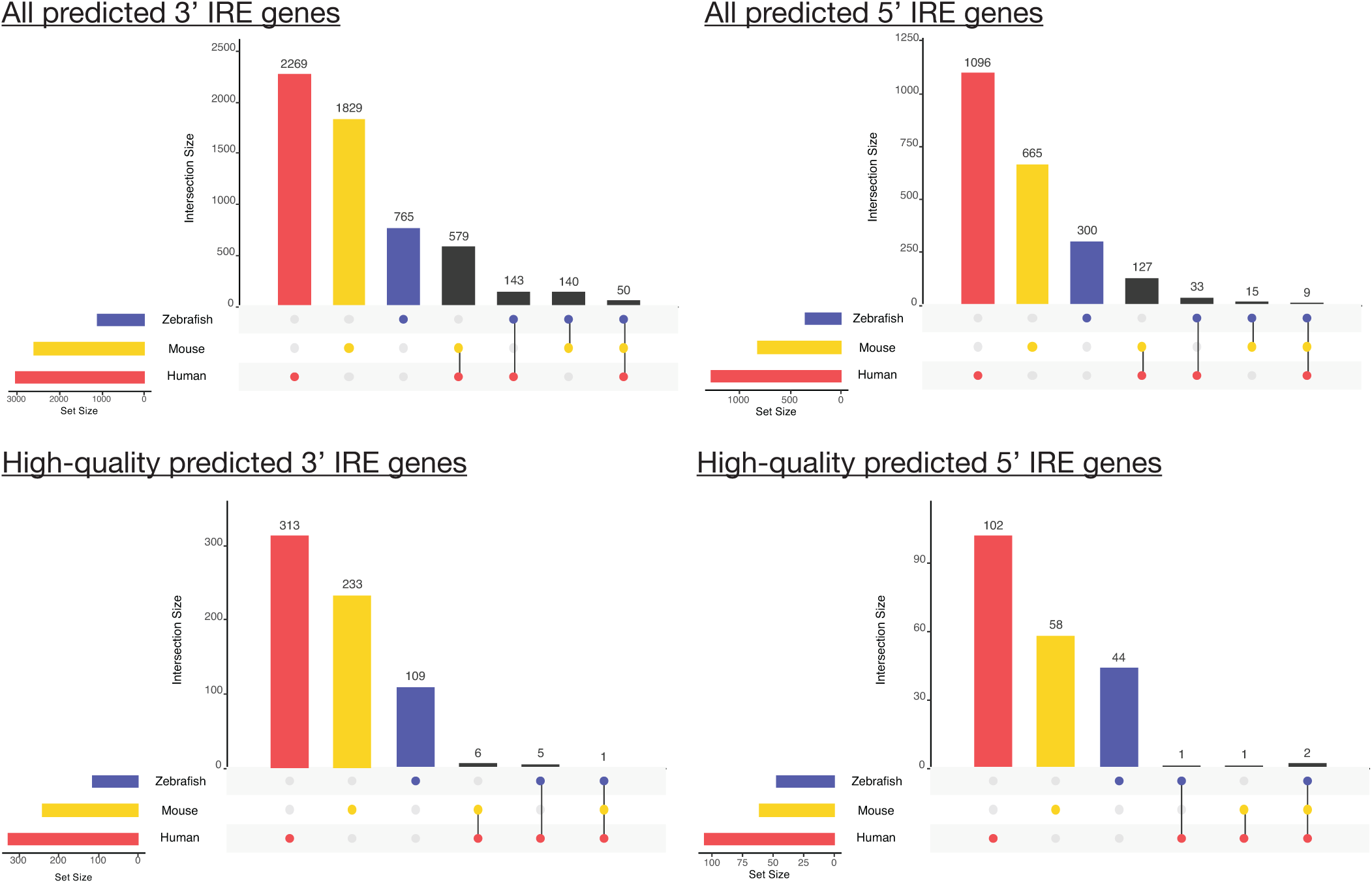
Overlap between predicted IRE gene sets for human, mouse, and zebrafish. The number of genes in the gene set for each species is shown at the bottom-left bars of each UpSet plot, while genes with shared homologs across species are indicated in the main plot region. IRE genes in mouse and zebrafish gene sets were excluded from this plot if they did not have a human homolog.

While not many high-quality IRE genes were identified across all three species, the few we identified are consistent with known and well-characterized IREs in the literature [56, 57]. For example, the single ***HQ 3’ IRE*** gene shared between the three species was *TFRC* (transferrin receptor), while the ***HQ 5’ IRE*** genes shared between the three species included *FTH1* (ferritin heavy chain 1) and *ALAS2* (5’-aminolevulinate synthase 2). We note that the scoring of genes for the ***HQ 3’ IRE*** and ***HQ 5’ IRE*** sets is considered stringent (i.e. only canonical IREs). These two gene sets recover known IRE genes in human, including *CDC14A* (3’ IRE; [50]), *CDC42BPA* (3’ IRE; [23, 58], *SLC40A1* (5’ IRE; [59]), and *FTL* (5’ IRE; [60]). These genes are also recovered in the ***All 3’ IRE*** and ***All 5’ IRE*** sets across mouse and zebrafish too. In addition, the 3’ IRE of *SLC11A2* (also known as *DMT1*) [22] which is only found in mammals, was also recovered in the human ***HQ 5’ IREs*** set and mouse ***All 3’ IREs*** set (but not for zebrafish). The expression of these previously characterized IRE genes across the human, mouse, and zebrafish datasets analyzed in this current work is described in **Supplementary Text 3** and discussed further in the analyses we later describe.

### IRE gene sets are over-represented within up-regulated AD genes, but overall not well-represented in existing gene sets

We explored the biological relevance of the predicted IRE gene sets described above by testing whether genes within them are over-represented in existing gene sets from the Molecular Signatures Database (MSigDB; available at https://www.gsea-msigdb.org/gsea/msigdb). MSigDB represents one of the largest publicly-available collections of gene sets collated from existing studies and other biological databases [29]. We limited our analysis to gene sets from the following collections: Hallmark (non-redundant sets of ∼200 genes each representing various biological activities), C2 (gene sets from databases including KEGG and Reactome and published studies), C3 (gene sets containing genes with motif elements), and C5 (gene sets based on gene ontology terms) (see **Methods**). We performed over-representation analysis for the predicted IRE gene sets (***all 3’ IREs*** and ***all 5’ IREs***) separately for each species (human, mouse, zebrafish), and used Wilkinson’s meta-analytic approach to determine gene sets that were significantly over- represented across the three species’ IRE gene sets. Our results indicated that 1,148 of 10,427 tested gene sets displayed over-representation of IRE gene sets across all three species (Bonferroni-adjusted Wilkinson’s *p*-value < 0.05) (**Figure 3**; **Supplementary Table 2**). The bar plots in **Figure 3** show the 15 MSigDB gene sets with the most significant enrichment for human, mouse, and zebrafish. Remarkably, the “Blalock Alzheimer’s Disease Up” gene set from the C2 collection was significantly over-represented in both the sets of ***all 3’ IREs*** (Bonferroni-adjusted Wilkinson’s *p-*value = 3.9e-14) and ***all 5’ IREs*** (Bonferroni-adjusted Wilkinson’s *p*-value = 2.1e-26) in the meta-analysis, and was also the gene set with the most significant over-representation of the human ***all 3’ IREs*** set (Bonferroni-adjusted Fisher’s exact test *p*-value = 7.8e-58) (**Supplementary Table 2**). This supports that disturbance of iron homeostasis particularly distinguishes Alzheimer’s disease from other disease conditions and pathways represented within the C2 collection. In addition, the top 15 MSigDB gene sets showing the most significant over-representation generally differed between species. However, in all cases, a large proportion of IRE genes from the predicted IRE gene sets were not contained within any of these top-ranked MSigDB gene sets or within the Heme Metabolism gene set belonging to the Hallmark collection (**Figure 3**). This is visualized through the network plots of **Figure 3**, which show that the predicted IRE genes we defined are not fully captured by existing gene sets. This suggests that IRE sets may be uniquely useful for investigating gene expression changes during the IRE-IRP response to iron dyshomeostasis.

**Figure 3.**
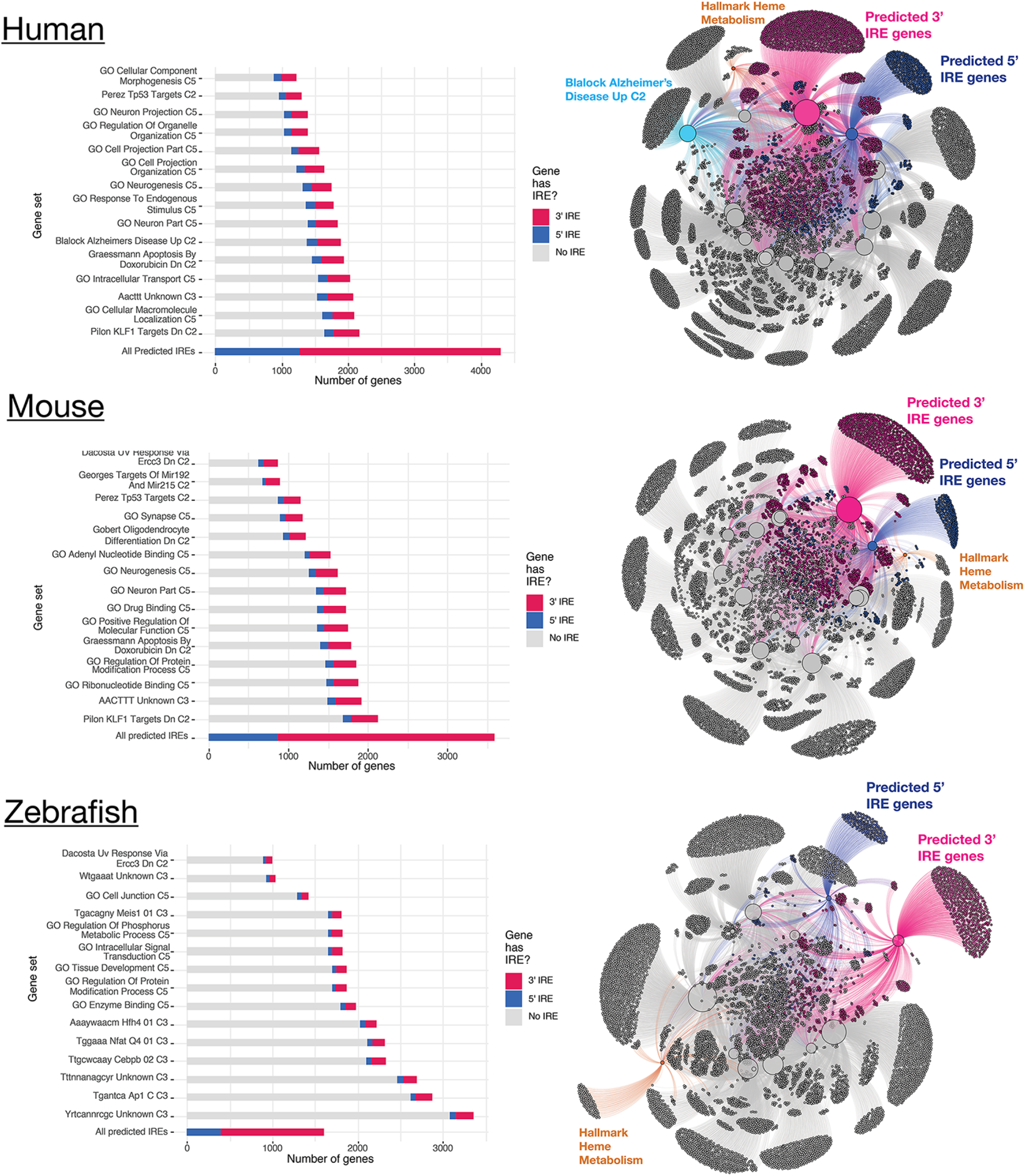
MSigDB gene sets showing over-representation of predicted 3’ and 5’ IRE genes in human, mouse, and zebrafish. The top 15 MSigDB gene sets ranked by Fisher’s exact test *p*-value (testing for over-representation of the *all 3’ IREs* and/or *all 5’ IREs* sets) are shown for each species. In the network plots, the top 15 MSigDB gene sets are shown as large nodes, with genes represented as small nodes. Edges connecting genes to gene sets indicate the gene set(s) that a gene belongs to. Overall, the *all 3’ IREs* and *all 5’ IREs* gene sets have a large proportion of genes which are not included in any of the top ranked MSigDB gene sets for each species.

### Involvement of transcription factor regulation within IRE gene sets

To be confident that IRE gene sets accurately capture information about IRP/IRE-mediated responses, we needed to investigate possible co-ordinate regulation by other factors. As a starting point, we examined whether known binding motifs for transcription factors were significantly over-represented in the promoter regions of genes within each IRE gene set (***all 3’ IREs***, ***all 5’ IREs***, ***HQ 3’ IREs***, and ***HQ 5’ IREs***) for each species. We detected significant over-representation of several transcription factors including the Klf14 motif in the zebrafish ***all 3’ IREs*** set (FDR-adjusted *p*-value = 0.049), the E2F motif in the human ***all 5’ IREs*** set (FDR-adjusted *p*-value = 0.049) and the Bach1 (FDR-adjusted *p*-value = 0.012), Nrf2 (FDR-adjusted *p*-value = 0.013), and NF-E2 (FDR-adjusted *p*-value = 0.019) motifs in the zebrafish ***HQ 5’ IREs*** set (**Supplementary Table 3**). This suggests that the following results should be interpreted with the knowledge that expression of subsets of genes in some of the IRE gene sets may be influenced by factors other than iron ions.

### Gene set enrichment testing approach

Our predicted IRE gene sets can be used in any standard gene set enrichment testing approach to detect potential changes in iron homeostasis between conditions. Our workflow, which we later apply to human, mouse, and zebrafish datasets, is shown in **Figure 4**. Due to variability in the results produced by different gene set enrichment testing methods, we were inspired by the EGSEA framework [61] to combine the results from different methods. Based on an initial analysis using EGSEA, we chose *fry/mroast* [35, 36], *camera* [37], and *fgsea* [38, 39] as the representative methods to use. (See **Supplementary Figure 2** for a principal component analysis of results from different gene set enrichment analysis approaches.) A summary of the different characteristics of the three methods is shown in **Table 1**. The raw enrichment *p-*values from these approaches can be combined to obtain an overall enrichment *p*-value for each gene set. In accordance with EGSEA default parameters, we used Wilkinson’s method to combine raw *p*-values, and then applied adjustment for multiple testing on all combined *p*-values. Along with performing this gene set enrichment testing on our IRE gene sets, we also recommend using the same method to test for enrichment for the MSigDB Hallmark gene sets as the diverse biological activities they represent help to provide context for interpreting the IRE enrichment results. To explore further the results of IRE gene set enrichment analysis, we use UpSet plots to display the overlap between sets of “leading-edge” genes in the ***all 3’ IREs*** and ***all 5’ IREs*** gene sets. The leading-edge genes can be interpreted as the core (often biologically important) genes of a gene set that account for the significant enrichment as calculated by GSEA [39].

**Figure 4.**
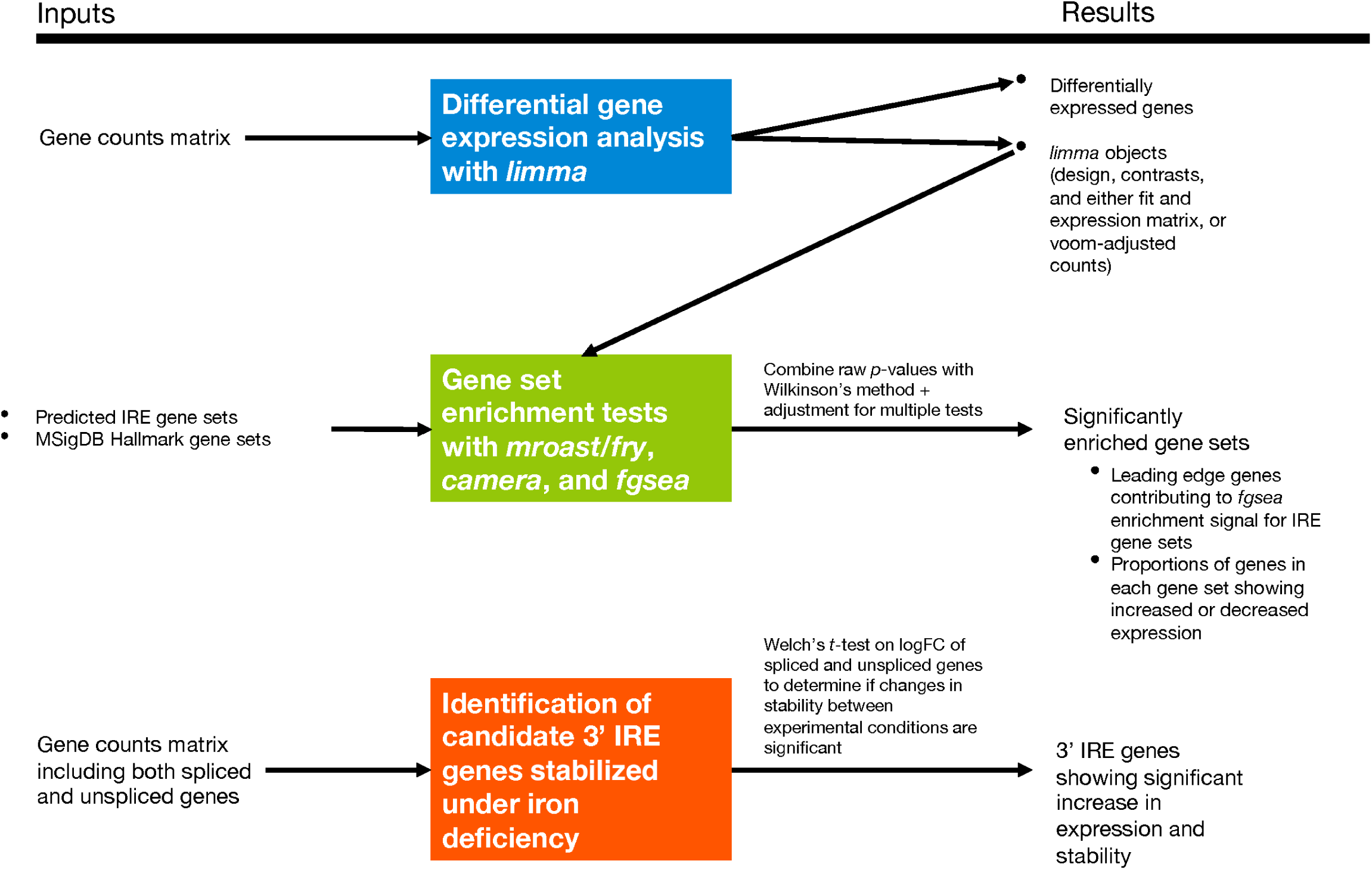
IRE-containing gene expression analysis workflow. The section including identification of candidate 3’ IRE genes stabilized under iron deficiency was only applied to the fAD-like zebrafish dataset due to unavailable raw RNA-seq reads for the other datasets needed to identify expression of unspliced genes.

**Table 1.** Summary of gene set testing approaches used in our analyses.

| Method | Hypothesis | Focused /<br>battery<br>approach | Accounts<br>for inter-<br>gene<br>correlation | R<br>implementation |
| --- | --- | --- | --- | --- |
| **Fry / Roast** | Self-contained null hypothesis (genes within a set have no association with experimental condition) | Focused (each gene set is tested on its own terms, and multiple testing adjustment is applied afterwards) | Yes | “fry” and “mroast” functions in the *limma* Bioconductor package [35,36] |
| **Camera** | Competitive null hypothesis (genes within a set do not have a stronger association with experimental condition compared to a random gene set) | Focused (each gene set is tested on its own terms, and multiple testing adjustment is applied afterwards) | Yes | “camera” function in the *limma* Bioconductor package [37] |
| **fgsea (fast implementation of GSEA in R)** | Self-contained null hypothesis (genes within a set have no association with experimental condition) | Battery (gene sets are pitted against each other to determine which ones are more significantly enriched) | No | “fgsea” function in the *fgsea* Bioconductor package [38,39] |

### Differences in IRE gene set enrichment during iron deficiency and iron overload in a cultured cell line

We first tested how our enrichment approach illuminated the effects of iron overload and deficiency in a cultured cell line microarray dataset from Caco-2 cells (GEO accession: GSE3573). As only one cell type contributed to this dataset, interpretation of the IRE enrichment results is simplified by not having to consider the differing iron requirements of different cell types. This allowed us to focus on whether the iron dyshomeostasis treatments (iron overload and iron deficiency) could be detected and distinguished in terms of their IRP/IRE system-driven transcript abundance response. Because the dataset was from a microarray experiment, we performed only the differential gene expression and gene set enrichment testing portions of our workflow. The results of Principal Component Analysis and differential gene expression analysis are provided in **Supplementary Figure 3**. We note that this particular dataset was chosen despite not being derived from a central nervous system cell type, as maintaining iron homeostasis is fundamental to all cell types (regardless of their tissue of origin). In addition, this particular dataset included iron deficiency and iron overload treatments in the same batch and with sufficient sample number, and no other cultured cell line gene expression dataset was publicly available at the time that fulfilled these criteria.

In general, we found iron deficiency and iron overload treatments resulted in different gene expression responses. In terms of classic IRE genes such as *TFRC, FTL,* and *FTH1*, iron deficiency is associated with increased expression of *TFRC* (consistent with increased transcript stabilization of this 3’ IRE gene) and decreased expression of the 5’ IRE genes *FTL* and *FTH1,* and these changes are in the opposite direction during iron overload (see **Supplementary Text 3** and **Supplementary Figure 1A**). Differential gene expression across all detected genes in the dataset additionally revealed that iron deficiency was associated with 96 differentially expressed genes of which 10 possessed predicted IREs while iron overload was associated with 212 differentially expressed genes (FDR-adjusted *p*- value from *limma* < 0.05) of which 33 possessed predicted IREs (**Supplementary Figure 3B**). There were 17 differentially expressed genes in common between iron deficiency and iron overload treatments, and all moved in opposite directions according to the treatment (i.e. increased abundance under iron deficiency and decreased abundance under iron overload, or vice versa). These clear differences between iron deficiency and iron overload were reflected in gene set enrichment analyses using the MSigDB Hallmark gene sets, where the gene sets involved and the proportions of gene transcripts with increased or decreased abundance differed (**Figure 5A**). As expected, IRE gene sets also showed significant enrichment under iron deficiency and overload conditions (**Figure 5B**) (Bonferroni adjusted *p*-value < 0.05).

**Figure 5.**
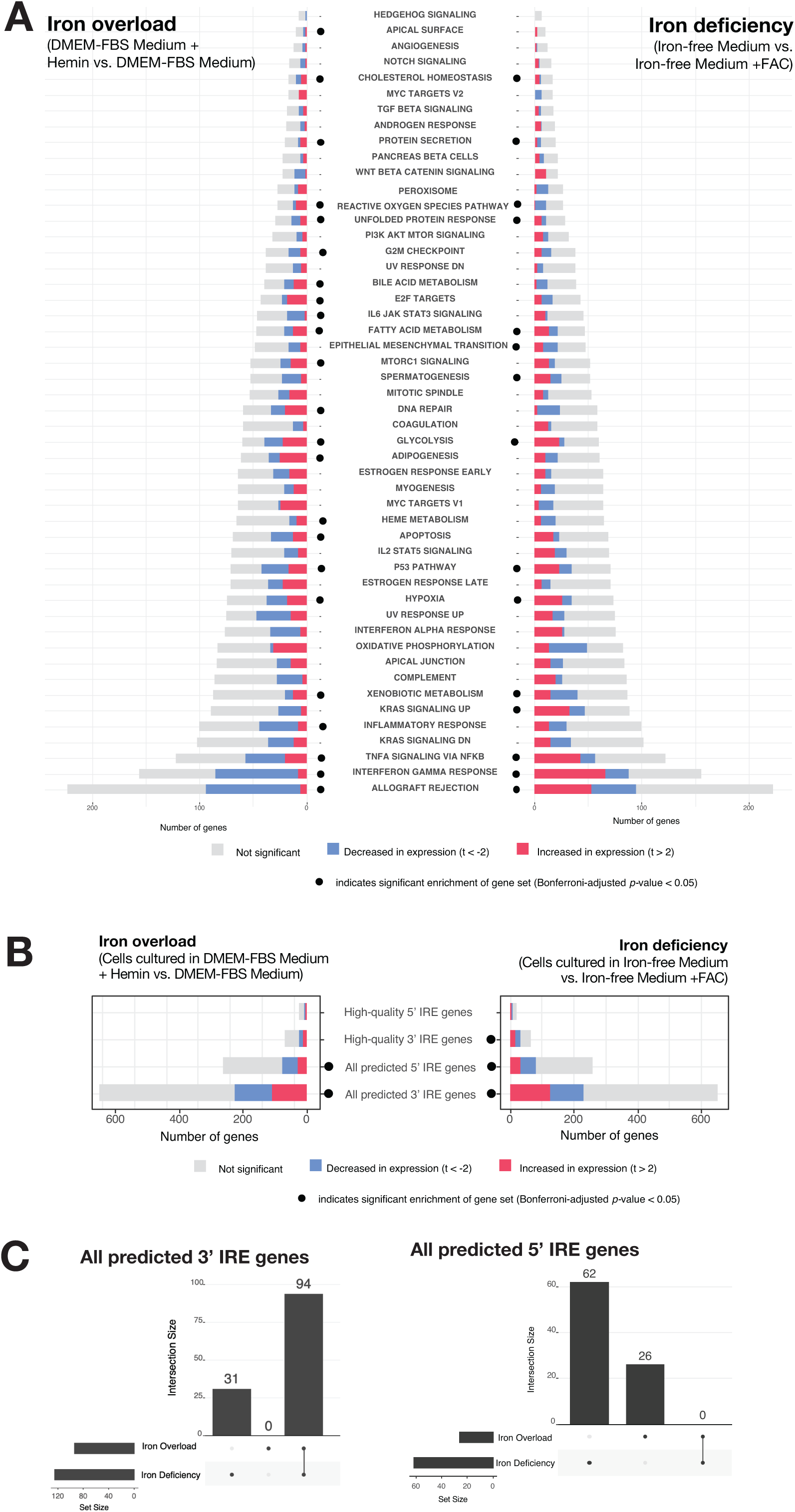
Analysis of Caco-2 cultured cell line microarray dataset. **A.** Gene set enrichment testing results for human predicted IRE gene sets for the iron overload and iron deficiency treatments. Dots indicate if a gene set was considered significantly enriched (Bonferroni-adjusted *p-*value < 0.05) in the iron overload (on left) or iron deficiency (on right) treatment. **B. Gene set enrichment testing results for MSigDB Hallmark gene sets in the iron overload and iron deficiency treatments.** Dots indicate if a gene set was considered significantly enriched (FDR-adjusted *p*-value < 0.05) in the iron overload (on left) or iron deficiency (on right) treatments. **C. UpSet plots showing overlap between iron overload and iron deficiency treatments in GSEA leading-edge genes for the *All 3’ IREs* and *All 5’ IREs* gene sets.** The bars to the lower-left indicate the number of the leading-edge genes for iron overload and iron deficiency treatments, while the bars in the main plot region indicate the number of leading-edge genes which are unique or shared between the treatments.

We expected that iron deficiency would result in increased expression of genes with 3’ IREs under the IRP/IRE paradigm similar to the *TFRC* gene as seen in **Supplementary Text 3**. However, both the 3’ and 5’ IRE gene sets displayed mixed patterns of increased and decreased expression under both the iron deficiency and iron overload treatments (**Figure 5B**). This indicates it would be difficult to distinguish between these conditions based purely on overall increases or decreases in the expression of IRE gene sets. Despite this, we see that the iron deficiency and iron overload treatments can be distinguished by their “leading-edge” genes (those genes contributing most to the enrichment signal for the predicted IRE gene sets (***all 3’ IREs*** and ***all 5’ IREs***) (**Figure 5C**). This supports that gene set enrichment using predicted IRE gene sets may be sufficiently sensitive to detect whether iron dyshomeostasis is present and potentially more informative than individual IRE genes in distinguishing between different IRP-IRE system-mediated gene expression responses in iron deficiency and iron overload treatments in this cell line.

### A distinct iron homeostasis response in human AD patients compared to other neuropathologies

Given that IRE gene sets could distinguish between iron overload and deficiency in a cultured cell line, we next tested our gene sets on a more complex dataset including cerebellum and temporal cortex tissue samples from post-mortem human brains. The brains originated from either healthy controls or patients afflicted with one of three conditions: Alzheimer’s disease (AD); pathological aging (PA), a condition involving amyloid pathology but no significant dementia symptoms; or progressive supranuclear palsy (PSP), a tauopathy without amyloid pathology [62]. An important characteristic of the dataset is that both cerebellum and temporal cortex tissue samples were available from each patient. A summary of the 236 patients whose data we included in this analysis is shown in **Table 2** and the results of differential gene expression analysis and IRE gene set enrichment analysis are shown in **Table 3** and **Figure 6A**, and summarised in comparison to the mouse and zebrafish datasets later analysed in **Table 5**. The expression of classic IRE genes including *TFRC*, *FTL,* and *FTH1* in this dataset is shown in **Supplementary Text 3** and **Supplementary Figure 1B-C**. In our analyses, we focus mainly on comparing conditions within each tissue rather than between tissues. This is because we found significant differences in the AD vs. control comparison in the temporal cortex compared to the cerebellum (see **Supplementary Figure 4**).

**Figure 6.**
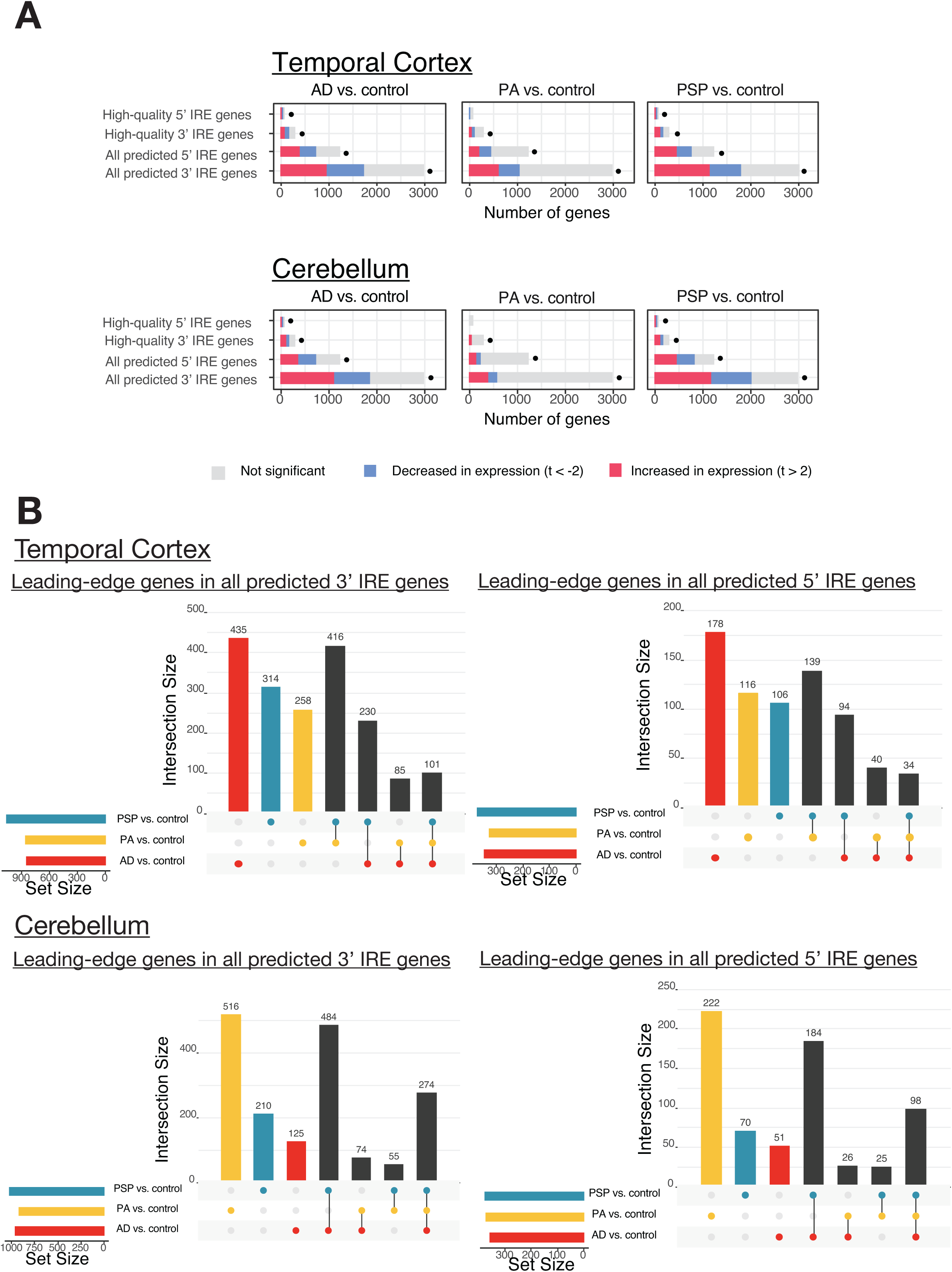
Analysis of Mayo Clinic RNA-seq dataset with human IRE gene sets. **A. Enrichment analysis results of IRE gene sets in comparisons of AD (Alzheimer’s Disease), PA (Pathological Ageing) and PSP (Progressive Supranuclear Palsy) vs. control.** The proportions of IRE genes within each IRE gene set with increased, decreased, or unchanged expression are indicated as coloured bars. The dots indicate that the IRE gene set was significantly enriched (Bonferroni-adjusted enrichment *p*-value < 0.05). **B. UpSet plots showing overlap between leading-edge IRE genes in comparisons of AD (Alzheimer’s Disease), PA (Pathological Aging), and PSP (Progressive Supranuclear Palsy) vs. control.** The bars to the lower-left region of each UpSet plot indicate the number of leading-edge IRE genes for each comparison, while bars in the main plot region indicate the number of leading-edge IRE genes which are unique or shared between comparisons.

**Table 2.** Samples analyzed from Mayo Clinic RNA-seq study.

| Group | Total number | % Female | Mean age ± s.d (years) | Diagnosis |
| --- | --- | --- | --- | --- |
| Alzheimer's disease (AD) | 134 | 56.7 | 82.6 ± 7.3 | Braak Stage ≥ 4, diagnosis according to NINCDS-ADRDA criteria |
| Control | 130 | 46.2 | 82.9 ± 8.3 | Braak Stage ≤ 3, No or sparse CERAD neuritic and cortical plaque density, no diagnoses for any neurodegenerative disease |
| Pathological Aging (PA) | 44 | 54.5 | 84.5 ± 4.3 | Braak Stage ≤ 3, CERAD neuritic and cortical plaque density of 2 or more, no diagnoses for any neurodegenerative disease or mild cognitive impairment |
| Progressive supranuclear palsy (PSP) | 164 | 40.2 | 73.8 ± 6.5 | Braak Stage ≤ 3, no or sparse CERAD neuritic and cortical plaque density |

**Table 3.** Differential gene expression and IRE gene set enrichment results from Mayo Clinic RNA-seq study. Asterisks (*) indicate significant enrichment (Bonferroni-adjusted Wilkinson’s *p*-value from *fry*, *camera*, and *fgsea* < 0.05).

|  | Comparison | No. of DE genes (FDR *p*-value < 0.05 and abs(logFC) > 0.5) | *All 3' IREs* enrichment *p*-value | *All 5' IREs* enrichment *p*-value | *HQ 3' IREs* enrichment *p*-value | *HQ 5' IREs* enrichment *p*-value |
| --- | --- | --- | --- | --- | --- | --- |
| Effect of Alzheimer's disease | AD vs. control (cerebellum) | Down: 1,422<br>Up: 1,308 | 0.0109* | 0.128 | 0.770 | 0.770 |
|  | AD vs. control (temporal cortex) | Down: 1,650,<br>Up: 1,929 | 9.03E-15* | 2.44E-14* | 4.50E-15* | 8.95E-08* |
| Effect of pathological aging (amyloid pathology, no dementia symptoms) | PA vs. control (cerebellum) | Down: 254, Up: 463 | 0.000116* | 0.00139* | 0.0201* | 0.770 |
|  | PA vs. control (temporal cortex) | Down: 466, Up: 512 | 0.000521* | 0.0461* | 0.000065* | 0.0000682* |
| Effect of progressive supranuclear palsy (PSP) | PSP vs. control (cerebellum) | Down: 2,550,<br>Up: 1,080 | 1.81E-22* | 3.45E-22* | 1.43E-12* | 0.0399* |
|  | PSP vs. control (temporal cortex) | Down: 1,271,<br>Up: 622 | 3.58E-11* | 5.17E-13* | 1.39E-06* | 0.0399* |
| Differences between Alzheimer's disease and pathological aging | AD vs. PA (cerebellum) | Down: 1,728,<br>Up: 1,006 | 0.000116* | 0.00139* | 0.0201* | 0.770 |
|  | AD vs. PA (temporal cortex) | Down: 2,633,<br>Up: 2,174 | 0.000521* | 0.0461* | 0.000065* | 0.0000682* |
| Differences between Alzheimer's disease and progressive supranuclear palsy | AD vs. PSP (cerebellum) | Down: 108, Up: 659 | 0.0019* | 0.00184* | 0.158 | 0.158 |
|  | AD vs. PSP (temporal cortex) | Down: 2,052,<br>Up: 2,354 | 9.12E-22* | 2.10E-22* | 5.35E-16* | 0.040* |
| Differences between cerebellum and temporal cortex tissue | Cerebellum vs. temporal cortex in controls | Down: 10,925,<br>Up: 9,302 | 0.00E+00* | 0.00E+00* | 2.82E-269* | 1.60E-81* |
|  | Cerebellum vs. temporal cortex in AD | Down: 11,966,<br>Up: 9,950 | 0.00E+00* | 0.00E+00* | 7.71E-302* | 9.35E-92* |
|  | Cerebellum vs. temporal cortex in PA | Down: 10,200,<br>Up: 8,737 | 1.87E-153* | 5.88E-160* | 1.22E-89* | 1.31E-88* |
|  | Cerebellum vs. temporal cortex in PSP | Down: 11,552,<br>Up: 9,576 | 0.00E+00* | 0.00E+00* | 0.00E+00* | 4.74E-104* |

Overall, our IRE enrichment analyses indicate significant enrichment of IRE gene sets in all pathological conditions (AD, PA or PSP) compared to healthy controls within the cerebellum and temporal cortex (**Figure 6A**). In all pathological conditions, 3’ and 5’ IRE gene transcripts show overall mixed patterns of abundance (e.g. increased and decreased). Further examination of the leading-edge genes from these IRE gene sets gives more insight regarding potential differences and similarities between AD, PA, and PSP (**Figure 6B**). Overall, AD, PSP, and PA appear to involve distinct yet partially overlapping IRE gene expression responses in a tissue-specific manner. Within the temporal cortex, there are 435 3’ IRE leading-edge genes and 178 5’ IRE leading-edge genes exclusively present in the “AD vs. control” comparison. These are greater numbers than for the other two conditions. Interestingly, AD and PA share only relatively small numbers of leading-edge genes despite the fact that many regard PA as a prodrome of AD [63]. Also interesting is that, in the cerebellum, while AD and PSP share many 3’ and 5’ IRE leading-edge genes, PA is associated with a large number of unique leading-edge 3’ and 5’ IRE genes, further emphasizing its difference from AD and also PSP. These observations are supported when considering expression of previously characterized IRE genes individually (see **Supplementary Text 3** and **Supplementary Figure 1B-C**). The results in this Supplementary Text suggest that, in particular, some 3’ IRE genes (e.g. *TFRC*, *CAV3*, *CDC14A*, *CDC42BPA, SERTAD2, TRAM1*) show increased expression in AD (relative to controls) whereas PA and PSP do not. Overall, our results suggest that IRP-IRE system-mediated gene expression changes may be sufficiently sensitive to discern and identify potential differences in the way iron homeostasis is regulated in AD compared to PA and PSP.

### Age-dependent disruption of IRE-driven iron homeostasis in the 5XFAD mouse model

Alzheimer’s disease is thought to develop over decades [64–66]. However, detailed molecular studies of AD brains can only be based on post-mortem tissues. To reveal the early molecular changes that initiate the progression to AD we must study the brains of animal disease models. Given that the IRE gene sets appear to work well in human datasets, we then tested our mouse IRE gene sets on an RNA-seq dataset derived from brain cortex tissue of the 5XFAD transgenic mouse AD model (GEO: GSE140286). The 5XFAD mouse is one of the most common systems used to model the amyloid beta histopathology of AD brains and, in a concordance analysis, brain gene expression patterns in this model were previously shown to resemble those of human AD brains to a greater degree than for four other mouse models (3xTg, CK-p25, J20, and Tg2576) [67]. 5XFAD mice possess two transgenes that include a total of five different mutations, each of which individually causes fAD in humans. In this dataset, the mice analyzed were either 3, 6 or 12 months of age. We acknowledge that the small sample size of this particular dataset means that results should be interpreted tentatively (see Discussion).

Using gene set enrichment testing methods as before, we observed significant enrichment of the ***all 3’ IREs*** and ***all 5’ IREs*** gene sets in several comparisons. These included 5XFAD vs. wild type comparisons and wild type aging comparisons (Bonferroni-adjusted enrichment *p*- value < 0.05) (**Figure 7**). Notably, even the youngest age group of 5XFAD mutant mice (3 months) displayed significant enrichment of genes containing 3’ or 5’ IREs compared to age-matched wild types. This is consistent with an enrichment of immune-related Hallmark gene sets that we observed in this age group (see **Figure 7B**) and with a previous transcriptome analysis suggesting immune activation in 5XFAD mice as early as 2-4 months [68]. UpSet plots of overlapping leading-edge genes suggest that the IRE responses due to aging and due to the 5XFAD transgenes may involve partially overlapping gene expression changes (**Figure 8**). In addition, the UpSet plots reveal subsets containing large numbers of IRE-containing genes which uniquely contribute to enrichment of IRE gene sets in only one comparison (e.g. 256 3’ IRE genes only contribute to enrichment in the “*5xFAD vs WT at 6 months*” comparison). Notably, there are 126 shared 3’ IRE genes contributing to enrichment in the “*5xFAD vs WT at 6 months*” and “*5xFAD vs WT at 3 months*” comparisons, but no genes shared between these comparisons and the “*5xFAD vs WT at 12 months*” comparison. These 126 genes were found to show over-representation of Gene Ontology terms including “Hsp70 protein binding” and “nuclear import signal receptor” (FDR-adj. *p*- value < 0.05) (**Supplementary Table 4**), suggesting that subsets of leading-edge genes may have biological relevance, and that these may give insight into the biologically distinct age-dependent iron dyshomeostasis responses caused by the 5XFAD transgenes. Although beyond the scope of our current analysis, further investigation into these subsets of genes may contribute to definition of differences in iron dyshomeostasis responses under different biological conditions of interest.

**Figure 7.**
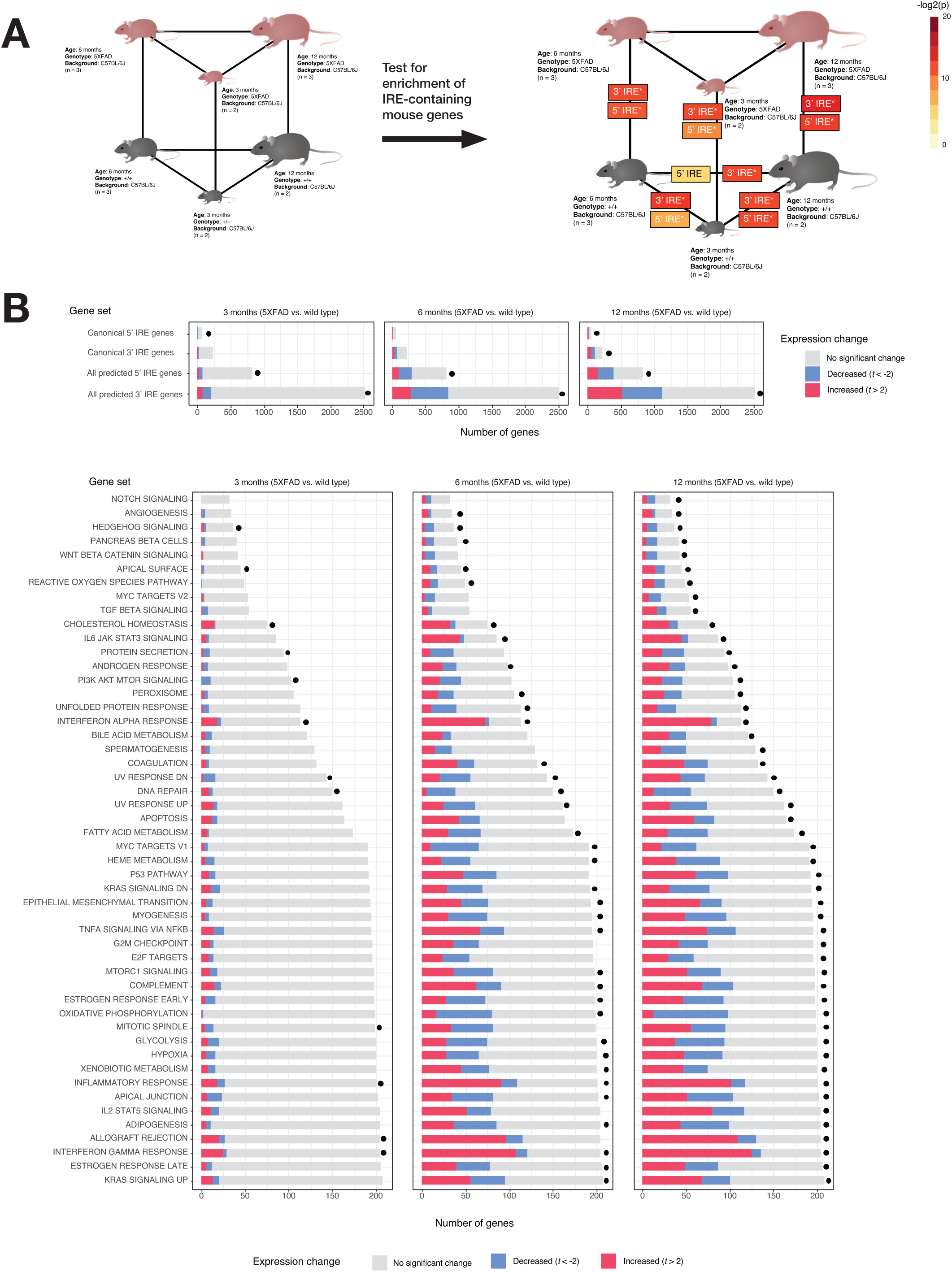
Analysis of 5XFAD mouse dataset. **A. Experimental design and results of IRE gene set enrichment analysis.** The gene sets *all 3’ IREs* and *all 5’ IREs* derived from searching for IRE sequences in the UTRs of genes in the reference mouse genome mm10 are represented here as “3’ IRE” and “5’ IRE” respectively. Asterisks (*) indicate that the gene set was significantly enriched in a particular comparison (Bonferroni-adjusted *p*-value < 0.05). **B. Proportions of genes in IRE and MSigDB Hallmark gene sets which are increased (t > 2) or decreased (t < -2) in expression in all “5XFAD vs. wild type” comparisons**. A dot next to a bar indicates that the gene set was significantly enriched (FDR-adjusted *p*-value < 0.05).

**Figure 8.**
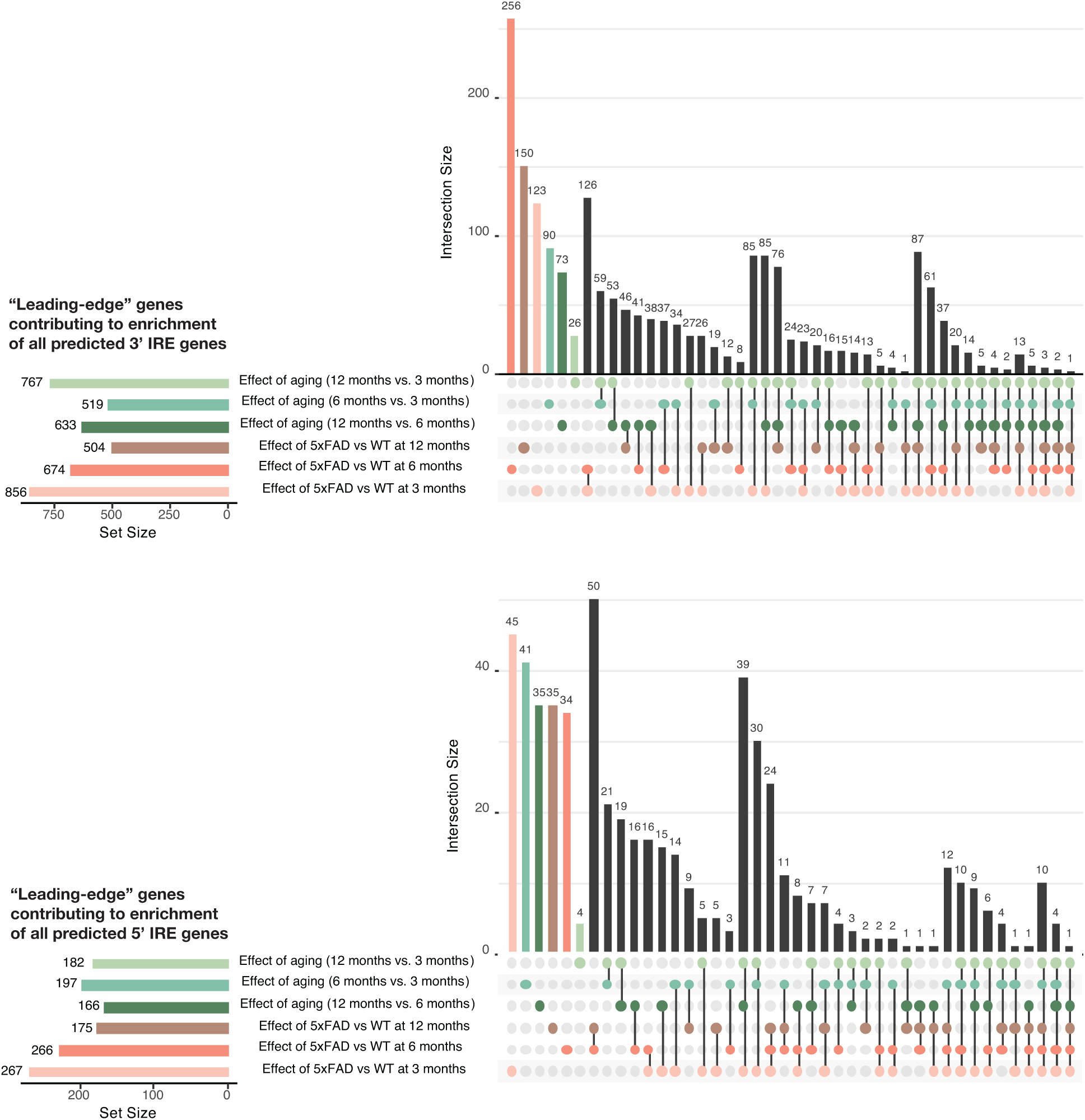
UpSet plots showing the overlap in GSEA leading-edge genes between all comparisons in the 5XFAD mouse datasets for the gene sets all 3’ IREs and all 5’ IREs. Numbers of genes for each intersection are shown above intersection bars.

### Similarities in IRE-driven iron homeostasis responses during hypoxia and from a familial AD-like mutation in a zebrafish model

Concerns have been raised over the relevance of transgenic mouse models in modelling the early stages of AD (reviewed by [69]). Meanwhile, knock-in models of familial AD (fAD) (that possess a single, heterozygous mutation in an endogenous gene like the human version of the disease) only exhibit subtle pathological changes with little to no visible amyloid or tau pathology present [70]. Despite the comparatively more subtle pathology seen in heterozygous, single fAD-like mutation knock-in models, since they closely mimic the genetic state of fAD, analysis of their brain gene expressionthem may reveal molecular changes relevant to early fAD states in human brains. We had access to a whole-brain RNA-seq dataset from a knock-in zebrafish model of fAD possessing a heterozygous, single fAD-like mutation in its endogenous *psen1* gene (*psen1*^Q96_K97del^/+) (henceforth referred to as the “fAD-mutation-like” model in this paper). Previous analysis of a subset of this dataset involving young adult (6-month-old) brains revealed gene expression changes related to energy metabolism and mitochondrial function, especially ATP synthesis and other ATP-dependent functions including lysosomal acidification [44]. (In contrast, the 3-month-old young adult 5XFAD mouse brain dataset is dominated by immune/inflammation responses, **Figure 7B**). Considering the critical role of iron homeostasis in energy metabolism, we decided to revisit this zebrafish dataset. We performed IRE gene set enrichment on the entire dataset in order to include exploration of the effects of aging and acute hypoxia (two important risk factors for sporadic late onset AD) and to analyze how these effects interact with the fAD-like *psen1^Q96_K97del^*/+ mutant genotype. The experimental design and the results of the differential gene expression analyses and gene set enrichment tests are summarized in **Figure 9**.

**Figure 9.**
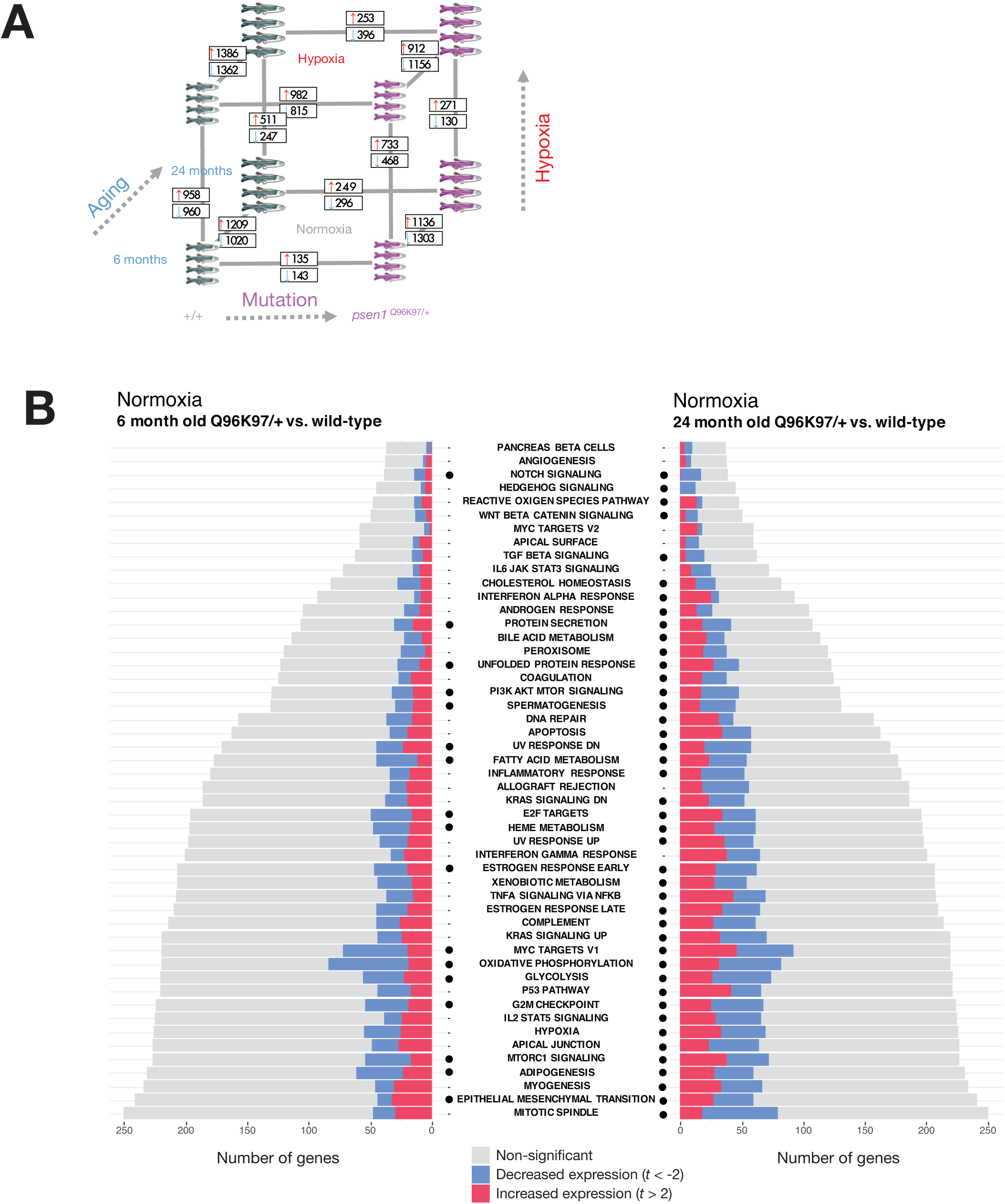
Differential gene expression and gene set enrichment analysis in the fAD-like zebrafish dataset. **A. Results of differential gene expression analysis. Genes which were significantly increased or decreased in expression are indicated in boxes.** These differentially expressed genes have FDR-adjusted *p*-value < 0.05. **B. Gene set enrichment with MSigDB Hallmark gene sets.** The comparisons between Q96_ K97del/+ fAD-like mutants and their wild-type siblings are shown for the 6-month-old (young adult) and 24-month-old (infertile adult) age groups. Dots indicate gene sets which are significantly enriched (FDR-adjusted *p*-value < 0.05).

We first turned our attention to the comparison between young adult (6-month-old) *psen1^Q96_K97del^*/+ mutant zebrafish and their wild-type siblings. At this age, gene expression changes in the mutant fish likely represent early stresses driving the development of fAD in humans. Gene set enrichment tests in this comparison identify alteration of energy metabolism-related Hallmark gene sets (e.g. OXIDATIVE PHOSPHORYLATION, GLYCOLYSIS) (**Figure 9B**) which is consistent with a previous analysis of gene ontology terms with this dataset [44]. In addition, we see enrichment of other gene sets including FATTY ACID METABOLISM, PI3K AKT MTOR SIGNALLING, MTORC1 SIGNALLING, and HEME METABOLISM. All of these gene sets showing enrichment in 6-month-old *psen1^Q96_K97del^*/+ mutants relative to wild-type siblings are also enriched in 24-month-old (“middle-aged”) *psen1^Q96_K97del^*/+ mutants relative to their wild-type siblings (**Figure 9B**). This supports that the biological activities represented in these gene sets are amongst the earliest altered in this particular fAD mutation model.

In Lumsden et al. [14] we predicted that fAD mutations in the major locus *PSEN1* might cause an early cellular deficiency of ferrous iron due to the observation of insufficient acidification of the endolysosomal pathway in in-vitro *PSEN1* mutation studies [71, 72]. In Newman et al. [44] we saw that Gene Ontology enrichment analysis of 6-month-old *psen1^Q96_K97del^*/+ zebrafish brains supported that lysosomal acidification was affected. Therefore, we applied our IRE enrichment analysis to test for evidence of iron dyshomeostasis in these fish. The enrichment of the ***all 3’ IREs*** gene set in 6-month-old *psen1^Q96_K97del^*/+ zebrafish brains supports that iron dyshomeostasis is an important stress in the early stages of fAD (**Figure 10A**).

**Figure 10.**
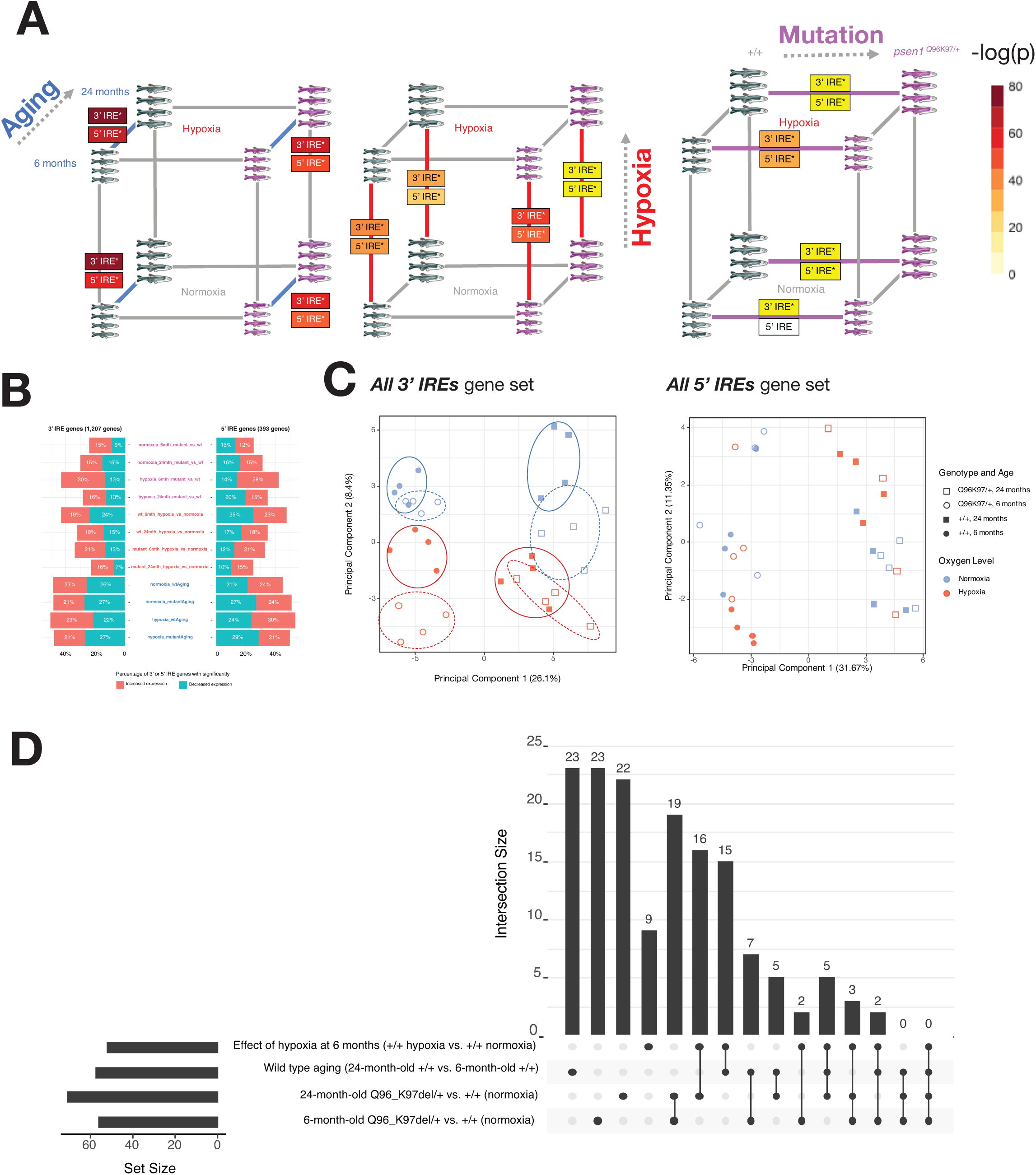
Iron Responsive Element (IRE)-containing gene expression in the fAD-like zebrafish dataset. **A. Results of gene set enrichment testing using predicted IRE gene sets.** We represent the gene sets all 3’ IREs and all 5’ IREs derived from searching for IRE and IRE-like sequences from z11 reference zebrafish gene UTRs as “3’ IRE” and “5’ IRE” in the panel. **B. Proportions of predicted IRE genes which are increased (t > 2) or decreased (t < -2) in expression for each pairwise comparison in the dataset. C. Principal component analysis of all genes in the sets all 3’ IREs and all 5’ IREs for all samples.** Circles on the 3’ IRE plot show that different conditions generally have distinct expression of genes in the *all 3’ IREs set* but not in the *all 5’ IREs* set. **D. UpSet plot showing overlap in leading-edge genes for the *all 3’ IREs* gene set for select comparisons.**

While almost all pairwise comparisons in the zebrafish dataset show significant enrichment of at least one IRE gene set (**Figure 10A**), the expression of IRE genes appears to differ in terms of the proportions of IRE-containing transcripts which show increased versus decreased abundance (**Figure 10B**). In addition, the Principal Component Analysis plot of expression of the ***all 3’ IREs*** gene set over all samples shown in **Figure 10C** suggests that different conditions appear to drive distinct patterns of expression of these genes. Across the first principal component, different age groups (6- and 24-month-old brains) differ in their expression of predicted 3’ IRE genes, while the second principal component appears to separate *psen1*^Q96_K97del^/+ mutants from their wild-type siblings. This separation between *psen1^Q96_K97del^*/+ mutants and wild type siblings is even more pronounced when both are exposed to hypoxia.

To gain more insight into the similarities and differences between the IRE responses in the *psen1^Q96_K97del^/+* vs. wild type comparison, we plotted UpSet plots of overlapping leading- edge genes (**Figure 10D**). These plots suggest that the IRE responses during hypoxia, during aging, and due to this fAD-like mutation, are mostly distinct from each other with unique leading-edge genes. However, similarities in the IRE responses between different conditions are suggested by the existence of some shared leading-edge genes. For example, for the set of ***all 3’ IREs***, the “*6-month-old psen1^Q96_K97del^/+ vs. wild type*” comparison shares 19 leading-edge genes with the “*24-month-old psen1^Q96_K97del^/+ vs. wild type*” comparison. As an initial exploration of the biological relevance of these genes, we submitted them to the STRINGR tool. This indicated that the proteins products of these genes were significantly associated with each other (e.g. in terms of text-mining from PubMed articles, experimentally determined interactions, and co-expression). These proteins were significantly over-represented in the gene sets “MAPK pathway”, “AP-1 transcription factor”, “Jun-like transcription factor”, and “signaling by TGF-beta family members” (FDR-adjusted over-representation p-value < 0.05; **Supplementary Figure 5**). The AP-1 and MAPK pathways have previously been shown to be stimulated by iron depletion [73, 74]. Therefore, mechanistically, it is possible that the IRE-IRP response to iron dyshomeostasis in the fAD-like zebrafish mutant involves these pathways.

The fact that aging, hypoxia, and a fAD-like mutation all cause changes in the abundance of IRE-containing transcripts in zebrafish raised concerns regarding the specificity of such changes. Therefore, as a negative control, we examined changes in IRE transcript abundance in a brain transcriptome dataset derived from zebrafish heterozygous for a *psen1* mutation now thought not to be fAD-like, *psen1^K97fs^* [75]. In this dataset, *psen1^K97fs^*/+ zebrafish are compared to their wild type siblings at 6 and 24 months. We tested this dataset for enrichment using our zebrafish 3’ and 5’ IRE gene sets but found no significant enrichment of any of our predicted IRE gene sets in *psen1^K97fs^*/+ vs. wild type comparisons at any age (**Table 4**; **Supplementary Figure 6**). Reassuringly, we still observed significant enrichment of both 3’ and 5’ IRE gene sets during wild type aging (*24-month-old* wild types *vs. 6-month-old* wild types), consistent with the equivalent comparison in the *psen1*^Q96_K97del^/+ dataset. These results support that IRE-containing transcript abundance changes are sufficiently sensitive to reflect differences in iron homeostasis between different mutation models.

**Table 4.**
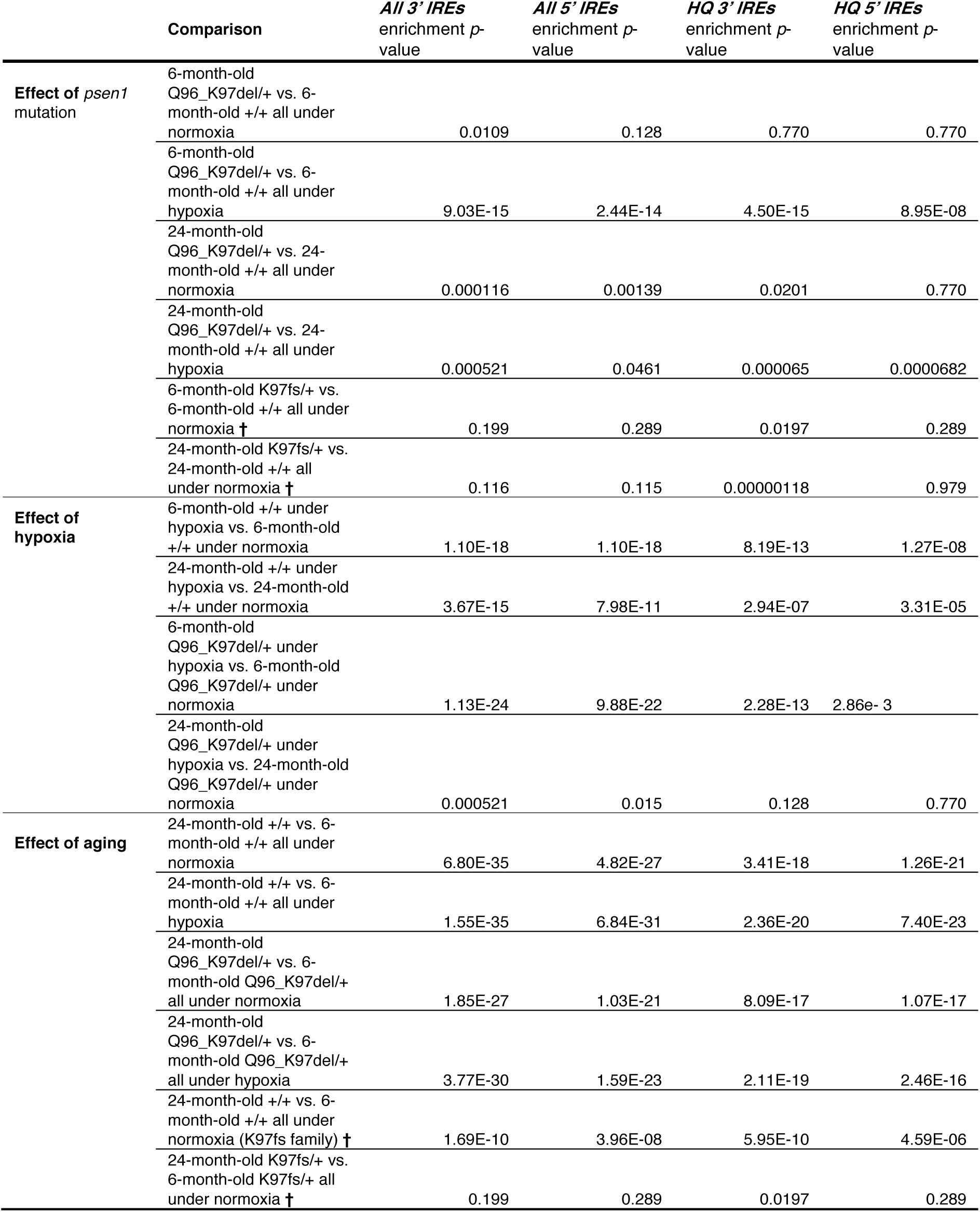
Enrichment of Iron Responsive Element (IRE) gene sets in fAD-like zebrafish dataset. Raw *p*-values from *fry*, *camera* and *fgsea* were combined with Wilkinson’s method, with combined *p*-values then Bonferroni-adjusted for multiple testing. The same process was repeated for the K97fs/+ dataset, which involves an independent family of fish (†).

**Table 5.** Summary of enriched gene sets from the MSigDB Hallmark Gene Set Collection in Datasets Analyzed. Gene sets shown here are representative enriched gene sets from the MSigDB Hallmark Collection (Bonferroni-adjusted Wilkinson’s *p*-value from *fry*, *camera*, and *fgsea* < 0.05). For full lists of enriched gene sets, see relevant figures or Supplemental Tables). For gene sets which had a significant direction, an up (↑) or down (↓) arrow is used to indicate whether most of the genes in the gene set are increased or decreased in expression. No arrow indicates gene sets involving both increased and decreased expression from different subsets of genes.

| Dataset | Comparison | Hallmark Gene Sets Enriched* |
| --- | --- | --- |
| Caco-2 cell line<br>(see Fig. 5) | Iron Overload | ↓ Interferon Gamma Response<br>↑ Reactive Oxygen Species Pathway |
|  | Iron Deficiency | ↑ Interferon Gamma Response<br>↓ Reactive Oxygen Species Pathway |
| Human Mayo-Clinic RNA-seq<br>Study (see Supplementary Table 7) | AD vs. control (Temporal Cortex) | ↓ Oxidative Phosphorylation<br>↑ TNFA signalling via NFκB<br>↑ Interferon Alpha Response<br>↑ Inflammatory Response<br>↑ Interferon Gamma Response |
|  | AD vs. control (Cerebellum) | ↑ G2M Checkpoint<br>↑ Mitotic Spindle<br>↑ PI3K AKT MTOR Signalling<br>↑ Hypoxia |
|  | PA vs. control (Temporal Cortex) | Estrogen Response<br>Heme Metabolism<br>Complement<br>Adipogenesis<br>Inflammation |
|  | PA vs. control (Cerebellum) | Estrogen Response<br>Glycolysis<br>Hypoxia<br>Unfolded Protein Response |
|  | PSP vs. control (Temporal Cortex) | ↓ Interferon Alpha Response<br>↑ Reactive Oxygen Species Signalling Pathway<br>↑ Glycolysis<br>↑ Hypoxia<br>↑ Estrogen Response |
|  | PSP vs. control (Cerebellum) | ↑ Myc Targets<br>↑ G2M Checkpoint<br>↑ Mitotic Spindle<br>↑ Heme Metabolism<br>↑ Hypoxia<br>↓ Fatty Acid Metabolism |
| Mouse 5XFAD model<br>(see Fig. 7) | 5XFAD vs. WT (3 months old) | ↑ Interferon Gamma Response<br>↑ Interferon Alpha Response<br>↑ Cholesterol Homeostasis |
|  | 5XFAD vs. WT (6 months old) | Above also enriched + many other gene sets, see Fig. 7 for more details) |
|  | 5XFAD vs. WT (12 months old) | Above also enriched + many other gene sets, see Fig. 7 for more details) |
| Zebrafish *psen1*Q96K97del/+<br>model<br>(see Fig. 9) | Q96K97del/+ vs. WT (6 months) | ↓ Oxidative Phosphorylation<br>↓ Fatty Acid Metabolism<br>MTORC1 signalling<br>PI3K AKT MTOR signalling |
|  | Q96K97del/+ vs. WT (24 months) | Above also enriched + many other gene sets, see Fig. 9 for more details) |

### Simultaneous stabilization of some 3’ IRE transcripts and destabilization of others

In the cultured cell line dataset analyzed above, we noticed that even a straightforward iron deficiency treatment resulted in the simultaneous increase and decrease in expression of 3’ IRE-containing genes. These findings are difficult to reconcile with the current paradigm that stabilization of 3’ IRE-containing genes occurs under iron deficiency and suggest that the current model of the IRP/IRE system may be incomplete or insufficient for describing the regulation of a broader collection of IRE genes. Given that many predicted 3’ IRE genes with non-canonical IREs (e.g. in the ***all 3’ IREs*** set) displayed enrichment and different gene expression patterns in the different genotype/age/oxygenation groups of the fAD-like zebrafish dataset, we decided to explore further the stability changes of these genes by comparing the expression of spliced and unspliced transcripts for each gene (**Supplementary Text 4**). We found that transcripts of some predicted 3’ IRE genes were significantly increased in stability while others were significantly decreased in stability (**Supplementary Figure 7**; **Supplementary Table 5**).

### Some validated IRE genes demonstrate expression changes inconsistent with the classic IRE paradigm

The work described above discovered both increased and decreased stability of transcripts predicted to possess IREs in fAD-like zebrafish brains relative to wild-type brains. However, it did not address directly whether such changes in stability occur in transcripts confirmed as targets of IRP binding and in systems subjected to disturbance of iron homeostasis. To support that our sets of 5’- and 3’-IRE-containing genes include both transcripts that increase in abundance and transcripts that decrease in abundance as iron levels change, we examined several IRE genes previously demonstrated by Sanchez et al. [24] to produce transcripts bound by IRPs (see below). We examined these transcripts in transcriptome datasets produced from cell lines subjected to changes in iron levels (**Figure 11**).

**Figure 11.** Expression of validated IRE genes in two cell line datasets. **A. Summary of workflow.** mRNAs found to bind mouse IRP1 and/or IRP2 via immunoprecipitation in Sanchez et al. [24] were filtered for those which contained a high-quality IRE in their 3’ or 5’ UTR in human and mouse Ensembl gene models. Expression of these validated IRE genes was examined in a mouse splenic B cell dataset and a human Caco-2 cell line dataset. Within each dataset, *t*-tests were used to test for significant expression differences of validated IRE genes between iron deficiency-treated cells and control cells. *t*-test *p*-values ≤ 0.05 were considered to indicate statistically significant differences. Statistically significant 3’ IRE genes were categorized as either “consistent with the IRE paradigm” (increased expression in iron deficiency-treated cells) or “inconsistent with the IRE paradigm” (decreased expression in iron deficiency-treated cells). 5’ IRE genes were not expected to show significant differences according to the IRE paradigm, but 5’ IRE genes showing significant differences in either direction in iron deficiency-treated cells were also recorded and plotted. **B. Expression of validated IRE genes showing statistically significant differences in iron deficiency-treated human Caco-2 cells.** Iron deficiency-treated cells refers to the “Iron-free Medium” cells relative to “Iron-free Medium + FAC” cells. Numbers over boxplots are *p*-values from *t*-tests between groups. **C. Expression of validated IRE genes showing statistically significant differences in iron deficiency-treated mouse splenic B cells.** Iron deficiency-treated cells refers to the deferoxamine (DFO)-treated cells relative to control cells. *p*-values are from *t*-tests between DFO-treated and control cells.

In Sanchez et al. [24], a total of 258 mRNAs were identified that immunoprecipitated with mouse IRP1 and/or IRP2 (44 immunoprecipitating with both IRP1 and IRP2, 113 immunoprecipitating exclusively with IRP1, and 101 immunoprecipitating exclusively with IRP2). We filtered these mRNAs for those with high-quality IREs (as predicted using SIREs) in the UTR of either the human or the mouse Ensembl gene model (**Figure 11A**). We found 30 mRNAs with high-quality IREs in human Ensembl gene models, and 50 mRNAs with high-quality IREs in mouse Ensembl gene models. These were then separated into those with IREs in their 3’ UTRs and those with IREs in their 5’ UTRs (see **Figure 11A** for more details), since transcripts with 3’ IREs are expected to show altered abundance under the classic IRE paradigm while the abundance of transcripts containing 5’ IREs is not expected to change (see **Figure 1**). Lists of experimentally-validated IRE genes filtered using these steps are available in **Supplementary Table 6**.

We then examined the expression of these experimentally-validated IRE-containing genes in transcriptome datasets from human and mouse cells subjected to iron level-altering treatments. We sought to observe whether transcript abundance differences were consistent with those predicted under the classic IRE paradigm. The datasets included the human Caco-2 cultured cell line dataset (previously described above), and a mouse splenic B cell dataset [76] (GEO Accession: <u>GSE77306</u>).

The Caco-2 dataset includes both iron overload (DMEM-FBS Medium + Hemin vs. DMEM-FBS Medium) and iron deficiency (Iron-free Medium vs. Iron-free Medium +FAC) treatments, while the mouse B cell dataset includes only a treatment with deferoxamine (DFO) to create iron deficiency (DFO-treated cells vs. control). We note that the DFO and control samples in the mouse B cell dataset are not directly comparable within each group, due to the unstimulated or stimulated anti-IgM or LPS treatments also used in this study. However, a Principal Component Analysis plot of the samples shows that the DFO-treated samples apparently separate from the control samples across Principal Component 2, indicating that DFO-treatment is likely to be one of the main causes of differential gene expression in this dataset (**Supplementary Figure 8**).

Overall, in the mouse B cells and human Caco-2 cells affected by changes in iron levels, approximately half of the genes identified with validated IREs show significant differential expression. Of those, similar proportions show increased or decreased transcript abundance in either the 3’ IRE or 5’ IRE groups and, of those, there are numerous genes showing changes in expression inconsistent with the IRE paradigm. In particular, in the Caco-2 cells, three 3’ IRE genes (*EIF4EB2, LAS1L,* and *ZNF207*) show decreased expression in cells treated to create iron deficiency (**Figure 11B**; t-test *p*-values ≤ 0.05). In the mouse B cells, DFO-treatment to cause iron deficiency increases the abundance of transcripts of four 3’ IRE-containing genes, including *Tfrc,* in accordance with the classic paradigm (**Figure 11C**; *t*-test *p*-values ≤ 0.05). In contrast, transcripts of four such 3’ IRE genes show decreased abundance (*Bub3, Cxcl16*, *Gstm6*, *Zfp71-rs1*) (**Figure 11C**; *t*-test *p*-values ≤ 0.05).

Although these sample sizes are small, they clearly indicate that the presence of an IRE in either the 5’ or 3’ UTR of a transcript cannot, by itself, be used to predict how the abundance of the transcript will change in response to iron availability.

## Discussion

In this study we identified comprehensive sets of genes predicted to contain IREs in humans, mice, and zebrafish. The idea of analysing gene expression datasets using gene sets is not novel, with many well-established techniques for gene set testing [35–39] that offer the opportunity to analyze the behavior of large groups of genes to detect statistically significant changes in state, even though these may be quite subtle for individual genes. **Table 5** summarises representative enriched Hallmark gene sets (representing common biological pathways/activities) across the human, mouse, and zebrafish datasets analysed in this work. Surprisingly, we found that IRE gene sets are generally not well-represented in existing gene sets. The existing gene set in which IRE genes are most highly represented is one previously shown to be up-regulated in AD brains (the “Blalock Alzheimer’s Disease Up” gene set from MSigDB). This supports a unique importance of IRE genes in AD. Consequently, the IRE gene sets identified here represent a novel resource for researchers interested in exploring iron dyshomeostasis gene expression responses in detail in different species, conditions, and treatments. In our results, we demonstrate that IRE gene sets are sufficiently sensitive to reveal broad differences across different conditions (both in human AD and animal models), while also offering the opportunity to study individual genes/transcripts contributing to enrichment in more detail. IRE gene sets displayed significant enrichment in postmortem brains from a human AD cohort, the 5XFAD mouse model, and a zebrafish model of a fAD-like mutation. Taken together, our results demonstrate, for the first time, the involvement of coordinated IRP/IRE-mediated gene expression responses associated with AD.

Overall, IRE gene sets were sufficiently sensitive to reveal several interesting phenomena. The relatively tightly controlled conditions in the mouse and zebrafish model datasets revealed strongly age-dependent effects on the transcript abundances of these predicted IRE-containing genes. Notably, 3’ IRE gene expression changes were amongst the earliest changes observable in the zebrafish heterozygous single fAD-like mutation model alongside changes in energy metabolism. These 3’ IRE gene expression changes preceded other signals of pathological change in the transcriptome (such as altered expression of inflammatory response pathways) commonly associated with AD. Our findings raise the question of how early IRE gene expression changes, and iron dyshomeostasis more broadly, may begin in fAD brains. In a recent analysis of transcriptome data from pools of 7- day-old zebrafish larvae heterozygous for the fAD-like mutation *psen1*^Q96_K97del^ (compared to wild type half-siblings), we observed over-representation of the GO term “iron ion transport” without significant changes in 3’ IRE or 5’ IRE transcript abundance [47]. This supports that the 3’ IRE transcript abundance signal in 6-month-old brains of this mutant line reflects changes in ferrous iron abundance that develop with age in response to an underlying defect in cellular iron regulation.

In relation to our findings associated with the 5XFAD mouse model, we acknowledge that low sample number is a limitation in this analysis and constrains interpretation of our results. Unfortunately, to our knowledge, no other publicly-available 5XFAD mouse dataset was available containing (1) both young and aged adults in the same batch (where young mice are defined as ∼2-4 months old as this is when the earliest signs of pathology are observed for this model [77, 78]), (2) 5XFAD and WT genotype mice in the same batch, (3) information about biological sex for each sample (so that sex-specific effects could be accounted for in the statistical model for differential gene expression), and (4) separation of groups by biological factors (e.g. age, genotype) rather than by unidentifiable batch effects when inspected through Principal Component Analysis. Our original rationale for analyzing the 5XFAD mouse model came from results from an independent concordance analysis of various mouse models and human AD [67]. In that study, it was found that 5XFAD mouse brain transcriptomes showed greatest (although still fairly poor) concordance to human AD compared to other mouse models including the 3xTg model. However, we also acknowledge the limitations associated with only analyzing the 5XFAD mouse model here. It would be useful to analyze other mouse models of AD in the future to compare the involvement of IRE genes across different models. Likewise, we anticipate that further analysis of fAD-like mutation models in general will help to extend our knowledge of the early molecular changes contributing to the development of AD.

Notably, both classic IRE genes (e.g. *TFRC*, *FTL*, and *FTH1*) and the IRE gene sets were sufficiently sensitive to distinguish not only between iron overload and deficiency in a cultured cell line dataset, but also between AD and other neuropathologies (i.e. PA and PSP). This suggests that the dysregulation of IRE-containing genes, (and iron homeostasis in general), in AD may differ from other neuropathological conditions. The differences in both classic IRE genes and IRE gene set expression between AD and PA are of particular interest. Considering that presence of amyloid pathology is required to diagnose individuals with AD [79, 80] and contributes to diagnosis of preclinical AD [1, 2], it will be interesting to explore further the biological relevance and impact of these IRE gene expression differences using other techniques to gauge whether PA is a prodrome of, or a separate pathological condition to, AD.

One of our original intentions for IRE gene set analysis of the human AD and animal model datasets was to gain insight into whether these brains were in a state of iron deficiency or overload using gene expression-level evidence. Unfortunately, it became clear that interpretation of IRE gene expression changes was more complex than expected, both because of the use of bulk brain tissue samples in the human and animal model datasets (which obscures differences between cell type and tissue types) and because the regulation of both 3’ and 5’ IRE-containing genes we observed did not follow the expected paradigm. To address the former issue, future work should expand the use of IRE gene sets to single-cell RNA-seq (scRNA-seq) datasets so that the IRE gene expression patterns of different cell types can be deconvoluted. Recently, a large-scale scRNA-seq study of AD brains indicated highly cell-type specific changes related to inflammation during early stages of disease progression within the prefrontal cortex [81], highlighting the importance of studying iron homeostasis responses at the single-cell level in the context of AD. The failure of 3’ IRE genes to follow the expected expression paradigm is a limitation we did not anticipate when we began this work. In particular, our observations indicating that (1) the stability of some 3’ IRE transcripts (rather than their “expression”/“levels” as addressed in preceding analyses) is significantly different under different conditions in zebrafish, and that (2) validated 3’ IRE genes show both increased and decreased expression under iron deficiency in two cell line datasets (suggesting they are not stabilized as predicted by the IRE paradigm), challenge the simplistic generalisation that 3’ IREs stabilise transcripts under ferrous iron deficiency.

Previous research found that, under iron deficiency, the transcripts of 3’ IRE-containing genes such as *TFRC* are stabilized causing increased expression [20] to increase ferrous iron availability. However, our findings reveal that this principle (the “classic IRE paradigm”) does not necessarily apply to other 3’ IRE-containing genes, including those with canonical IREs (e.g. *SLC11A2* in the Caco-2 cultured cell line), and genes with IRE-like elements deviating from the canonical IRE sequence. Interestingly, Tybl et al. [82] found that the 3’ IRE in *SLC11A2* (*DMT1)* appears to influence transcript abundance in a development- specific manner in mice, by promoting transcript expression during postnatal growth and suppressing expression in adulthood. Together with our findings, this demonstrates that the regulation of IRE genes is more complex than commonly appreciated.

Aside from 3’ IRE genes, 5’ IRE genes also showed unexpected behavior in the datasets we analyzed. In general, the classic IRE paradigm holds that binding of IRPs to 5’ IREs blocks translation but is not expected to change transcript abundance. We initially sought to investigate 5’ IRE transcripts as a negative control group to which changes in the stability of the 3’ IRE transcripts could be compared. Unexpectedly, our results indicated that 5’ IRE gene sets are significantly enriched under various conditions. Upon reflection, this is not unreasonable, considering that binding of any protein to a transcript must, inevitably, changes the transcript’s stability to some degree. However, the direction of that stability change is difficult to predict as there is no generalizable cellular “rule” for the outcome of protein binding. This is consistent with the reality that each gene represents a largely independently evolving system constrained only by cells’ requirements for its functionality. For example, inactivation of RNA-binding proteins has been demonstrated previously to cause both up- and down-regulation of different genes simultaneously [83]. In addition, the 5’ canonical IRE and surrounding acute box sequences of the H-ferritin (*FTH*) transcript have been shown to stabilize it, which might appear contrary to the expected function of a 5’ IRE (of decreasing protein translation under iron deficient conditions) [84]. H-ferritin is one of the better-characterised IRE genes, and it is possible that other, yet-to-be characterised regulatory mechanisms involving 5’ IREs could contribute to stability changes in other transcripts. Aside from 3’ IREs, other 3’ UTR regulatory motifs are known that bind to proteins to regulate multiple biological activities (e.g. AU-rich 3’ UTR motifs as described in [85]). Therefore, mechanistically, it is reasonable to assume that IRE-based regulation of transcript stability may be co-opted for other biological activities beyond iron homeostasis (and see below). Alternative splicing of IRE genes may be one mechanism for this, where a non-IRE-containing isoform is expressed in certain tissues or conditions, as is the case for mammalian *DMT-1* (*SLC11A2*) [22]. Differential transcript isoform inclusion of IREs is beyond the scope of our current work, but will be important to explore in future. We expect our findings to provide a starting point for further research involving IRE genes and their complex regulation beyond the classic paradigm.

It is important to acknowledge that some of the complexity involved with IRE gene expression, and the large number of genes identified with putative IREs, may reflect the involvement of IREs in functions other than iron homeostasis. These functions include regulation of oxidative stress [86–88]), oxidative phosphorylation [89], and cell cycle control [90, 91]. In 2019, it was proposed that the tricarboxylic acid cycle (TCA) may have evolved around the reactivity of iron [92], and TCA intermediates themselves also perform important cellular regulatory functions [93]. In addition, our analyses identified several transcription factor motifs potentially associated with other biological pathways as over-represented in IRE gene sets. Similarly, although beyond the scope of our current work, a regulatory role for micro RNAs (miRNAs) in binding IRE structures is emerging [94–97]. This may be particularly relevant to AD, as a miRNA has been described that can bind the 5’ IRE-like structure in AβPP transcripts to up-regulate AβPP its expression [98]. Co-regulation of IRE-containing genes within other biological pathways may explain some of the IRE gene set enrichment observed in the datasets we analyzed, and some of the unexpected expression changes (e.g. decreased expression of 3’ IRE genes in conditions where stabilization of their transcripts might be expected, and increased expression of 5’ IRE genes even when no change in stability would be expected). Hence, the changes in expression of predicted IRE genes we observed must be interpreted carefully as reflecting not only iron dyshomeostasis responses, but also other co-occurring changes in linked processes. This was particularly evident in our analysis of the Caco-2 cultured cell line dataset where even straightforward iron overload and iron deficiency treatments resulted in significant enrichment of gene sets related to reactive oxygen species and hypoxia pathways. We note that the relationship between iron homeostasis and hypoxia would be interesting to explore in future work, particularly their similarities at the the level of gene expression. Previous research has demonstrated that desferrioxamine (that induces iron deficiency) can also induce HIF-1 activity, which activates erythropoietin (important for red blood cell formation to carry oxygen) [99]. This supports that iron deficiency and hypoxia involve similar molecular changes in cells (i.e. the view that iron deficiency generates a form of “pseudohypoxia”). We expect that future work deconvoluting the expression of IRE genes at the cellular level will be required to determine to what extent such expression changes are influenced by cell / tissue type compared to other linked biological activities.

Most previous work has assumed that accumulation of iron in the brain with age (a phenomenon observed broadly across animal phyla [100, 101]) is indicative of cellular iron overload driving oxidative stress [102]. Indeed, iron accumulation and oxidative stress apparently contribute to the insolubility of amyloid plaques and neurofibrillary tangles in AD brains [103]. However, a recent publication by Yambire et al. [104] showed disturbed lysosomal function leading to increased lysosomal pH and causing a deficiency of functional ferrous iron (Fe^2+^) while non-functional ferric iron (Fe^3+^) accumulated in lysosomes. The deleterious effects of this on the brains of mice (defective mitochondrial biogenesis and function and stimulation of inflammatory responses) could be alleviated by dietary iron supplementation. The observations of Nixon and colleagues that acidification of the endolysomal pathway is affected both by fAD mutations in *PSEN1* [71] and excessive dosage of the amyloid beta A4 precursor protein (*AβPP*) gene [105], together with our observations from our fAD-mutation-like *psen1*^Q96_K97del^/+ mutant zebrafish in both this study and in ref. [44] (where gene expression changes associated with lysosome and mitochondria for this model were first described), support the possibility that AD brains may suffer a ferrous iron deficiency in a background of ferric iron overload. This is discussed further below.

Although the evidence from human AD brains is more tentative at this stage, our findings of increased expression of *TFRC* expression and other well-characterized 3’ IRE genes in the temporal cortex tissue of human AD brains is consistent with the existence of a cellular ferrous iron deficiency, although more research is required to confirm this at the single-cell level. Intriguingly, the greatest genetic risk factor for late onset AD, the *ε*4 allele of the gene *APOE*, appears to increase lysosomal pH [106] but *ε*4’s increased risk of AD is alleviated in individuals who possess the *HFE* 282Y allele that predisposes to the iron overload disease hemochromatosis [107]. Given that many other risk loci for sporadic late onset AD also affect endolysosomal pathway function (reviewed in [108]), it is reasonable to suggest that disturbed iron homeostasis may afflict brains with this disease.

Our work provides evidence consistent with Yambire et al. [104] and Lee et al. [71] to suggest that fAD mutations may drive a ferrous iron deficiency that contributes to AD pathology. However, iron export is a complex process, and it remains unclear whether iron deficiency or overload is more relevant to the pathology seen in AD. Much of the existing research has focused on the role of AβPP and its contributions to elevating iron levels within cells and to increasing iron export to the cerebral interstitium in AD. For example, Venkataramani et al. [109] demonstrated that decreased AβPP levels resulted in stabilisation of ferroportin, resulting in increased iron export from neuroblastoma cells in two mouse models. Later work by McCarthy et al. [110] proposed a mechanism for this phenomenon, involving a motif in sAβPPa (cleavage product of AβPP) binding to ferroportin and increasing iron export from cells through increasing the number of ferroportin molecules in the plasma membrane but not their activity. A complicating factor is that ferroportin-mediated iron export from cells requires both (1) stabilisation of ferroportin in the membrane, and (2) ferroxidase activity (provided by either haephaestin or ceruloplasmin) [110]. The role of haephestin as a ferroxidase in this process of iron export from cells has sometimes been overlooked, despite that it is well-established as complexing with, and stabilizing, ferroportin to export iron [111–113]). For example, recently Tsatsanis et al. [114] described the effects on intracellular iron and iron export of changes in amyloidogenic cleavage (via β-secretase) and non-amyloidogenic cleavage (via γ-secretase) of AβPP. It was observed that amyloidogenic cleavage of AβPP led to destabilisation of ferroportin on the cell surface and increased iron retention in cells. In addition, inhibiting β-secretase mitigated this effect, increasing ferroportin stabilization and decreasing iron retention in cells. However, an alternative explanation not considered by those authors is that haephestin may also be cleaved by β-secretase (see Figure 7 in [115]), so it is possible that reduced β-secretase activity could increase the haephestin available and also contribute to the effects on iron export observed. Overall, the situation around iron export from cells is a complex area of ongoing research involving many cellular factors, and it will be increasingly important in future work to understand these mechanisms more completely in the context of AD.

We have previously noted that all the fAD mutations of AβPP (including gene duplications) are expected to have the singular quality of increasing the abundance of the amyloidogenic β-CTF / C99 fragment of AβPP (see Figure 2 in Lumsden et al. [14]). Recently Jiang et al. [105] showed that increased β-CTF reduces lysosomal acidification. According to the results of Yambire et al. cited above, this would be expected to reduce cellular ferrous iron levels (while causing accumulation of ferric iron in lysosomes). This idea is consistent with the existence of an IRE in the 5’ UTR of AβPP transcripts that inhibits AβPP translation (and hence β-CTF production) when cytosolic ferrous iron levels are low [116]. Thus, AβPP may influence cellular ferrous iron levels by modulating lysosomal pH to regulate importation of ferrous iron into the cytosol, rather than by modulating ferroportin stability to regulate iron export from the cytosol.

Overall, our results demonstrate for the first time the involvement of a coordinated IRP/IRE-mediated gene expression response in the context of AD. By searching entire human, mouse, and zebrafish transcriptomes for all genes containing potential IREs, we formed comprehensive gene sets applicable to the analysis of any human, mouse, or zebrafish gene expression dataset. For the first time, our work suggests the existence of gene expression changes associated with the IRP/IRE system in the young, pre-pathology brains of *PSEN1* fAD mutation carriers, and is consistent with the idea of iron dyshomeostasis as an early contributor to the disease process. It also reinforces the critical importance of IRE genes in general in the development of the late onset, sporadic form of AD. More broadly, our approach highlights how changes in the stability and abundance of IRE-containing transcripts can be used to give insight into iron dyshomeostasis-associated gene expression responses in different species, tissues, and conditions.

## Acknowledgements and Funding

The authors thank the Carthew Family and Prof. David Adelson for their encouragement and helpful discussions. This work was financially supported by grants GNT1061006 and GNT1126422 from the National Health and Medical Research Council of Australia (NHMRC) and by funds from the Carthew Family Charity Trust. Lead author NH was supported by an Australian Government Research Training Program Scholarship and by the South Australian Genomics Centre (SAGC) post-graduation. Co-author MN was supported by funds from the grants listed above. Joint senior co-authors SP and ML were/are academic employees of the University of Adelaide respectively. The authors also thank the University of Adelaide’s Phoenix HPC team for providing supercomputing resources for this work.

## Declarations

### Ethics approval and consent to participate

No experiments of human subjects were performed in this study as all human data was accessed from public databases and was anonymous. All zebrafish work was conducted under the auspices of the Animal Ethics Committee (permit numbers S-2017-089 and S- 2017-073) and the Institutional Biosafety Committee of the University of Adelaide.

### Consent for publication

All authors give consent for publication of the manuscript.

### Datasets and Data Articles

The fAD-like zebrafish dataset supporting the conclusions of this article is available in the GEO repository with accession number GSE149149.

### Authors’ contributions

NH performed the bioinformatics analysis and drafted the manuscript. MN generated the mutant zebrafish, isolated brain RNA and protein, drafted the description of this and edited the manuscript. SP directly supervised NH, guided the analysis, and edited the manuscript. ML conceived the project, supervised all aspects of it, and edited the manuscript. All authors read and approved the manuscript.

## Supplementary Information

### Supplementary Text

- **Supplementary Text 1.** Effects of biological sex on gene expression in the fAD-mutation-like zebrafish dataset.
- **Supplementary Text 2.** Neural cell type proportions in the gene expression datasets analyzed in this paper.
- **Supplementary Text 3.** Expression of previously characterized 3’ and 5’ IRE genes in the gene expression datasets analyzed in this paper
- **Supplementary Text 4.** Identifying predicted 3’ IRE genes with altered stability.

### Supplementary Tables

- **Supplementary Table 1.** Iron Responsive Element (IRE) gene sets defined from human, mouse, and zebrafish transcriptomes. The untranslated regions (UTR) of all known genes were searched for IRE and IRE-like motifs. Four gene sets are included for each species: ***all 3’ IREs*** (all genes with predicted IRE in their 3’ UTR), ***all 5’ IREs*** (all genes with predicted IRE in their 5’ UTR), ***HQ 3’ IREs*** (genes with high-quality predicted IRE in 3’ UTR), and ***HQ 5’ IREs*** (genes with high-quality predicted IRE in 5’ UTR). Genes are provided as ENSEMBL gene identifiers.
- **Supplementary Table 2.** Molecular Signature Database (MSigDB) gene sets with significant over-representation of IRE gene sets in human, mouse, and zebrafish. Gene sets were defined to have significant over-representation of IRE gene sets if the FDR-adjusted Fisher’s exact test *p*-value was < 0.05.
- **Supplementary Table 3**. Promoter motif over-representation analysis results for all IRE gene sets in human, mouse, and zebrafish. Promoter regions were defined as being 1500 bp upstream and 500 bp downstream of genes. See Methods for details. We define significant over-representation as FDR-adjusted enrichment *p*-value < 0.05.
- **Supplementary Table 4.** Gene ontology (GO) analysis results for leading edge genes shared between the “5XFAD vs. WT (3-months-old)” and “5XFAD vs. WT (6- months-old)” comparisons. Significant over-representation of GO terms was determined as GO terms having an FDR-adjusted *p*-value < 0.05.
- **Supplementary Table 5.** Altered stability of transcripts in the fAD-mutation-like zebrafish dataset. See Methods and Supplementary Text 4 for details on how stability was defined and calculated. Genes which contain a 3’ IRE (belonging to one of the 3’ IRE gene sets) or 5’ IRE (belonging to one of the 5’ IRE gene sets) are indicated with the “has3ire” or “has5ire” columns.
- **Supplementary Table 6.** Validated IRE genes with high-quality IREs in their 3’ or 5’ UTRs in human and mouse, based on Ensembl gene models. The genes in this table are derived from information contained within Figure 2 and Supplementary Tables S1 and S2 of Sanchez et al. [24], who used immunoprecipitation to determine mRNA transcripts binding to IRP1 and/or IRP2 in mouse.
- **Supplementary Table 7.** Gene set test results for all comparisons in the Mayo Clinic RNA-seq dataset. Gene sets tested are those in the Hallmark Collection from the Molecular Signatures Database. Gene sets were considered significantly enriched if their Bonferroni-adjusted combined p-value from *fry*, *camera*, and *fgsea* was < 0.05.

### Supplementary Figures

- **Supplementary Figure 1.** Expression of previously characterized IRE genes in datasets analyzed. **A.** Caco-2 cell line dataset. **B.** Mayo Clinic RNA-seq (Human) dataset – Temporal Cortex tissue. **C.** Mayo Clinic RNA-seq (Human) dataset – Cerebellum tissue. **D.** 5XFAD mouse dataset. **E.** fAD-mutation-like zebrafish dataset.
- **Supplementary Figure 2.** Principal Component Analysis of results from different gene set testing methods.
- **Supplementary Figure 3.** Analysis of cultured Caco-2 cell line dataset. **A.** Principal Component Analysis plot of gene expression in the cultured Caco-2 cell line dataset. **B.** Volcano plots indicating differential gene expression due to iron overload (DMEM-FBS Medium + Hemin vs. DMEM-FBS Medium) and iron deficiency (Iron-free medium vs. Iron-free medium + FAC) treatments in the Caco-2 cell line dataset.
- **Supplementary Figure 4.** Gene set enrichment testing results for the AD vs. control comparison in cerebellum and temporal cortex.
- **Supplementary Figure 5.** STRINGR protein-protein interaction network plot between the 19 shared leading-edge genes between the *psen1*^Q96_K97del/+^ and *psen1*^+/+^ comparisons at 6 and 24-months-old.
- **Supplementary Figure 6.** Principal Component Analysis plots showing lack of clear association between IRE gene expression and the *psen1*^K97fs/+^ mutant genotype.
- **Supplementary Figure 7.** Predicted 3’ IRE-containing transcripts with significant differences in stability between conditions in the fAD-like zebrafish dataset.
- **Supplementary Figure 8.** Principal Component Analysis (PCA) plot of splenic mouse B cell dataset (GEO Accession: GSE77306).

