## Supplementary Text 1 for "Iron Responsive Element (IRE)-mediated responses to iron dyshomeostasis in Alzheimer’s disease"

### Effect of biological sex on gene expression in the fAD-mutation-like zebrafish dataset.

In humans, women have a higher incidence of AD compared to men. This was our original rationale for using only female zebrafish in previous work with a different zebrafish model to minimize any chance of sex-based bias in gene expression (described in [1]). However, we did not exclude male zebrafish from the current work described in the main text as subsequent experience has shown that, like other teleost fish in general, the brain transcriptomes of zebrafish appear to show relatively few sex differences [2–4]. Our analysis of the fAD-mutation-like zebrafish dataset used in the current work also supports a lack of significant difference between male and female brains in the following ways:

1. Principal Component Analysis (PCA) of whole-brain gene expression across the entire dataset shows that age, oxygen level (acute hypoxia or normoxia), and genotype ($psen1^{Q96K97/+}$ or $psen1^{+/+}$) are responsible for the variation in the dataset captured by Principal Components 1-4. Within each group, there appear to be no distinguishable differences between male and female brains (see Figures below).
2. Differential gene expression analysis between male and female fish for each age group (6-months-old or 24-months-old) results in only 12 DE genes between male and female 24-month-old brains, and 10 DE genes between male and female 6-month-old brains (FDR-adjusted *p*-value < 0.05) (see Table of genes below). Differential expression analysis was performed in the same way as described in the Methods, but using the model ~0 + genderage, where "genderage" groups the samples into one of the following groups: 24-month female brain, 24-month male brain, 6-month female brain, 6-month male brain. The comparisons (contrasts) defined involved comparing 24-month female brains to 24-month male brains, and 6-month female brains to 6-month male brains. None of the genes identified as DE due to sex are significantly DE in any of the biologically-relevant comparisons described in the main manuscript involving age, hypoxia, and/or genotype. Compared to

the effects of age, hypoxia, and genotype (ranging from ~200 to ~2,000 DE genes), our results support the idea that sex differences are modest in this particular dataset and would be unlikely to contribute significantly to gene expression changes observed or the interpretation of these results.

**Principal Component Analysis (PCA)**

In the following PCA plots, the biological group ("Group") in the key denotes zebrafish brains which are either the *psen1*$^{Q96\_K97del/+}$ ("q96") or *psen1*$^{+/+}$ (wt) genotype, aged either 24-months-old ("24") or 6-months-old ("6"), and have either been exposed to acute hypoxia ("1") or are in a condition of normoxia ("0"). There are four samples (whole brains) for each biological group.

The PCA plots below show that age accounts for the largest source of variation in the data (Principal Component 1). Principal components 2, 3, and 4 incompletely capture variation due to genotype and/or hypoxia. Biological sex does not appear to associate with any of these biological variables (genotype, hypoxia, or age).

Principal components 1 and 2:

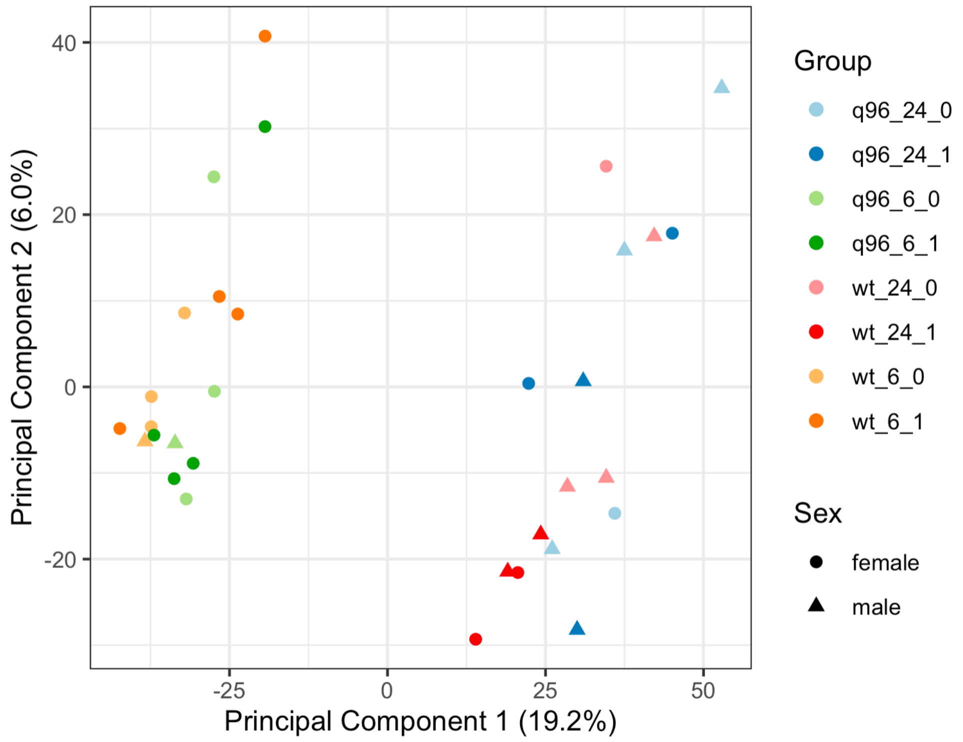

Principal components 1 and 3:

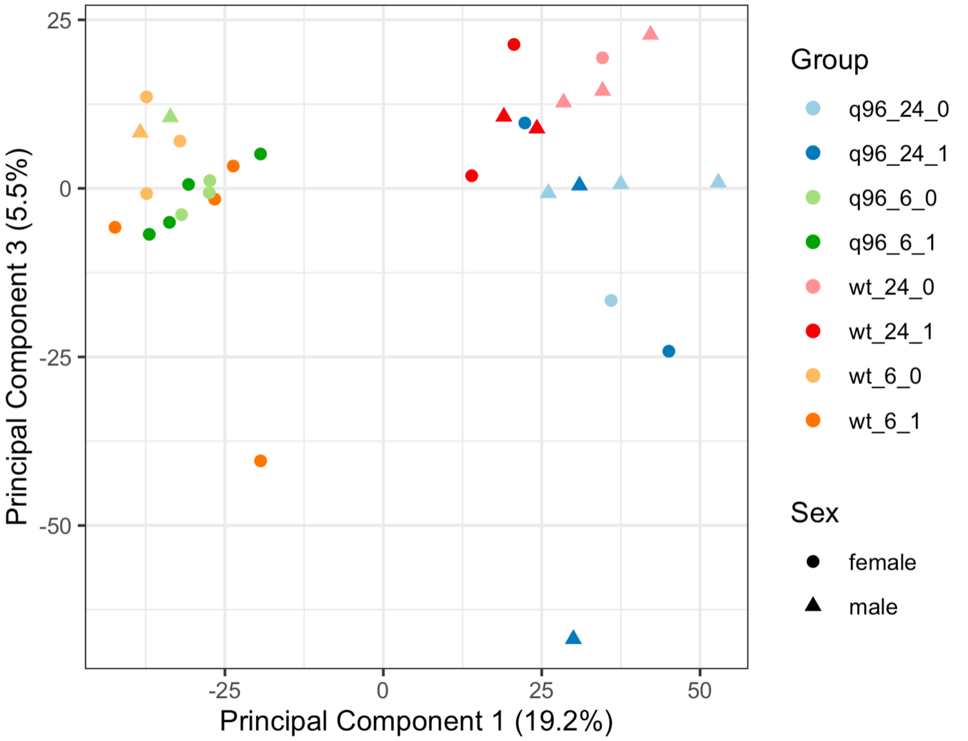


Principal components 1 and 4:

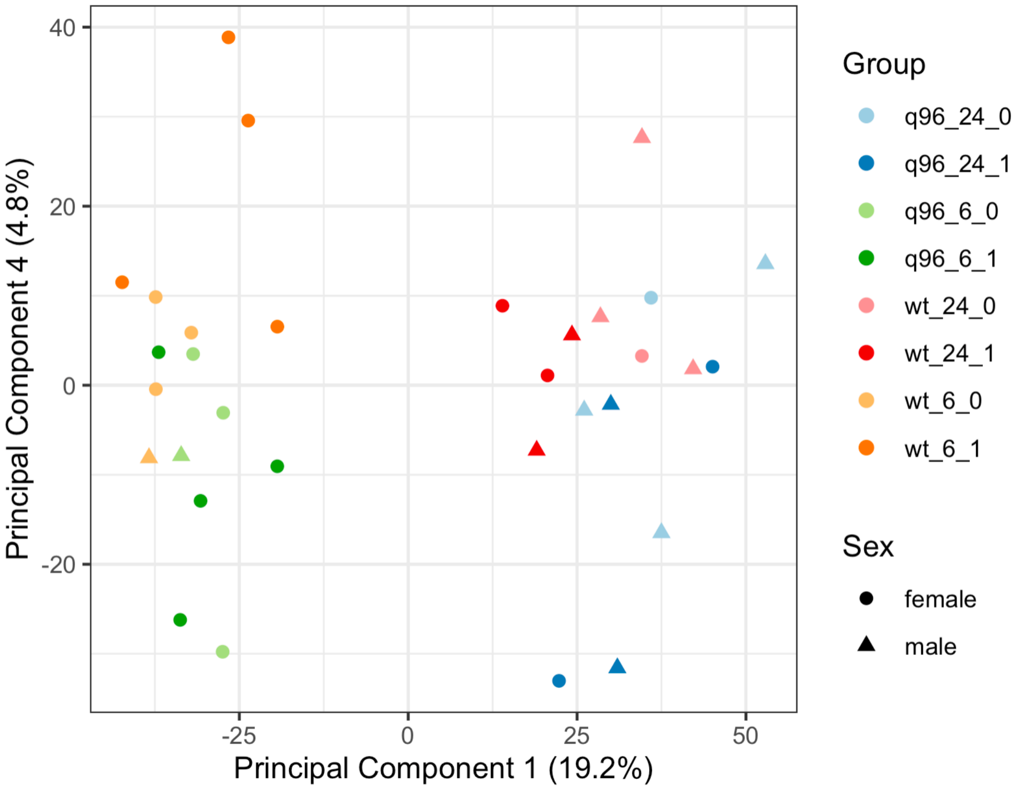

**Genes showing differential expression between female and male fish aged at either 24-months or 6-months in the fAD-mutation-like zebrafish dataset.**

Genes were defined as differentially expressed if the *limma* FDR-adjusted *p*-value < 0.05.

| Ensembl gene id | Gene name | logFC (24 month old) | logFC (6 month old) | FDR adj. *p* (24 month) | FDR adj. *p* (6 month) |
|---|---|---|---|---|---|
| ENSDARG00000003303 | stc1 | -0.0590089 | -0.9002048 | 1 | 0.01973971 |
| ENSDARG00000008912 | slc8a2a | -0.6207354 | -1.0184799 | 0.93448323 | 0.04005972 |
| ENSDARG00000012671 | inhbaa | -0.8852708 | -1.1368565 | 1.00E-03 | 2.28E-04 |
| ENSDARG00000016514 | gtf2h2 | -0.3257569 | -0.1394244 | 0.02198203 | 1 |
| ENSDARG00000023287 | hsd17b3 | -1.3108173 | -1.5393945 | 3.39E-09 | 3.28E-09 |
| ENSDARG00000038938 | itprid2 | -0.6445171 | -0.7411683 | 0.0017087 | 0.00700374 |
| ENSDARG00000046013 | rasl11a | -0.827503 | -0.861207 | 0.02113446 | 0.87564156 |
| ENSDARG00000053652 | ndufaf6 | -0.2763525 | -0.8948093 | 1 | 1.48E-04 |
| ENSDARG00000056510 | gstk1 | -0.7273964 | -0.4572104 | 0.00307532 | 1 |
| ENSDARG00000070717 | slc25a18 | -0.8550827 | -0.8958048 | 1.20E-08 | 3.97E-07 |
| ENSDARG00000078847 | si:dkey-238o13.4 | -1.446942 | -1.7137473 | 5.45E-04 | 3.28E-04 |
| ENSDARG00000092052 | gstk4 | -0.7080225 | -1.9402585 | 1 | 9.96E-05 |
| ENSDARG00000094132 | igf1 | -1.0524014 | -0.7939133 | 2.07E-05 | 0.13604296 |
| ENSDARG00000094857 | dio2 | -1.3364248 | -1.6631651 | 1.70E-04 | 6.54E-05 |
| ENSDARG00000104656 | si:dkeyp-97a10.3 | 3.59512879 | -0.3230445 | 0.04142206 | 1 |
| ENSDARG00000104735 | nkx6.2 | -0.4610814 | -0.1967992 | 0.00355871 | 1 |
