## Supplementary Text 2 for "Iron Responsive Element (IRE)-mediated responses to iron dyshomeostasis in Alzheimer’s disease"

### Neural cell type proportions in the gene expression datasets analyzed in this paper

Changes in gene expression seen in the human, mouse, and zebrafish datasets analyzed in this study may reflect changes in the proportions of cell types rather than changes in cellular gene expression. This is important considering a decrease in the number of neurons might be expected during neurodegenerative conditions. To explore possible changes in the proportions of cell types in the datasets analyzed, we compared marker gene expression for four common neural cell types (astrocytes, neurons, microglia, and oligodendrocytes) within each dataset (see plots on following pages).

The marker genes for astrocytes, neurons, and oligodendrocytes were derived from gene sets by Lein et al. [1] and are based on studies in mice (available on MSigDB). The marker genes for microglia were derived from Bonham et al. [2] and based on studies in humans and mice. All gene IDs were converted to human, mouse, or zebrafish Ensembl IDs where necessary using BioMart.

Overall, we noticed that overall expression of neural marker genes was largely stable across different conditions and ages in the zebrafish and mouse datasets, with little evidence of being systemically increased or decreased in 5XFAD or fAD-like conditions respectively (see following pages). In the human Mayo Clinic RNA-seq study however, we noticed trends suggesting that AD samples tended to express astrocyte marker genes at overall higher levels than controls. This information should be considered in interpretation of differential expression in this dataset.

**Human Mayo Clinic RNA-seq dataset**

Microglia marker gene expression (20 genes)

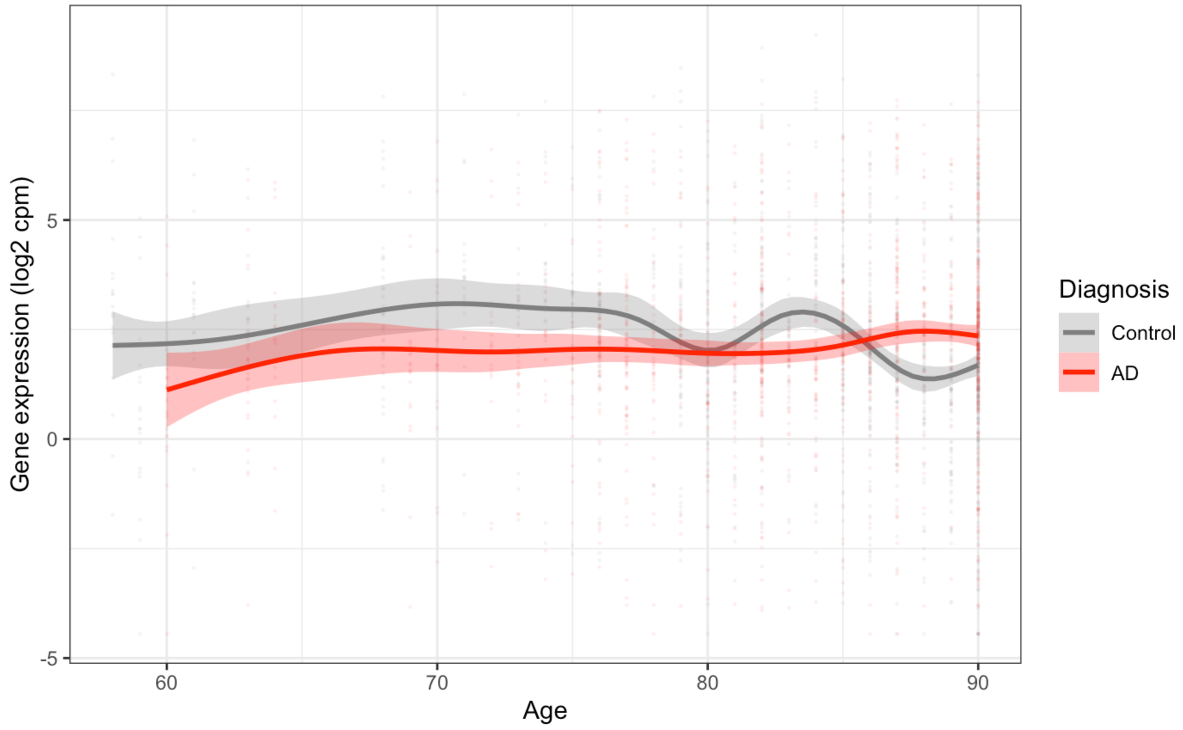


Astrocyte marker gene expression (44 genes)

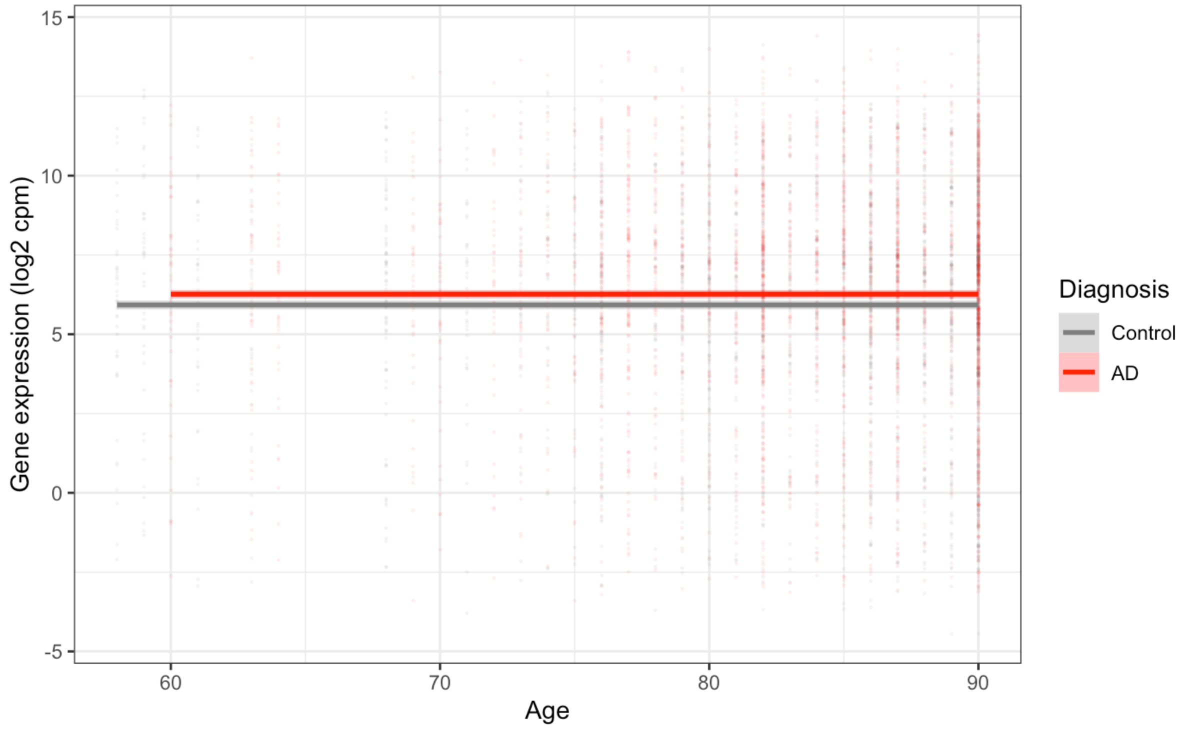

Neuron marker gene expression (74 genes)

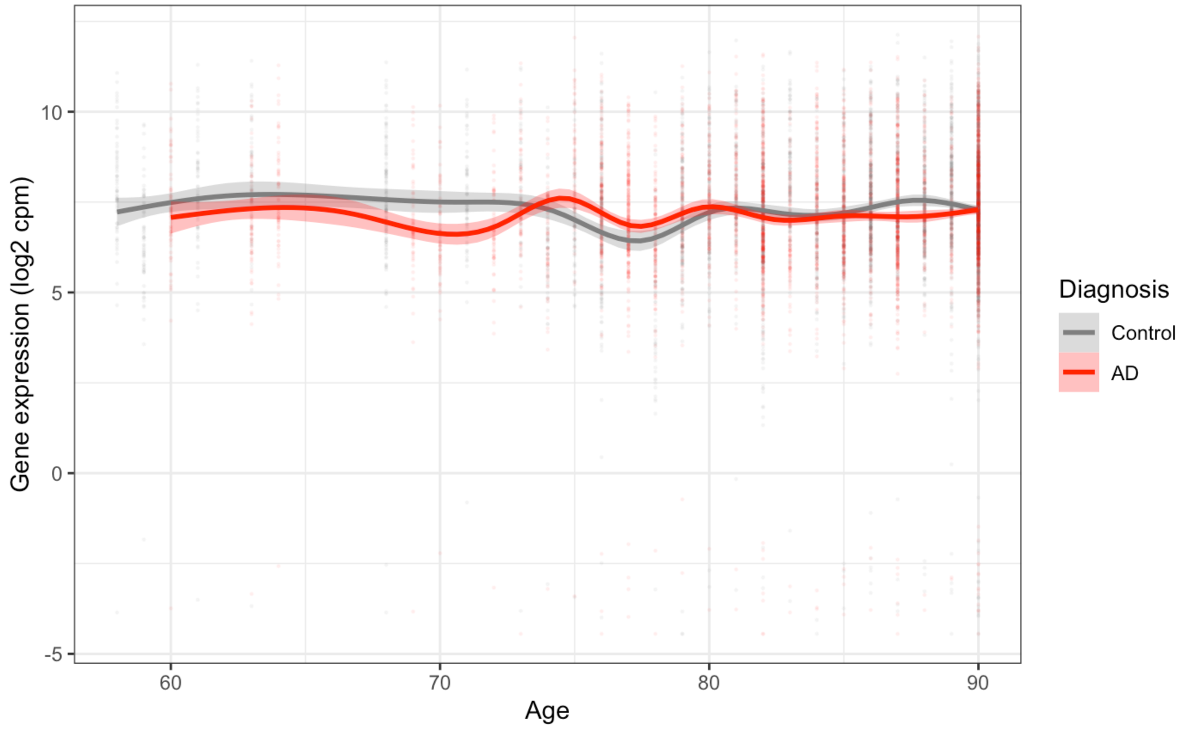


Oligodendrocyte marker gene expression (83 genes)

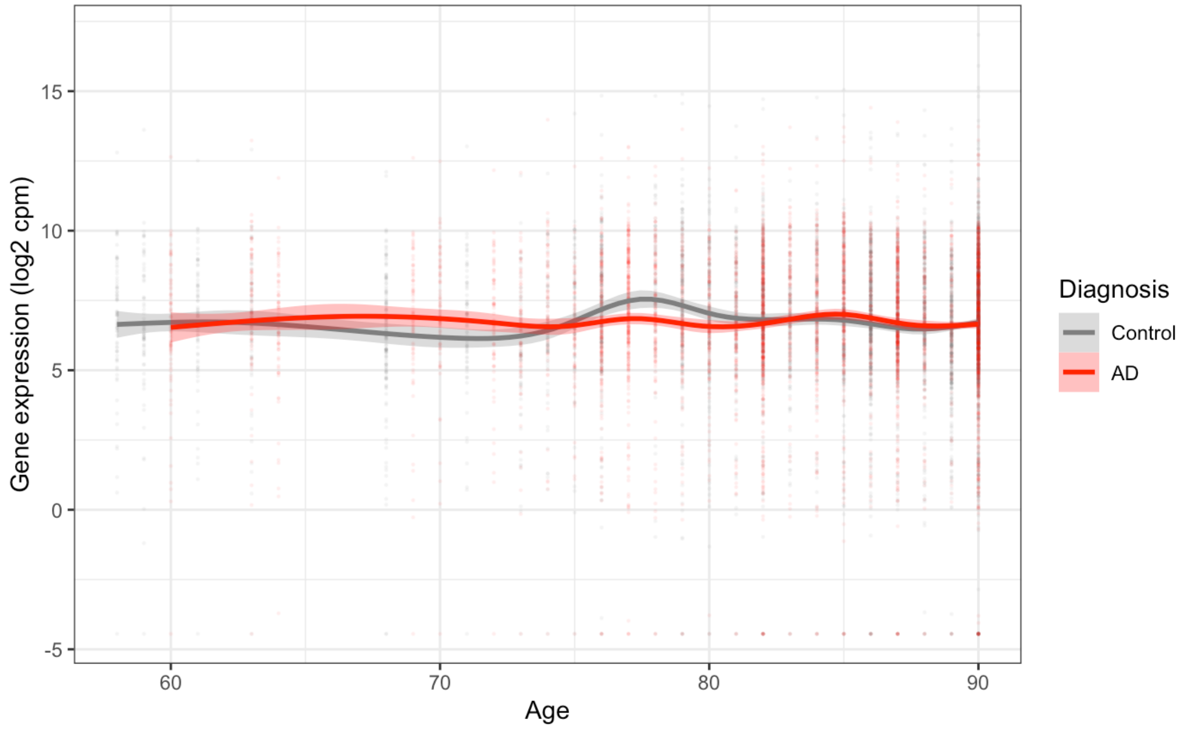


Error bars represent the 95% confidence interval of trend lines. Overall, while we see differences in expression, there does not appear to be an overall systematic difference between AD vs. control for oligodendrocyte, neuron and microglial marker gene expression. For the astrocyte markers, AD brains appear to systemically have higher expression of these genes.

**Mouse cortex dataset**

Microglial marker gene expression (22 genes)

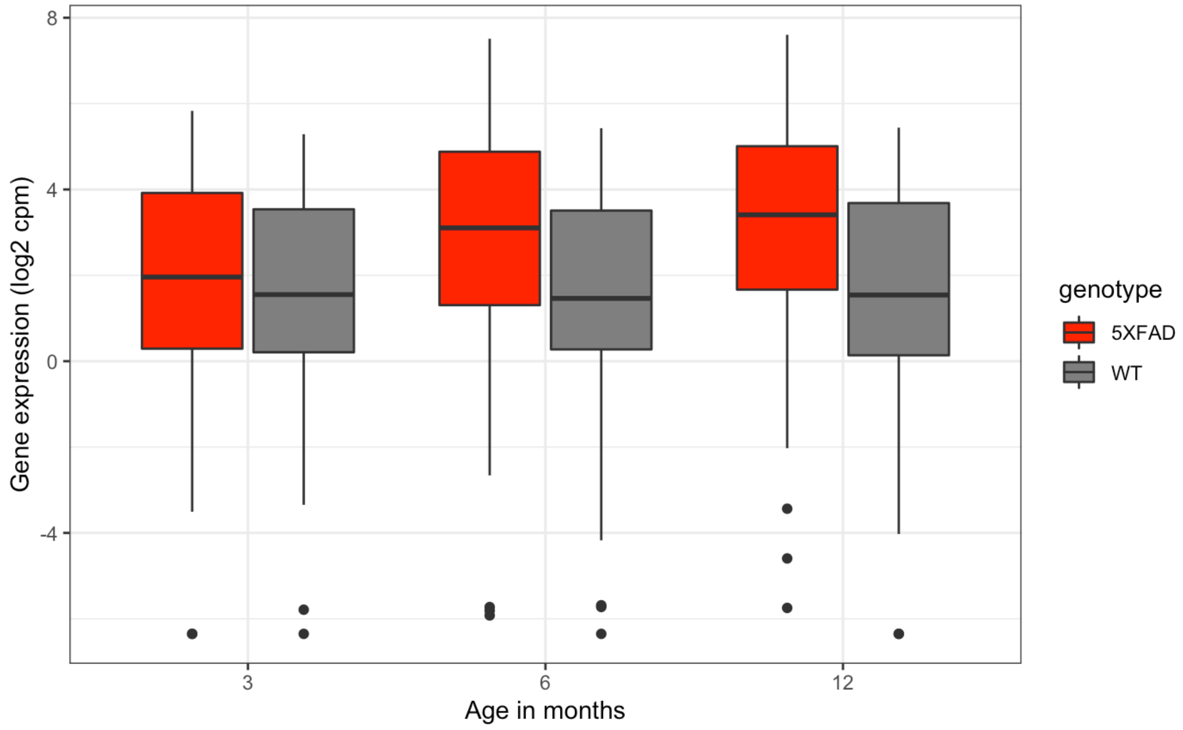


Astrocyte marker gene expression (51 genes)

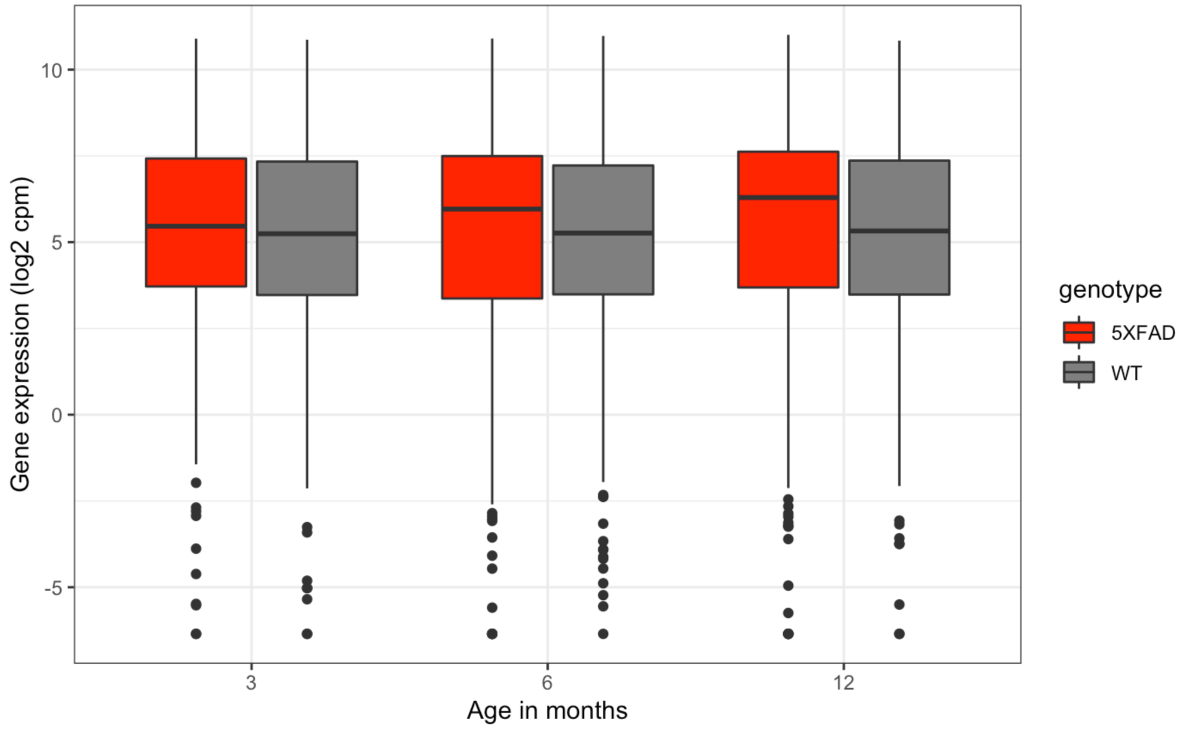

Neuron marker gene expression (67 genes)

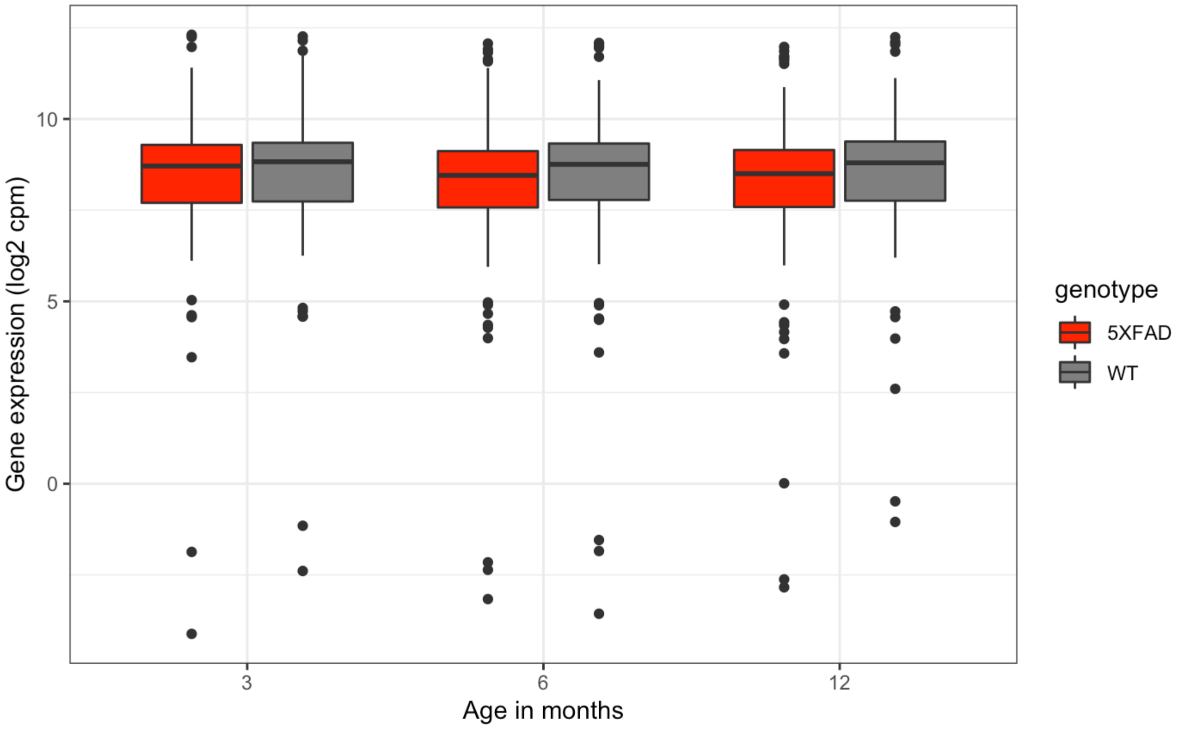


Oligodendrocyte marker gene expression (77 genes)

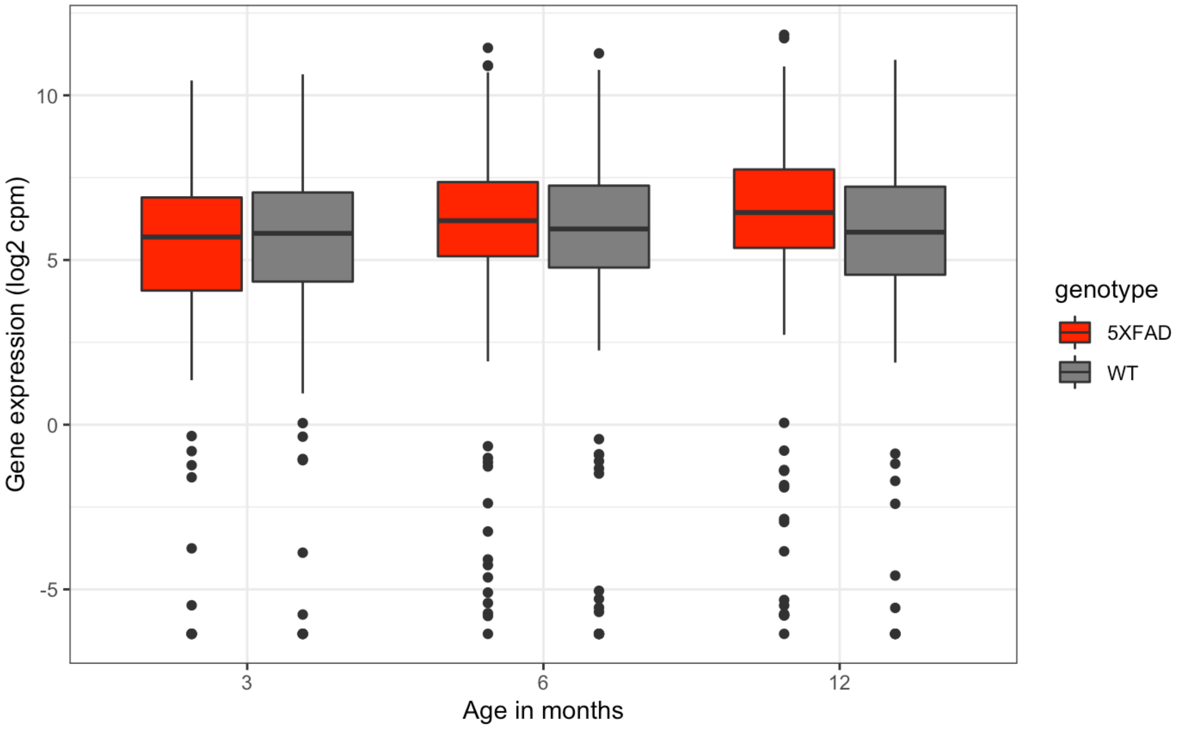

**Zebrafish dataset**

Microglial marker gene expression (16 genes)

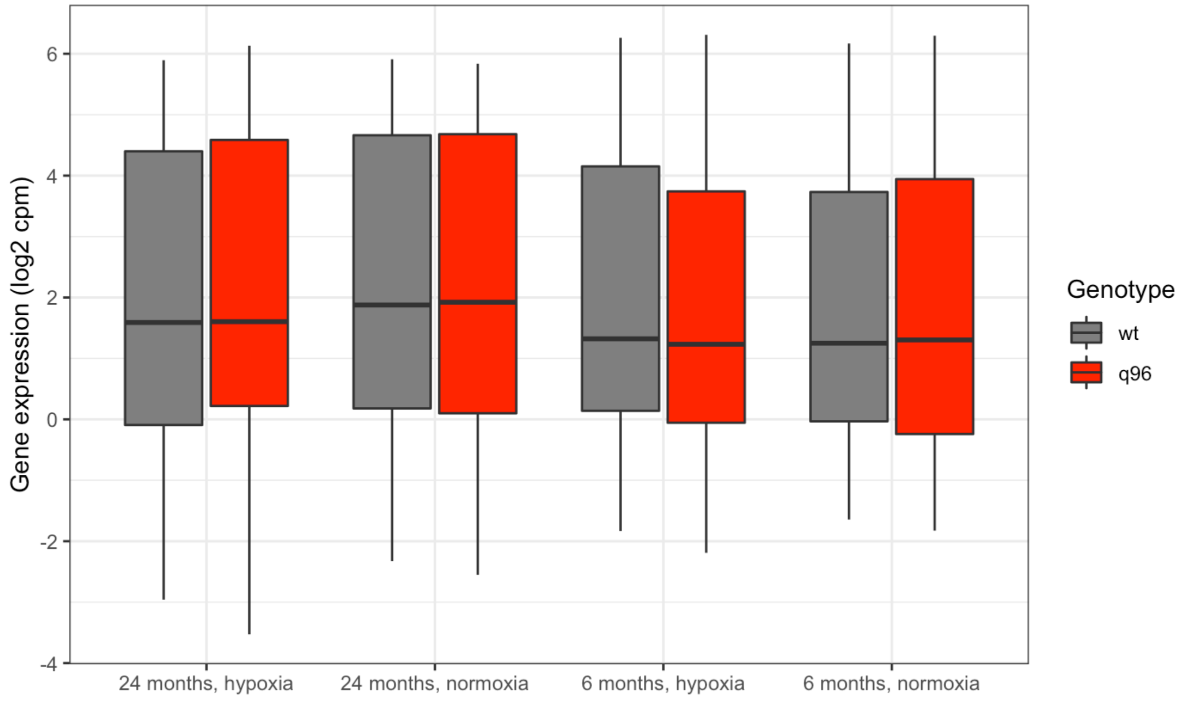


Astrocyte marker gene expression (61 genes)

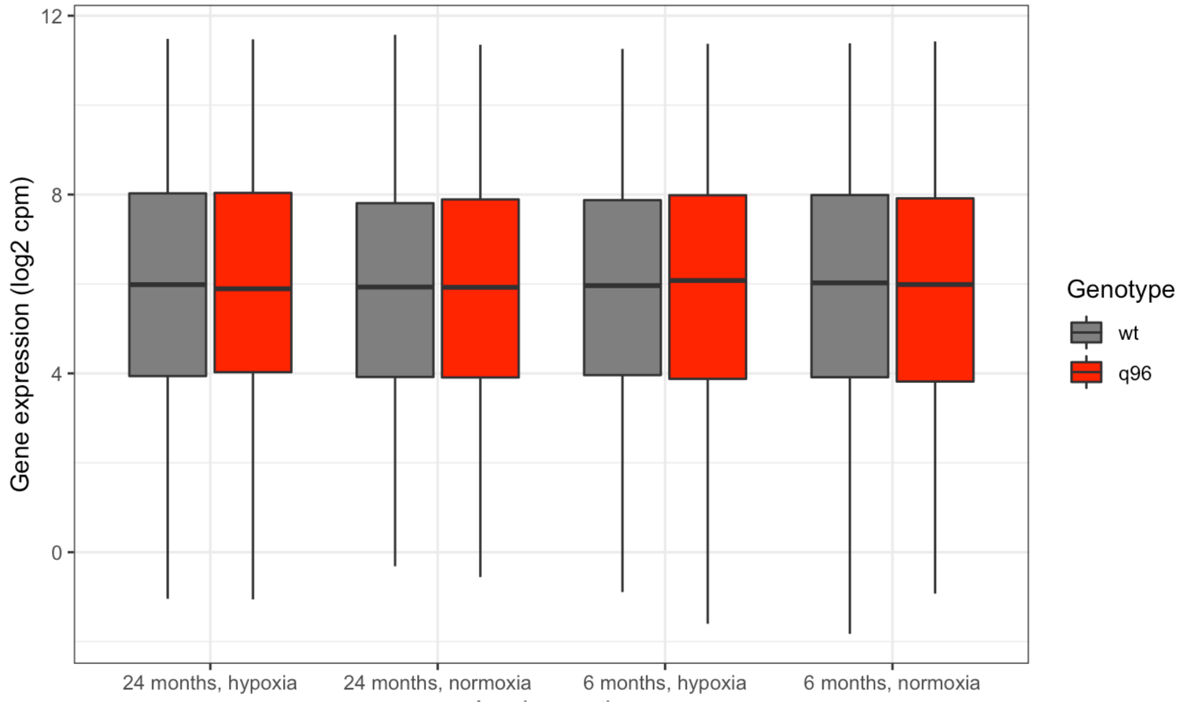

Neuron marker gene expression (87 genes)

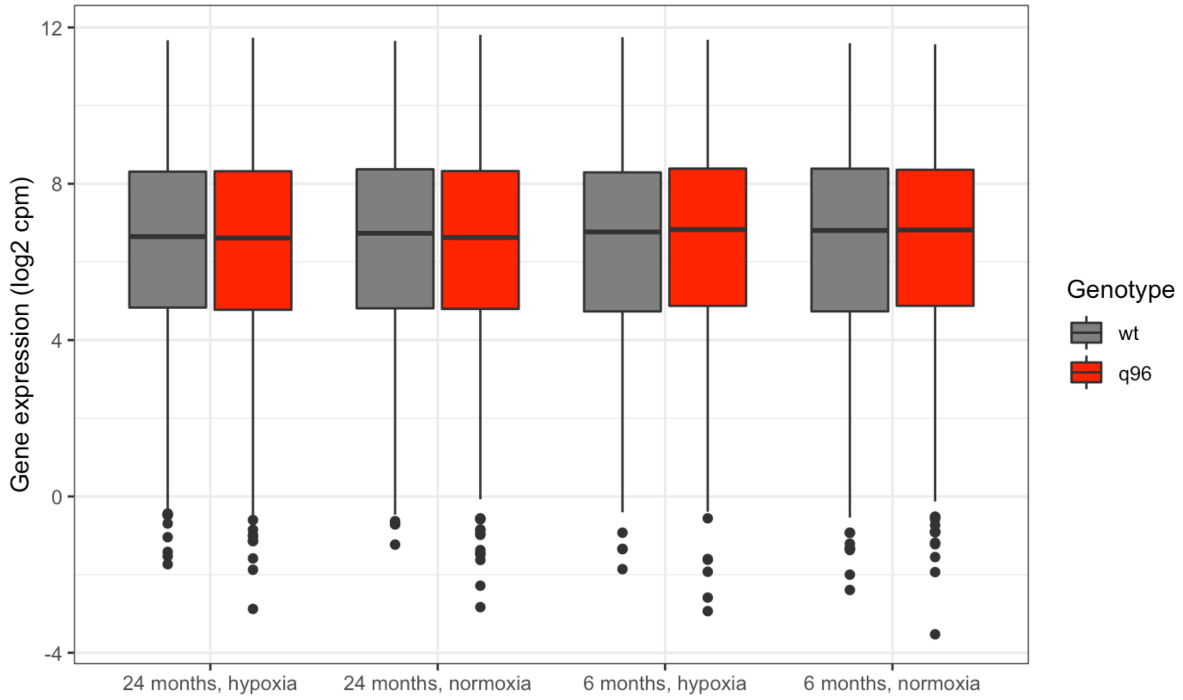


Oligodendrocyte marker gene expression (107 genes)

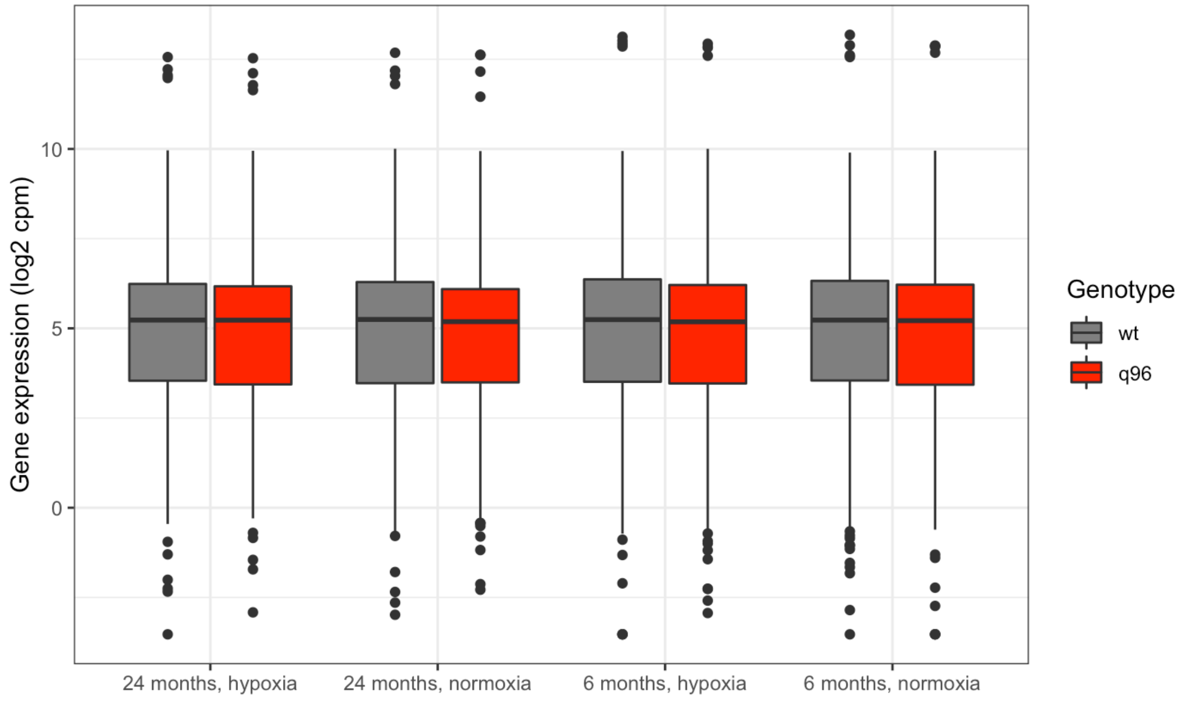
