## Supplementary Text 3 for "Iron Responsive Element (IRE)-mediated responses to iron dyshomeostasis in Alzheimer’s disease"

### Expression of previously characterized 3' and 5' IRE genes in the gene expression datasets analyzed in this paper

The IRP-IRE paradigm whereby binding of IRPs to 3' IREs causes stabilization of the transcript (and hence increased expression) and binding of IRPs to 5' IREs blocks translation was originally constructed through study of classic IRE-containing genes such as 3' IRE genes encoding ferroportin (SLC11A2 or DMT1, [1]) and transferrin receptor (TFRC, [2]), along with 5' IRE genes encoding ferritin light chain (FTL, [3]), ferritin heavy chain (FTH1, [4]), and Delta-aminolevulinate synthase 2 (ALAS2, [5]). Many other IRE-containing genes have since been identified in various species, some of which have functional IRE-like motifs in lieu of canonical IREs (please see Table 1 in ref. [6] and Table 1 in ref. [7] for summaries).

Below, we briefly discuss the expression of these previously characterised IRE genes in the datasets analysed in the main text of this paper, including the Caco-2 cell line dataset, 5XFAD mouse dataset, fAD-mutation-like zebrafish dataset, and Mayo Clinic human RNA-seq dataset. Boxplots showing the expression of these genes across all samples in each dataset are provided in **Supplementary Figure 1**. We note that some of the canonical IRE-containing genes in humans do not have IREs or IRE-like motifs in mouse or zebrafish and are hence not shown in the mouse/zebrafish figures. In addition, IRE genes which were not detected at sufficient levels in a RNA-seq dataset (at least 1 cpm across all samples in a dataset) are also not shown for the specific dataset in these figures.

**Supplementary Figure 1A** shows the expression of ferroportin (SLC11A2), transferrin receptor (TFRC), ferritin heavy and light chains (FTH1 and FTL respectively), and delta-aminolevulinate synthase 2 (ALAS2) in the Caco-2 cell line dataset. As the dataset consists of one cell type only, we expect the cells within each sample to respond in a similar way to iron dyshomeostasis, simplifying interpretation of these observations. Overall, the expression changes of FTL, FTH1 and TFRC under iron deficiency and iron overload conditions are consistent with the IRP-IRE binding paradigm. We note that ALAS2 (5' IRE) and SLC11A2 (DMT1, 3' IRE) do not match the paradigm completely. SLC11A2 shows decreased expression under iron overload, similar to TFRC, which might be expected as it has a 3’ IRE. However, we do not see a detectable increase in expression under iron deficiency as would be expected. Likewise, ALAS2 shows apparent stabilisation (or at least increased expression) under iron deficiency, despite having a 5’ IRE. This shows that variability exists even amongst canonical IRE genes; not all 3’ IRE genes can be expected to behave like TFRC, while not all 5’ IRE genes can be expected to behave like FTH1 or FTL.

**Supplementary Figure 1B** (temporal cortex) and **Supplementary Figure 1C** (cerebellum) shows the expression of several previously characterised human 3’ and 5’ IRE genes (see Table 1 in ref. [6] and Table 1 in [7]) in the Mayo Clinic RNA-seq dataset. In this dataset, temporal cortex and cerebellum tissue was sampled from the same post-mortem brains. The conditions represented in this dataset include healthy control brains, Alzheimer’s disease (AD), pathological aging (PA, defined as presence of amyloid pathology but not neurodegeneration), and progressive supranuclear palsy (PSP, a tauopathy without amyloid pathology). Mean age at death of samples in each condition is similar (see Methods in main text for details). **Supplementary Figure 1B** shows expression of these IRE genes in the temporal cortex while **Supplementary Figure 1C** shows expression of these IRE genes in the cerebellum. In the temporal cortex, it can be seen that several 3’ IRE genes (TFRC, TRAM1, SERTAD2, CDC14A, and CAV3) show significantly increased expression in AD temporal cortex tissue compared to controls. This increased expression is not seen in PA or PSP conditions relative to controls. Notably, expression of the TRAM1 3’ IRE gene is in the opposite direction in PA compared to AD. Overall, PA and PSP show mostly dissimilar IRE gene expression changes compared to AD, suggesting that the regulation of iron homeostasis through the IRP-IRE system differs between these neuropathological conditions. The increased expression of 3’ IRE genes in AD samples may suggest that the temporal cortex of AD brains may, overall, be in a state of ferrous iron deficiency relative to controls. However, this observation is tentative as there is a large amount of variation in the expression of these genes. IRE gene expression patterns in the cerebellum tissue differs from temporal cortex tissue in that the 3’ IRE gene SERTAD2 still shows increased expression in AD relative to control, but the other 3’ IRE genes (TFRC, TRAM1, CDC14A, CAV3) do not. In addition, the 3’ IRE gene CDC42BPA is increased in AD cerebellum but not in the temporal cortex. Overall, these observations suggest that gene expression varies across brain region and tissue type, and supports the known phenomenon of the cerebellum potentially being relatively less affected in AD compared to the temporal cortex. Notably, both of these 3’ IRE genes also show increased expression in PSP relative to controls but not in PA. The differences seen between IRE gene expression in AD and PA give initial support to the idea that amyloid pathology by itself is not sufficient to cause the iron dyshomeostasis seen in AD, at least at the gene expression level, and this effect is demonstrated more broadly across the predicted IRE gene sets in the main text.

**Supplementary Figure 1D** and **Supplementary Figure 1E** show the expression of previously characterised IRE genes in the 5XFAD mouse and fAD-mutation-like zebrafish respectively. Because these samples consist of many cell and tissue types (zebrafish samples are whole brains while mouse samples are cortex tissue), we would not expect the IRE expression differences to be as pronounced in these datasets compared to the Caco-2 cell line. In addition, the small sample sizes of these datasets compared to the human Mayo Clinic RNA-seq dataset likely do not give sufficient power to detect subtle differences when looking at individual IRE genes. Overall **Supplementary Figures 1D** and **1E** do not show clear differences in expression of the individual IRE genes between 5XFAD mice and their wild type siblings, and between fAD-mutation-like (*psen1*$^{Q96\_K97del/+}$) zebrafish and their wild type siblings, and from the expression of these genes alone, it is not clear whether the iron homeostasis state in these brains is closer to iron overload or iron deficiency. Using much larger gene sets in the main text of this paper, we are able to show more subtle, coordinated shifts in IRE gene expression which suggest disruptions to IRP-IRE system-mediated iron homeostasis at the gene expression level. However, further research deconvoluting the different cell types involved would be required to make an assessment of whether the brains of these animal models are in a state of iron deficiency or overload. This is beyond the scope of our method and we expand on potential ideas for addressing this limitation in the Discussion of the main text.
