## Supplementary Text 4 for "Iron Responsive Element (IRE)-mediated responses to iron dyshomeostasis in Alzheimer’s disease"

### Identifying 3’ IRE genes with altered stability

In the datasets examined in this paper, we noticed that expression of 3’ IRE-containing genes could be increased or decreased under various conditions. The current model of the IRP/IRE-system proposes that 3’ IRE-containing transcripts are stabilized and hence increased in expression under ferrous iron deficiency. However, even in the cultured cell line dataset subjected to iron deficiency, we saw that a large proportion of 3’ IRE-containing genes appeared to be decreased in expression, which does not support the idea that they are stabilized. These findings are difficult to reconcile with the current paradigm and suggest that our knowledge on the IRP/IRE system may be incomplete, especially for less well-characterized IREs.

We attempted to further explore stability changes by identifying genes whose transcripts showed differential stability between conditions in the fAD-like zebrafish dataset. We compared changes in their spliced and unspliced (but polyA-containing) transcripts and defined the null (no stabilization of transcript) and alternate (stabilization of transcript) hypotheses for each gene as follows, where *s* and *u* refer to the spliced and unspliced versions of a particular gene:

$$H_0: \ \mathrm{logFC_s} = \mathrm{logFC_u}$$
$$H_a: \ \mathrm{logFC_s} \neq \mathrm{logFC_u}$$

In this way, $\mathrm{logFC_s} > \mathrm{logFC_u}$ would be interpreted as an increase in stability while $\mathrm{logFC_s} < \mathrm{logFC_u}$ would be interpreted as a decrease in stability. Because the distributions $\mathrm{logFC_s}$ and $\mathrm{logFC_u}$ are likely to have unequal variances, we used Welch’s t-test (t-test of unequal variances) to determine whether stability was significantly different.

The full stability analysis results for all genes (3’ IRE, 5’ IRE, and no IRE) for each comparison are provided in **Supplementary Table 5**. Interestingly, there were 3’ IRE genes with significant increases and decreases in stability in all comparisons, some of which are shown in **Supplementary Figure 7.**
