## Supplementary Figures for "Iron Responsive Element (IRE)-mediated responses to iron dyshomeostasis in Alzheimer’s disease"

### Supplementary Figure 1.

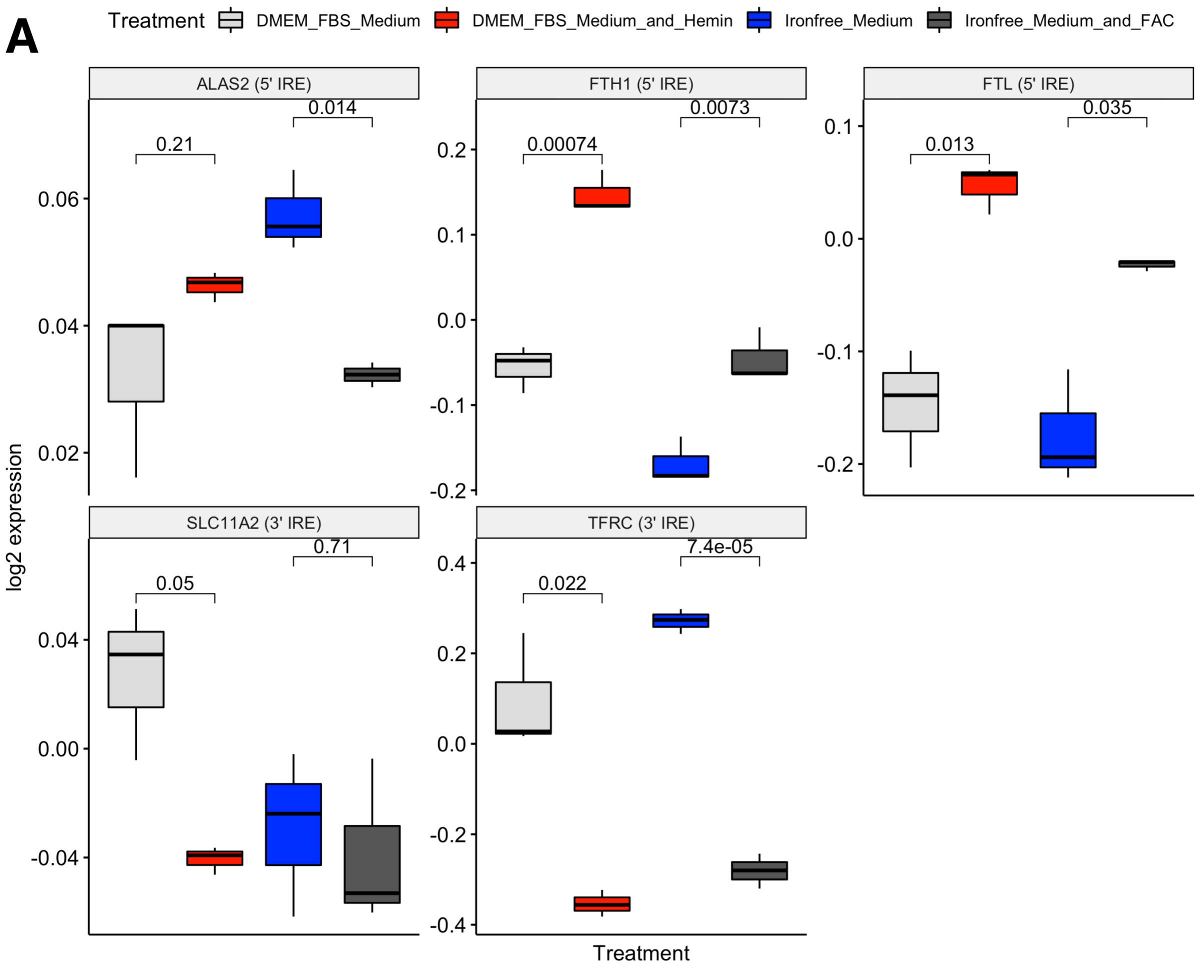


**Supplementary Figure 1A. Boxplots showing gene expression (log2 RMA-normalised intensity) of previously characterised 3’ and 5’ IRE-containing genes across all samples in the Caco-2 cell line dataset.** Numbers above boxplots represent *p*-values from *t*-tests to test if there is a difference in mean expression during iron overload (“DMEM FBS Medium and Hemin” vs. “DMEM FBS Medium”) or iron deficiency (“Ironfree Medium” vs. “Iron Free Medium and FAC”). Significant differences are defined as $p < 0.05$.

**B**

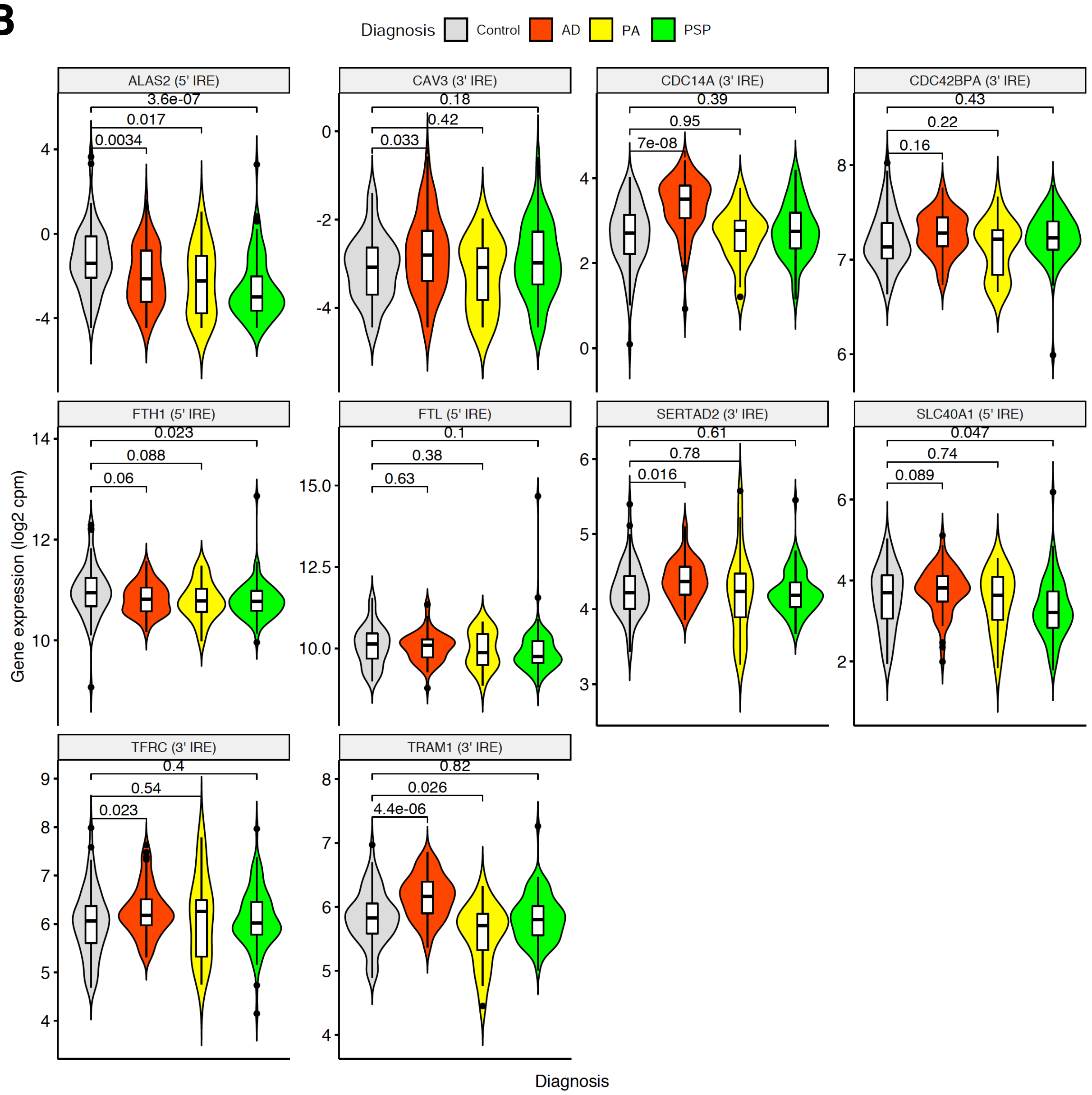


**Supplementary Figure 1B. Boxplots and Violin Plots showing gene expression (log2 TMM-normalised cpm) of previously characterised 3' and 5' IRE-containing genes detected at sufficient levels (> 1 cpm) across temporal cortex samples of the Mayo Clinic RNA-seq dataset.** Numbers above boxplots represent *p*-values from *t*-tests to test if there is a difference in mean expression in Alzheimer's disease (AD), pathological aging (PA), or progressive supranuclear palsy (PSP) compared to control samples. Temporal cortex and cerebellum tissues were taken from the same brains. Significant differences are defined as $p < 0.05$.

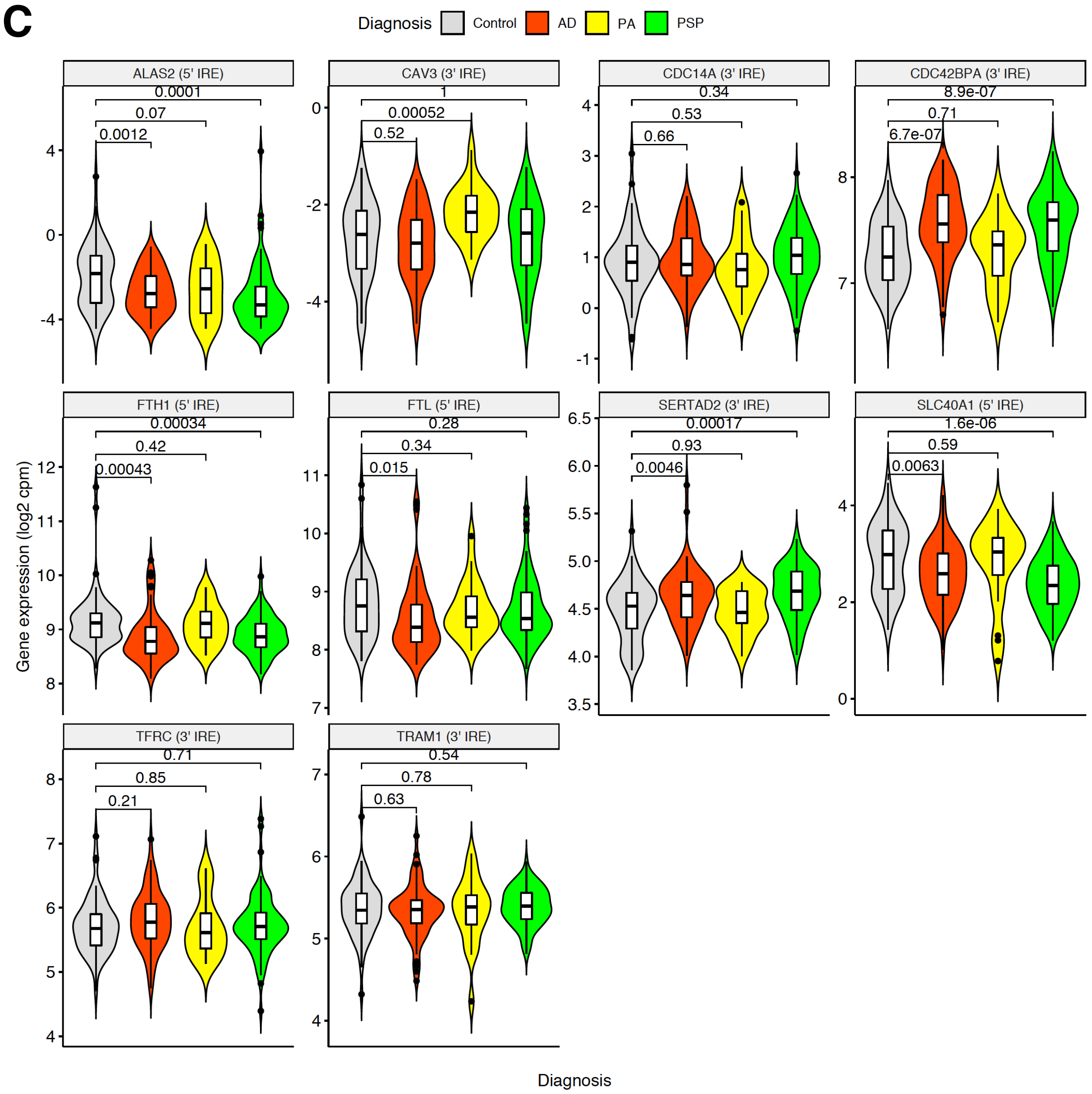


**Supplementary Figure 1C. Boxplots and Violin Plots showing gene expression (log2 TMM-normalised cpm) of previously characterised 3' and 5' IRE-containing genes detected at sufficient levels (> 1 cpm) across cerebellum samples in the Mayo Clinic RNA-seq dataset.** Numbers above boxplots represent *p*-values from *t*-tests to test if there is a difference in mean expression in Alzheimer's disease (AD), pathological aging (PA), or progressive supranuclear palsy (PSP) compared to control samples. Temporal cortex and cerebellum tissues were taken from the same brains. Significant differences are defined as $p < 0.05$.

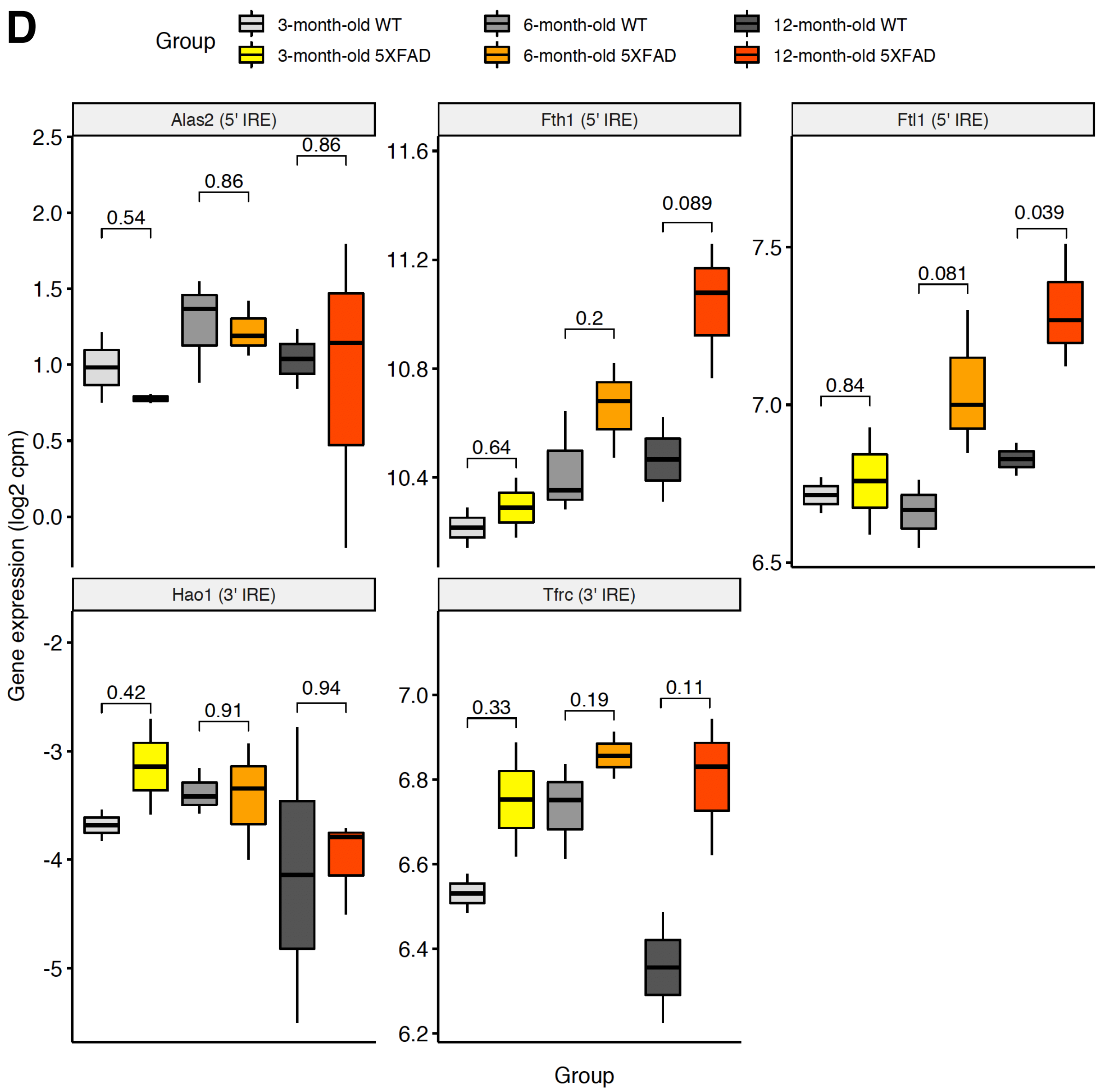


**Supplementary Figure 1D. Boxplots showing gene expression (log2 TMM-normalised cpm) of previously characterised 3' and 5' IRE-containing genes detected at sufficient levels (> 1 cpm) across 5XFAD and wild type (WT) mouse cortex samples.** Numbers above boxplots represent *p*-values from *t*-tests to test if there is a difference in mean expression in 5XFAD mouse cortex samples compared to wild type siblings. Significant differences are defined as $p < 0.05$.

**E**

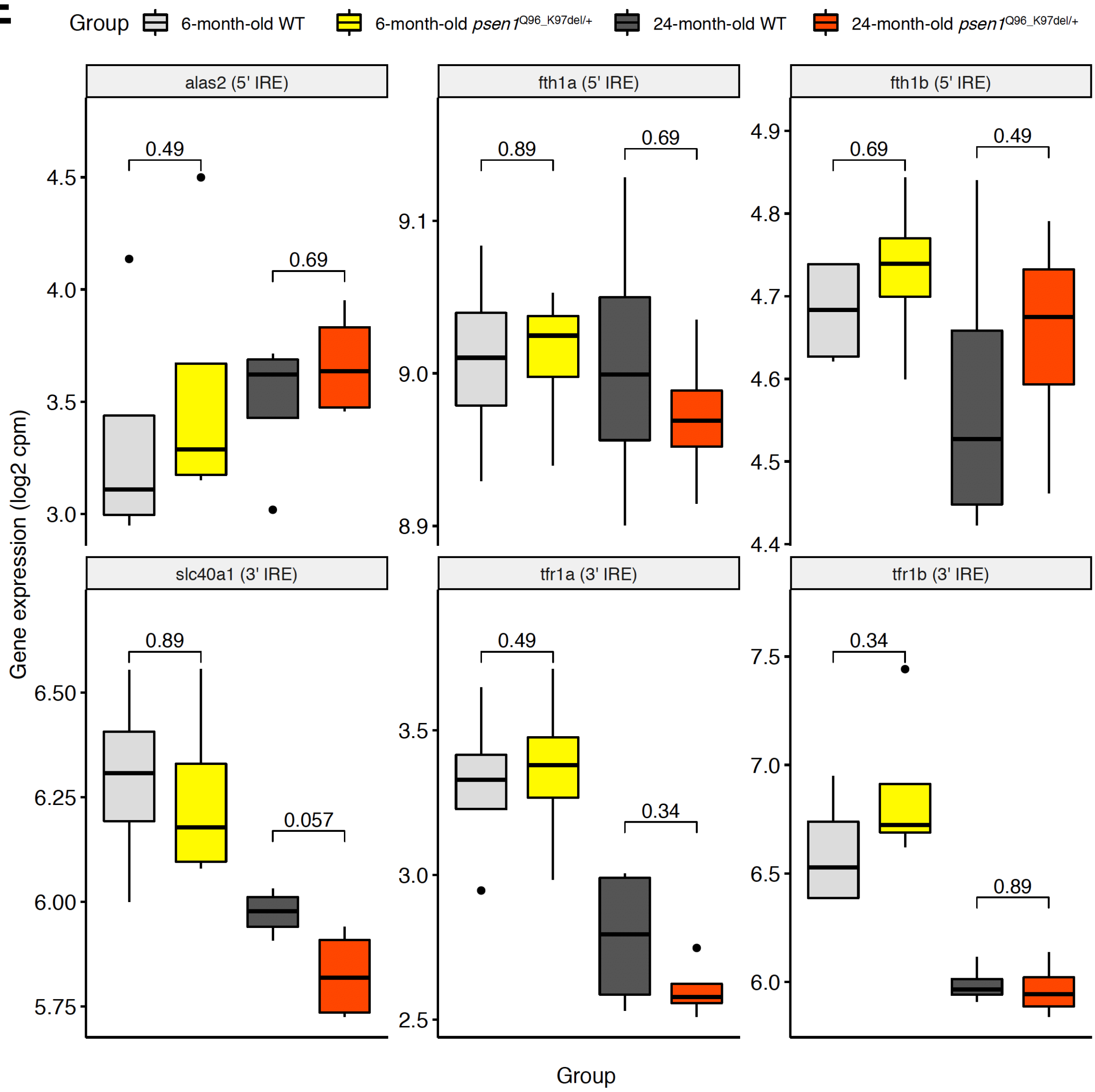


**Supplementary Figure 1E. Boxplots showing gene expression (log2 TMM-normalised cpm) of previously characterised 3’ and 5’ IRE-containing genes detected at sufficient levels (> 1 cpm) across fAD-mutation-like *psen1*$^{Q96\_K97del/+}$ and *psen1*$^{+/+}$ (WT) zebrafish whole brains.** Only the normoxia samples are shown in this figure. Numbers above boxplots represent *p*-values from *t*-tests to test if there is a difference in mean expression in *psen1*$^{Q96\_K97del/+}$ whole brains compared to wild type *psen1*$^{+/+}$ siblings. Significant differences are defined as $p < 0.05$.

**Supplementary Figure 2.**

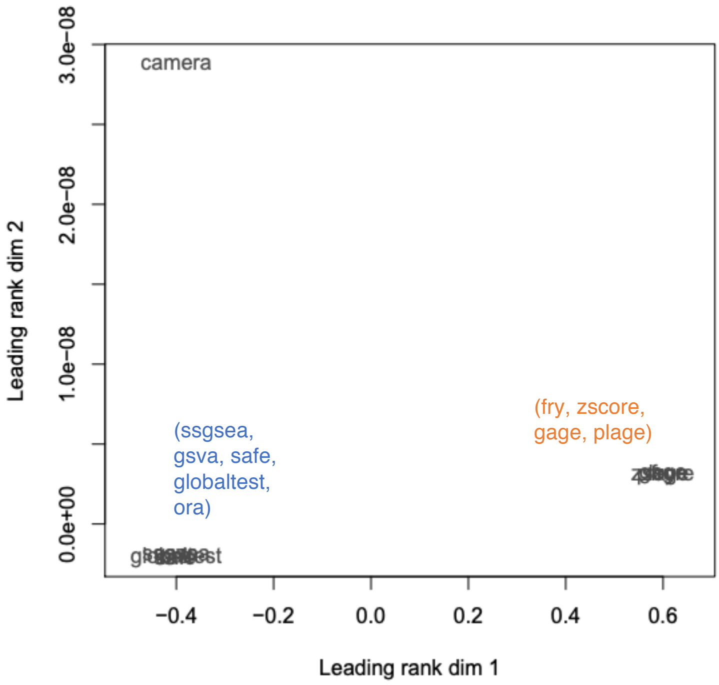


**Principal Component Analysis of results from different gene set testing methods.**

We used EGSEA (v.1.10.1) with default settings with R objects (design, contrasts, voom) from *limma* analysis of the fAD-like zebrafish dataset (n=32). EGSEA was run with the following methods: camera (limma v.3.38.3), safe (safe v.3.22.0), gage (gage v.2.32.1), plage (GSVA v.1.30.0), zscore (GSVA v.1.30.0), gsva (GSVA:1.30.0), ssgsea (GSVA v.1.30.0), globaltest (globaltest v.5.36.0), ora (stats v.3.5.2), fry (limma v.3.38.3). The Principal Component Analysis plot shows the relative similarity of the results obtained from running different methods, overall revealing three main groupings. To minimise the chance of *p*-value inflation from combining multiple methods that give essentially the same results, we decided to use one representative method from each group in our final analyses: camera, fry, and fgsea. Although fgsea is not a method included in EGSEA, we chose to include it as it implements the classic GSEA algorithm and also includes information about leading-edge genes which we make use of in our analyses.

### Supplementary Figure 3.

**A**

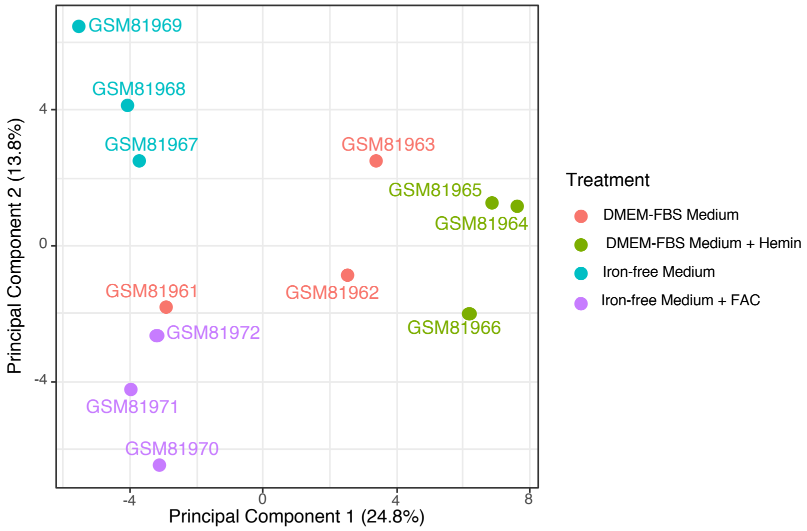


**B**

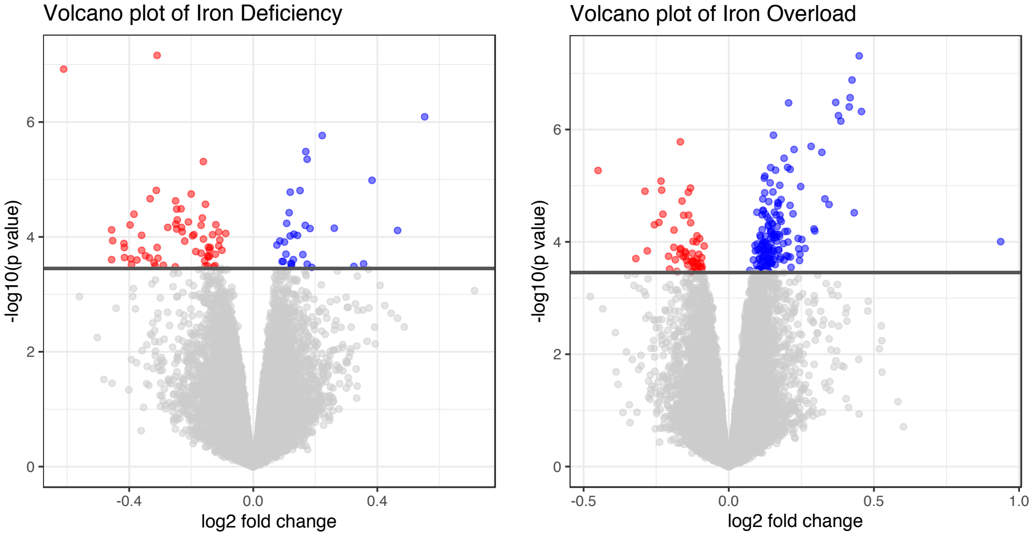


A. **Principal Component Analysis plot of gene expression in the cultured Caco-2 cell line dataset.** The plot uses pre-processed and normalised expression values for 22,153 probesets across all samples.

B. **Volcano plots indicating differential gene expression due to iron overload (DMEM-FBS Medium + Hemin vs. DMEM-FBS Medium) and iron deficiency (Iron-free medium vs. Iron-free medium + FAC) treatments in the Caco-2 cell line dataset.** Horizontal lines indicate an FDR-adjusted *p*-value of 0.05, with all genes with FDR-adjusted *p*-value < 0.05 considered significantly differentially expressed (DE). In the iron deficiency treatment, this resulted in 96 significantly DE genes (65 down, 31 up), while in the iron overload treatment there were 212 significantly DE genes (55 down, 157 up).

### Supplementary Figure 4

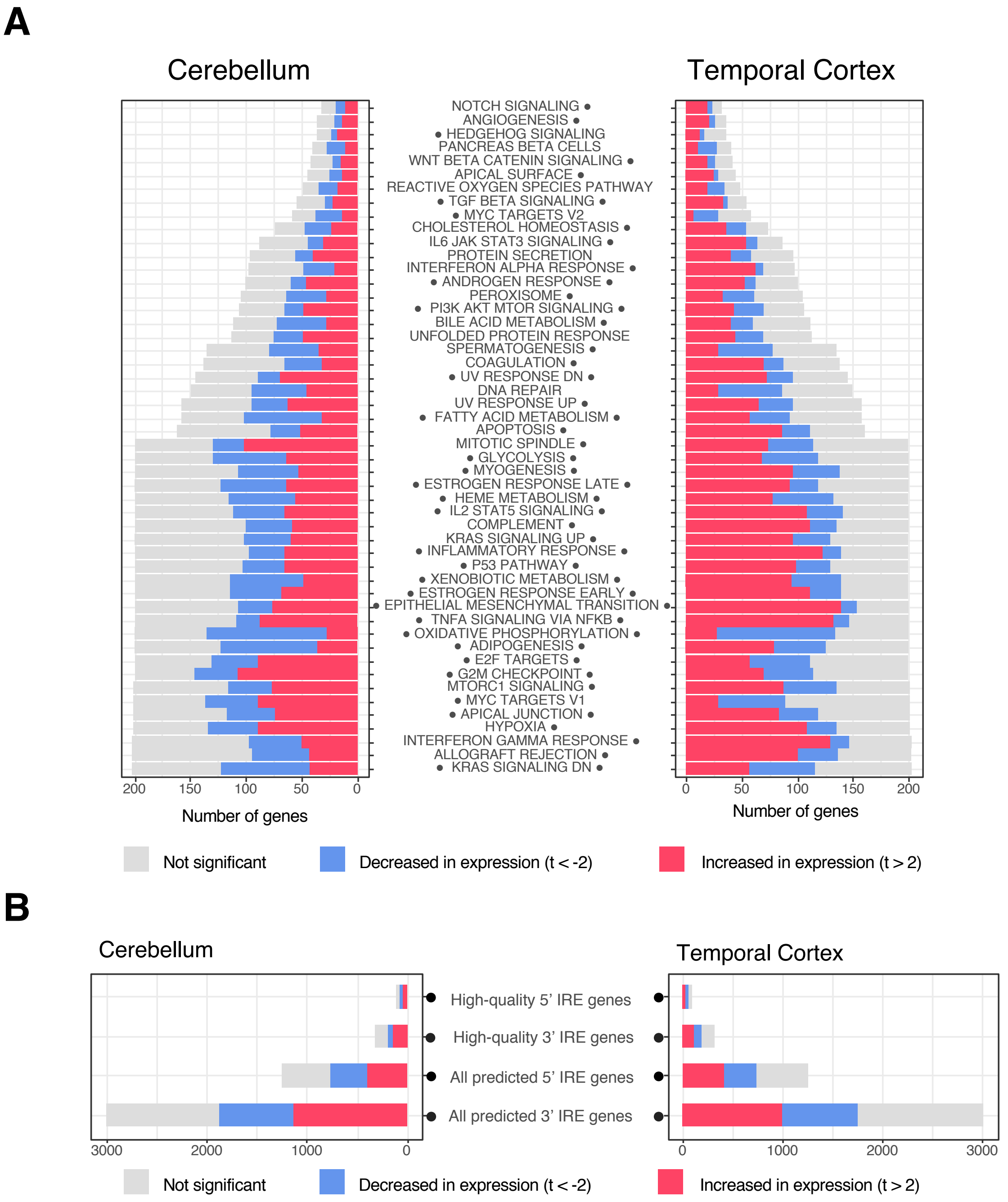


**Gene set enrichment testing results for the AD vs. control comparisons in cerebellum and temporal cortex.**

**A. Enrichment results for MSigDB Hallmark gene sets**
**B. Enrichment results for human IRE gene sets**

Dots to either the left or right side of the gene set name indicate that the gene set is significantly enriched in either the "AD vs. control with cerebellum tissue" or "AD vs. control with temporal cortex tissue" comparisons respectively (Bonferroni-adjusted *p*-value < 0.05).

**Supplementary Figure 5.**

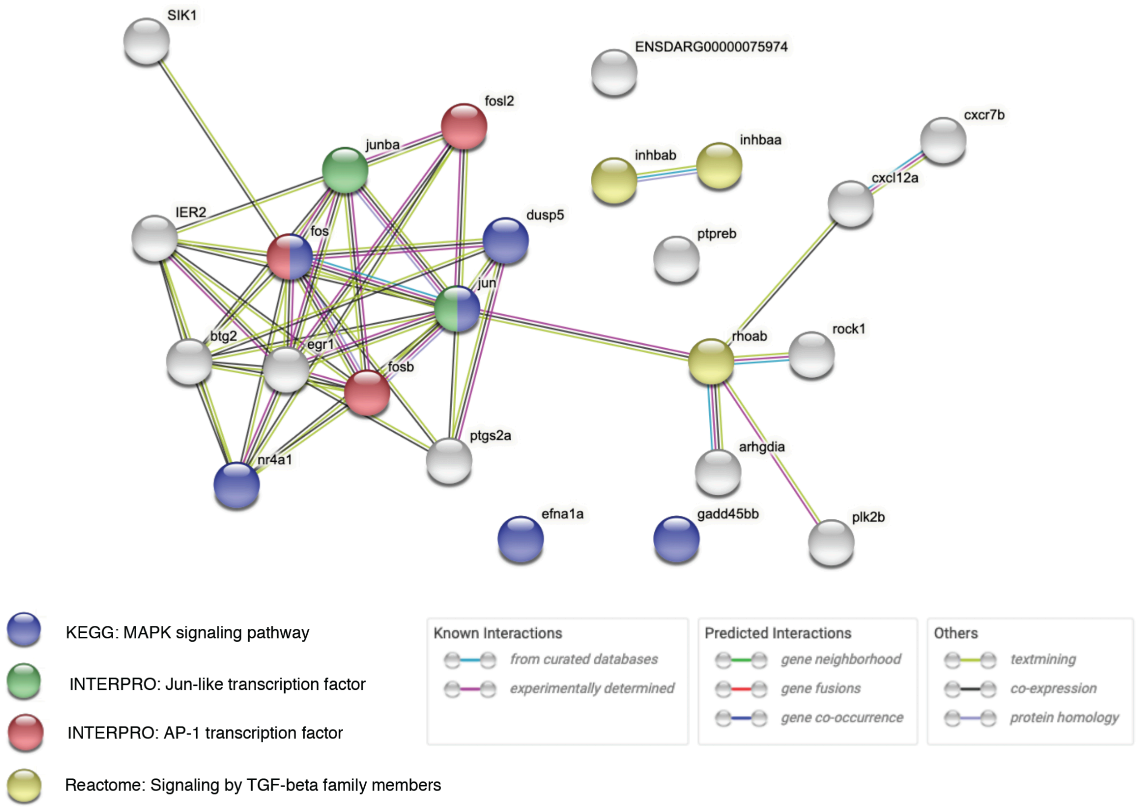


**STRINGR protein-protein interaction network plot between the 19 shared leading-edge genes between the *psen1*$^{Q96\_K97del/+}$ and *psen1*$^{+/+}$ comparisons at 6 and 24-months-old**. Nodes are named with gene symbols and edges indicate a known or predicted protein-protein interaction. Coloured nodes indicate genes contributing to significant over-representation of particular gene sets (FDR-adjusted *p*-value < 0.05).

### Supplementary Figure 6.

**A** 3’ IRE genes 5’ IRE genes

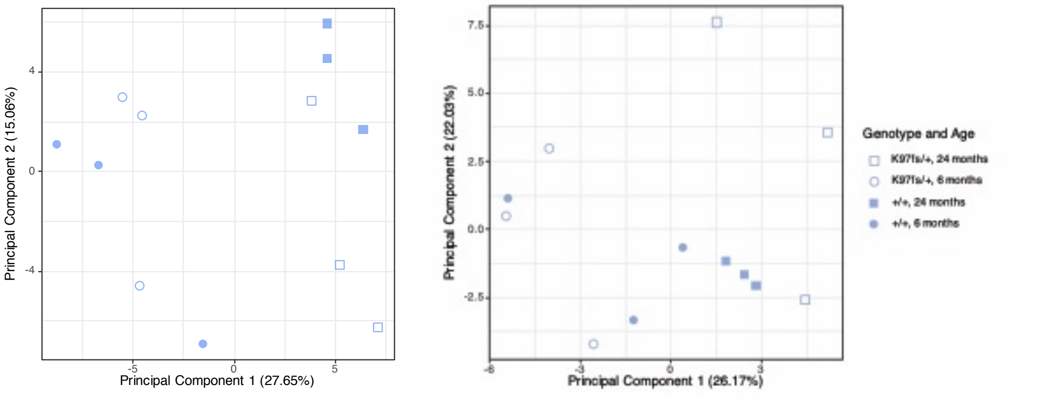


**B** All genes (18,296 genes)

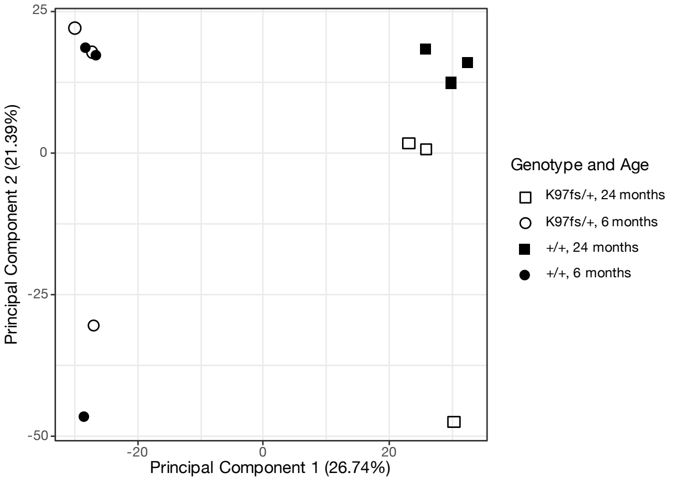


**Principal Component Analysis plots demonstrating the minimal association between IRE gene expression and the *psen1*$^{K97fs/+}$ mutant genotype.**

A. **Principal Component Analysis plots using expression of the zebrafish “all predicted 3’ IRE genes” and “all predicted 5’ IRE genes” gene sets.**

B. **Principal Component Analysis plot of all genes detected in the dataset.** Unlike for the fAD-like *psen1*$^{Q96\_K97del/+}$ mutant, the non fAD-like *psen1*$^{K97fs/+}$ mutant does not show significant changes in gene expression of 3’ and 5’ IRE genes.

### Supplementary Figure 7.

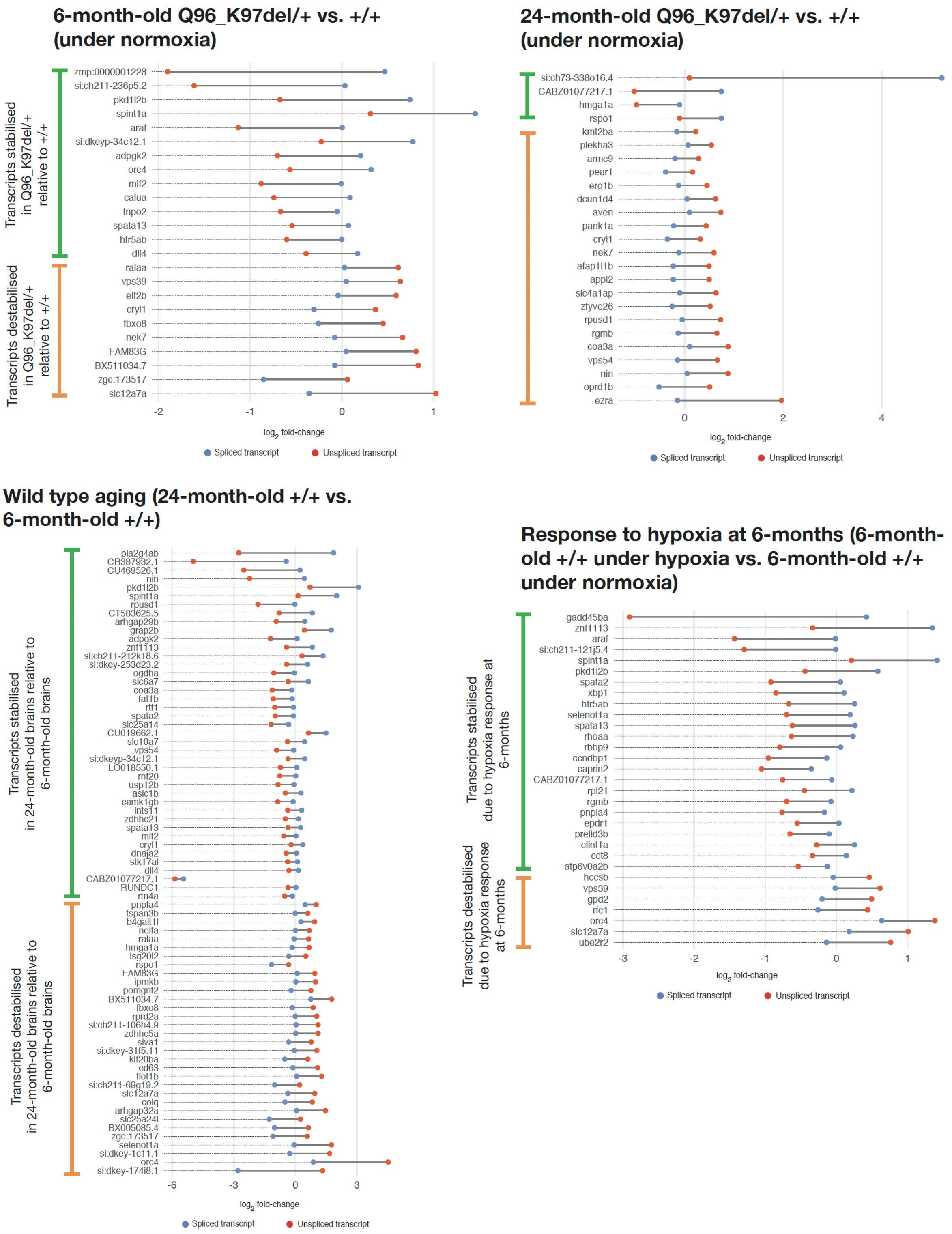


**Predicted 3' IRE-containing transcripts with significant differences in stability between conditions in the fAD-like zebrafish dataset.** Transcripts are defined to be stabilised if the log2 fold-change of the spliced transcript is significantly greater than the log2 fold-change of the unspliced transcript. Likewise, transcripts are defined to be destabilised if the log2 fold-change of the spliced transcript is significantly less than the log2 fold-change of the unspliced transcript. See Methods for details. All tests were done using Welch's t-test with FDR-adjusted *p*-value < 0.05.

**Supplementary Figure 8.**

Microglia marker gene expression (20 genes)

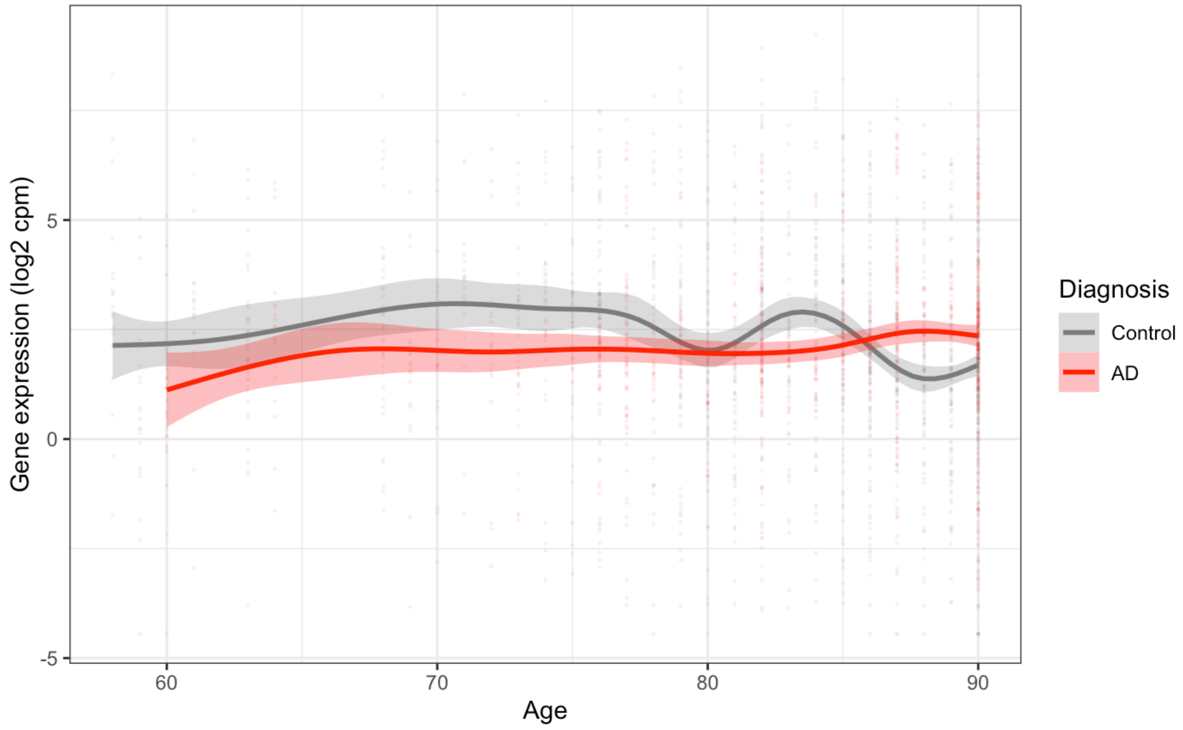


Astrocyte marker gene expression (44 genes)

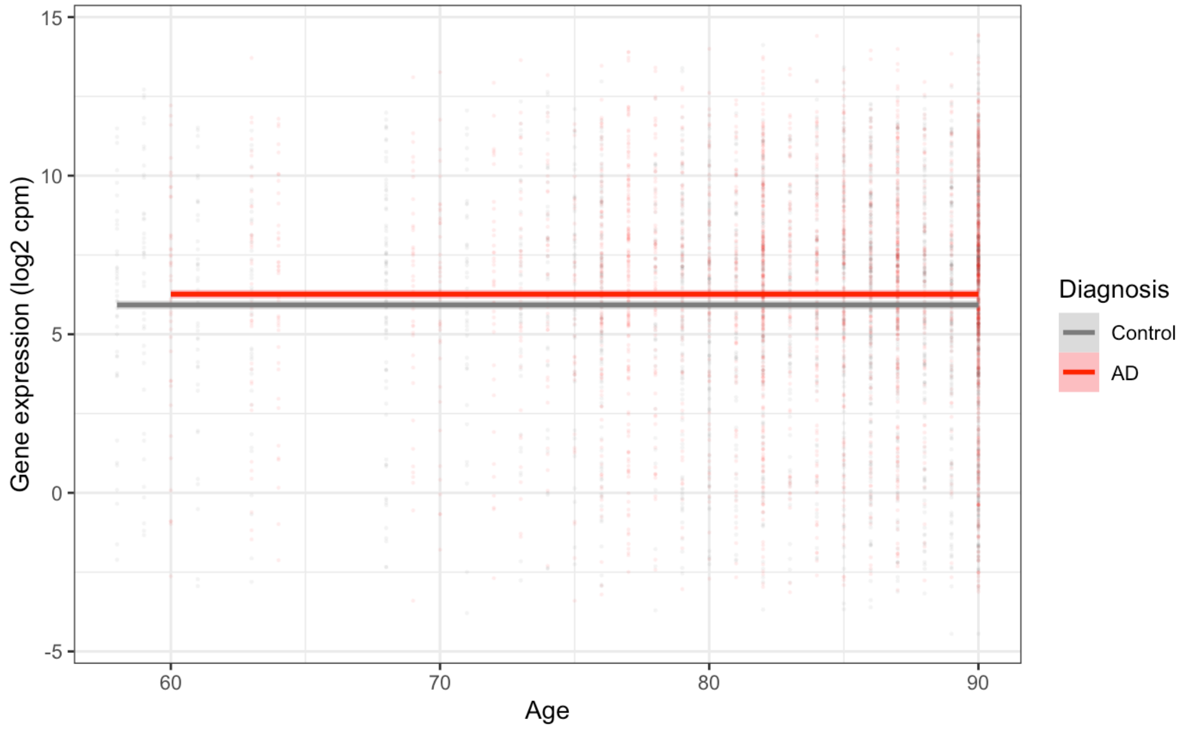

Neuron marker gene expression (74 genes)

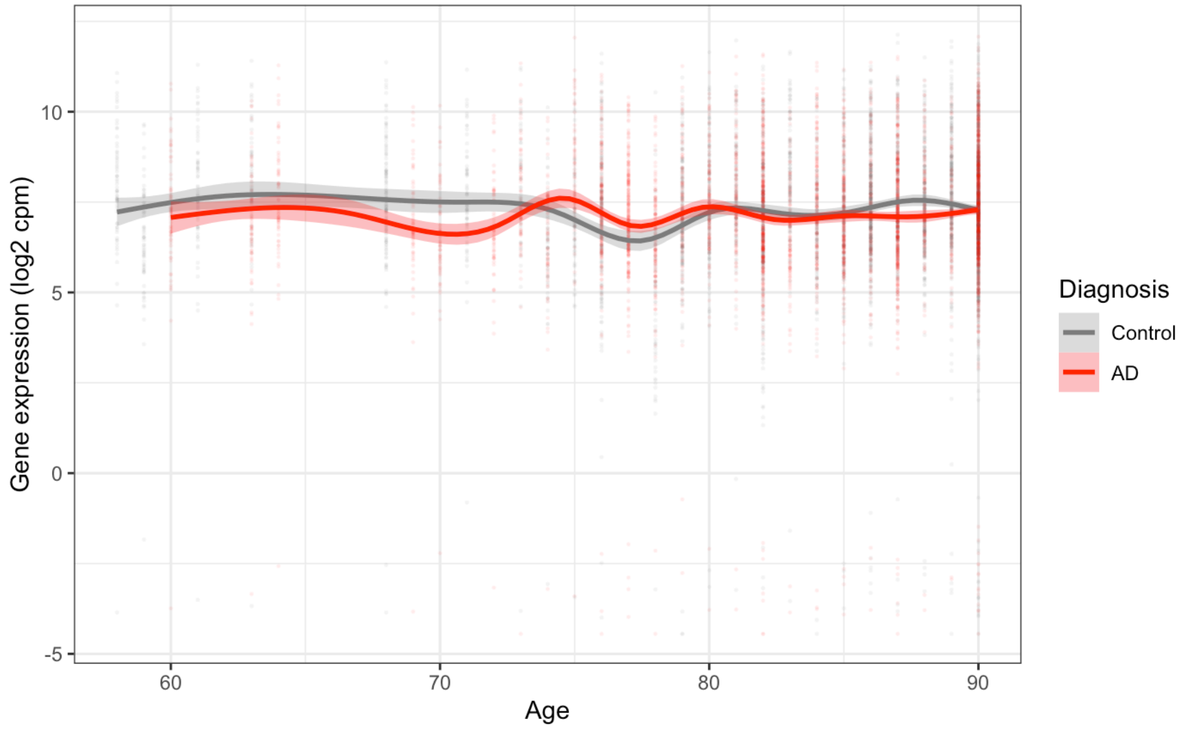


Oligodendrocyte marker gene expression (83 genes)

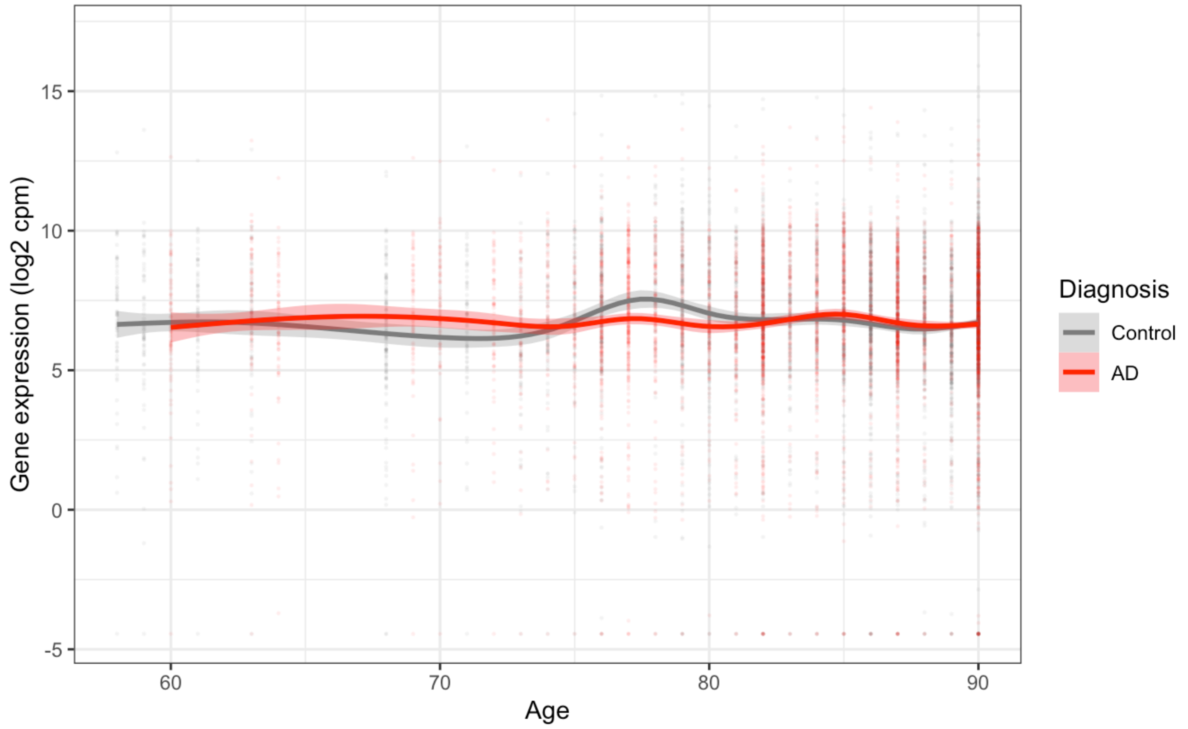


**Age-dependent expression of neural marker genes (microglia, astrocyte, neuron, oligodendrocyte) in human Mayo Clinic RNA-seq dataset analysed.**

There is a possibility that changes in gene expression between may reflect changes in the proportions of cell types rather than changes in expression in each cell. For example, less neurons might be expected during neurodegeneration. To explore the possible changes in the proportions of cell types, we looked in marker genes of three common neural cell types: astrocytes, neurons, microglia, and oligodendrocytes.

The marker genes for astrocytes, neurons, and oligodendrocytes were derived from gene sets by Lein et al. (2007) and are based on studies in mice (available on MSigDB). The marker genes for microglia were derived from Bonham et al. (2019) and based on studies in human and mouse. All gene IDs were converted to human Ensembl IDs where necessary using BioMart.

Error bars represent the 95% confidence interval. Overall, while we see differences in expression, there does not appear to be an overall shift / overall difference between AD vs. control for oligodendrocyte, neuron and microglial marker gene expression. For the astrocyte markers, AD brains appear to systemically have higher expression of these genes.

**Supplementary Figure 9.**

Microglial marker gene expression (22 genes)

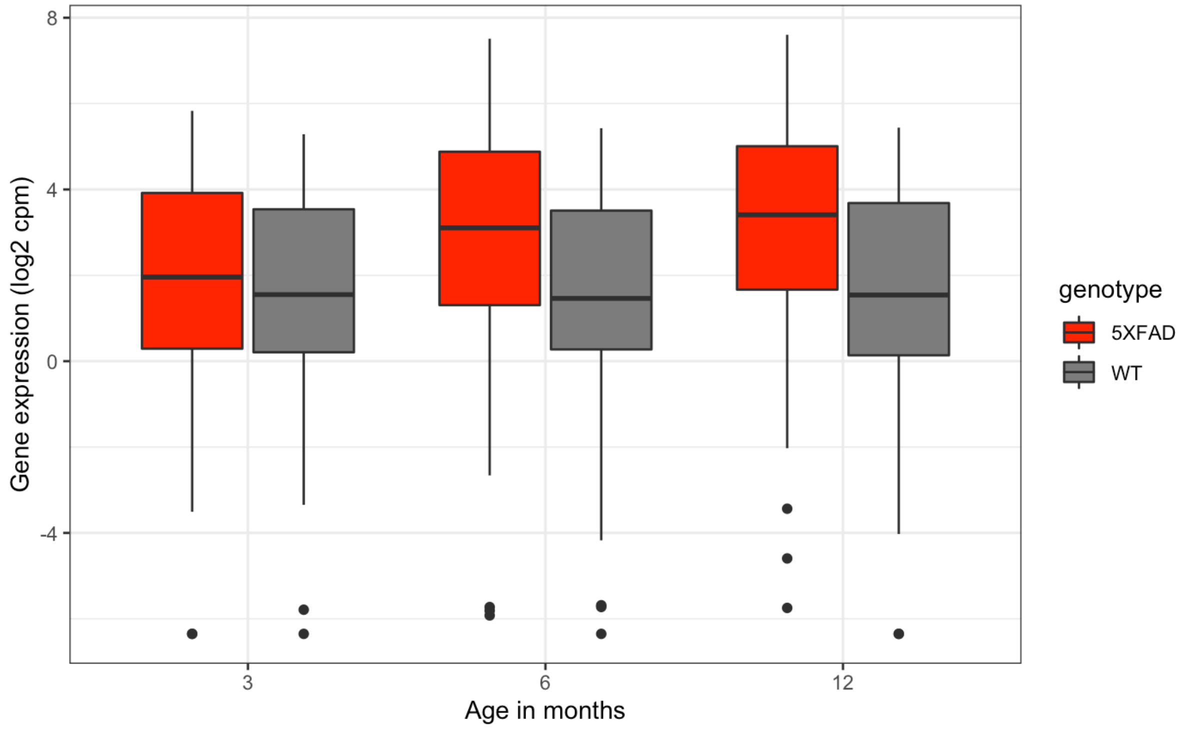


Astrocyte marker gene expression (51 genes)

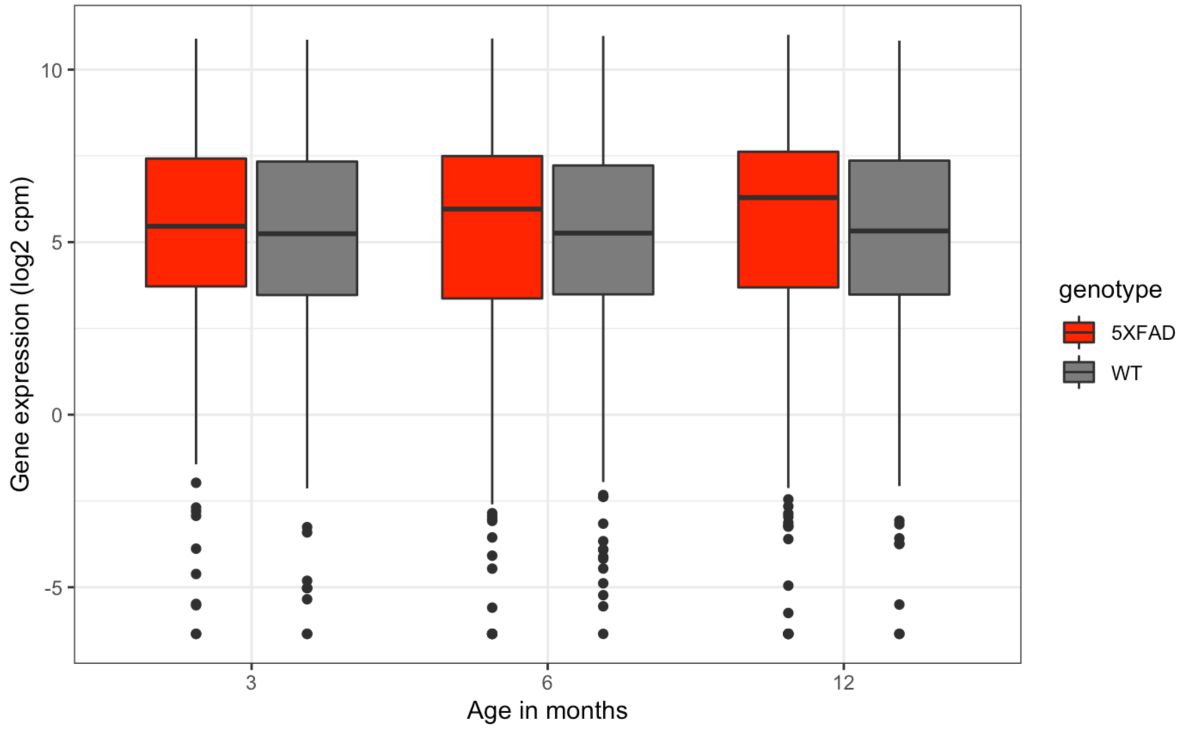

Neuron marker gene expression (67 genes)

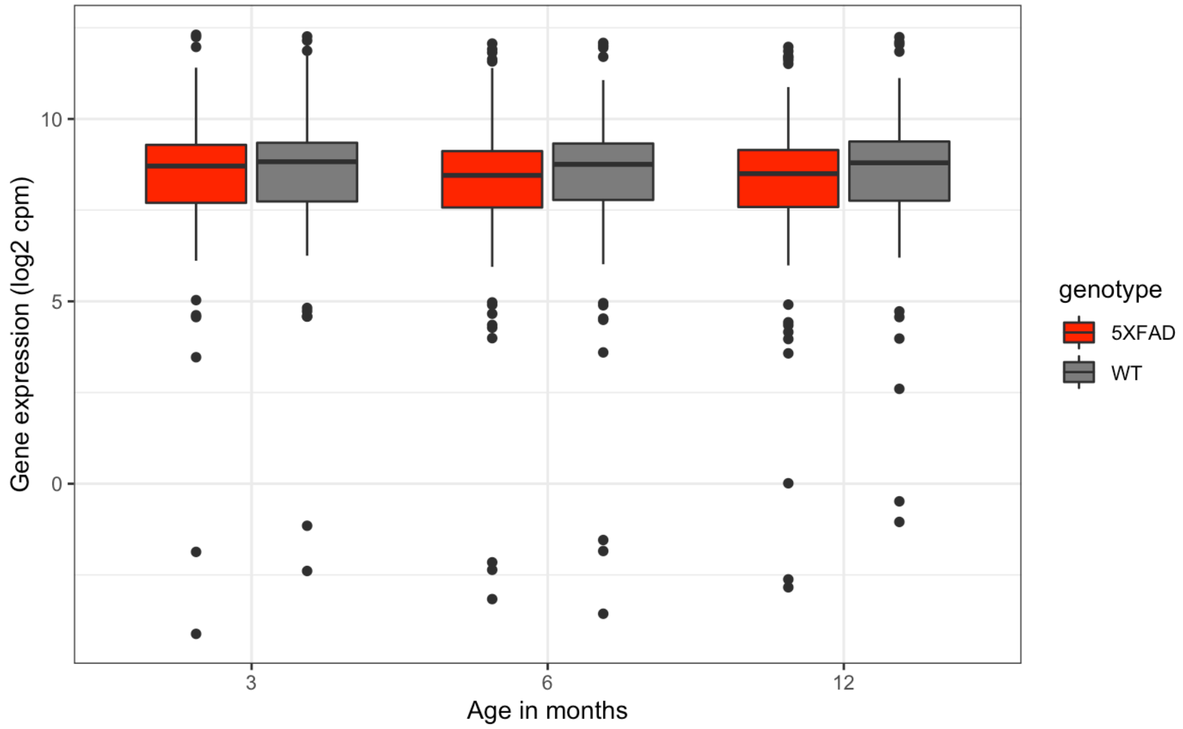


Oligodendrocyte marker gene expression (77 genes)

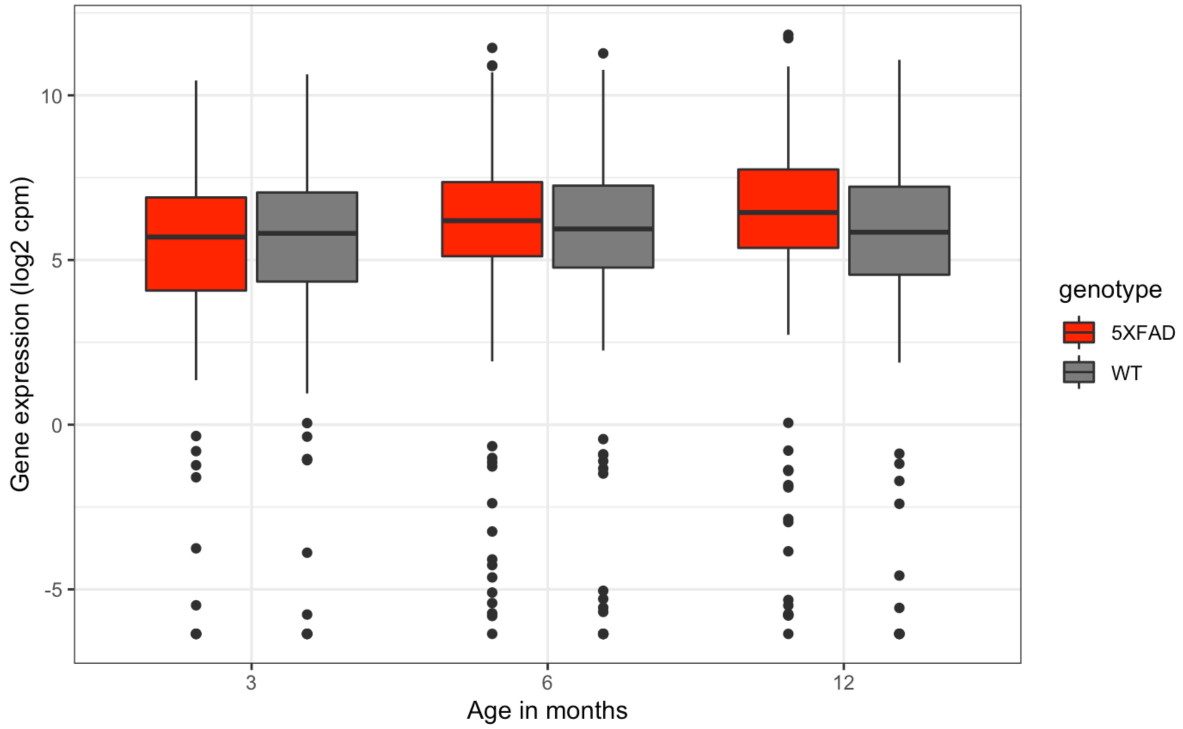


**Age-dependent expression of neural marker genes (microglia, astrocyte, neuron, oligodendrocyte) in 5XFAD mouse cortex RNA-seq dataset.**

There is a possibility that changes in gene expression between may reflect changes in the proportions of cell types rather than changes in expression in each cell. For example, less neurons might be expected during neurodegeneration. To explore the possible changes in the proportions of cell types, we looked in marker genes of three common neural cell types: astrocytes, neurons, microglia, and oligodendrocytes.

The marker genes for astrocytes, neurons, and oligodendrocytes were derived from gene sets by Lein et al. (2007) and are based on studies in mice (available on MSigDB). The marker genes for microglia were derived from Bonham et al. (2019) and based on studies in human and mouse. All gene IDs were converted to mouse Ensembl IDs using BioMart.

Overall, expression of these neural marker genes appears to be stable across different conditions and ages for neuron and oligodendrocyte marker genes. microglial and astrocyte marker gene expression appear to show age-dependent increased expression in the 5XFAD brains.

**Supplementary Figure 10.**

<u>Microglial marker gene expression (16 genes)</u>

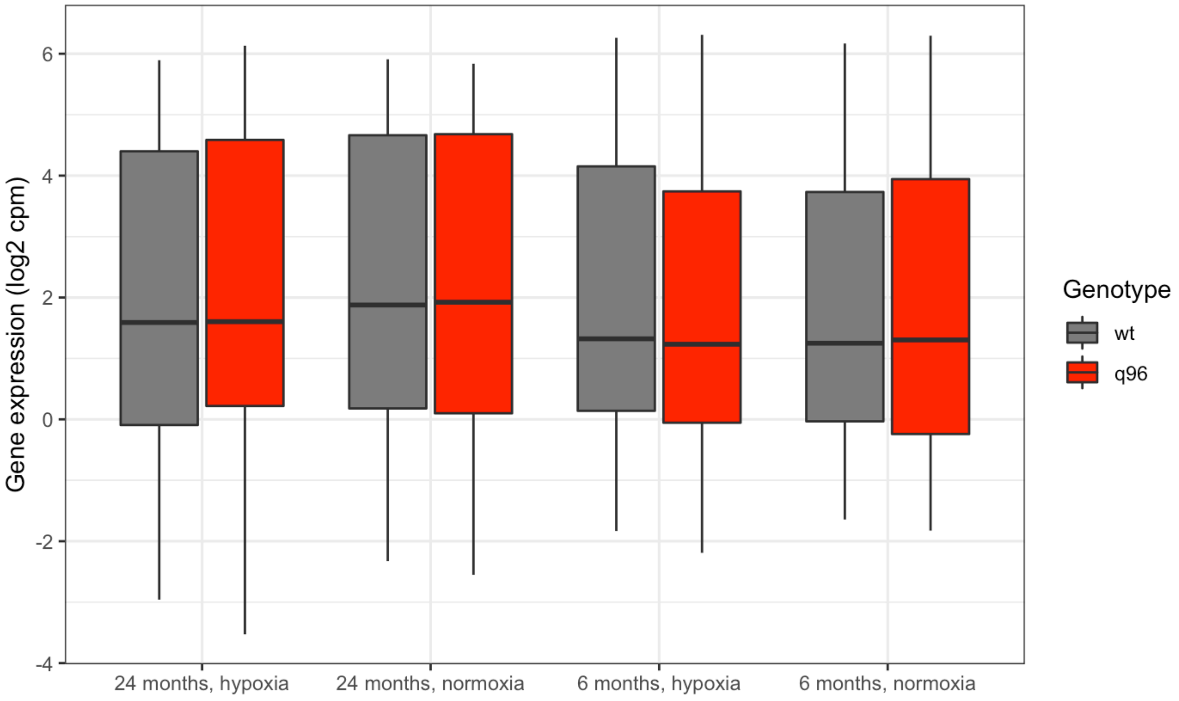


<u>Astrocyte marker gene expression (61 genes)</u>

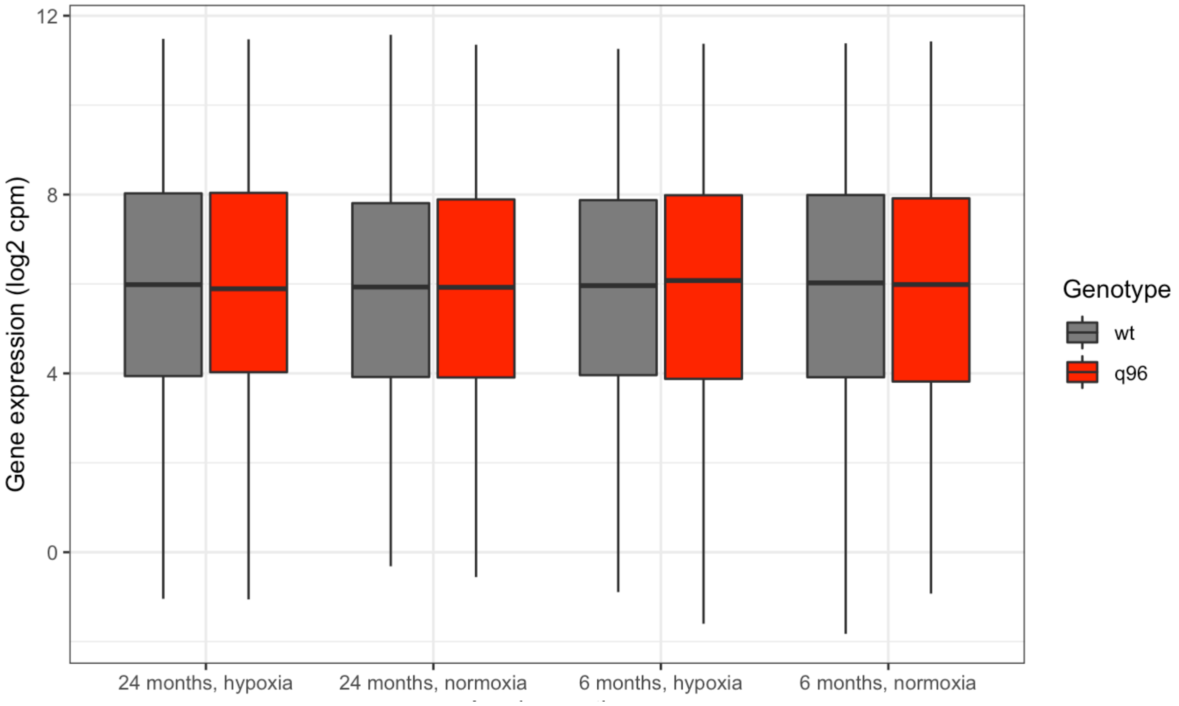

Neuron marker gene expression (87 genes)

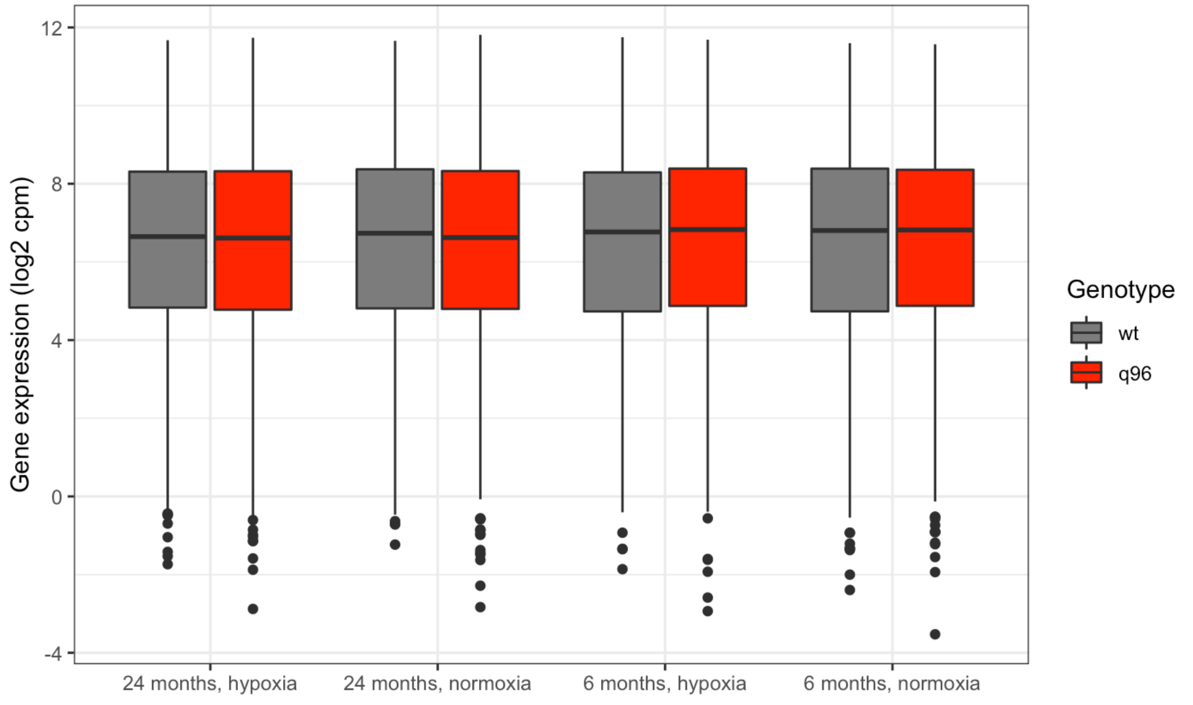


Oligodendrocyte marker gene expression (107 genes)

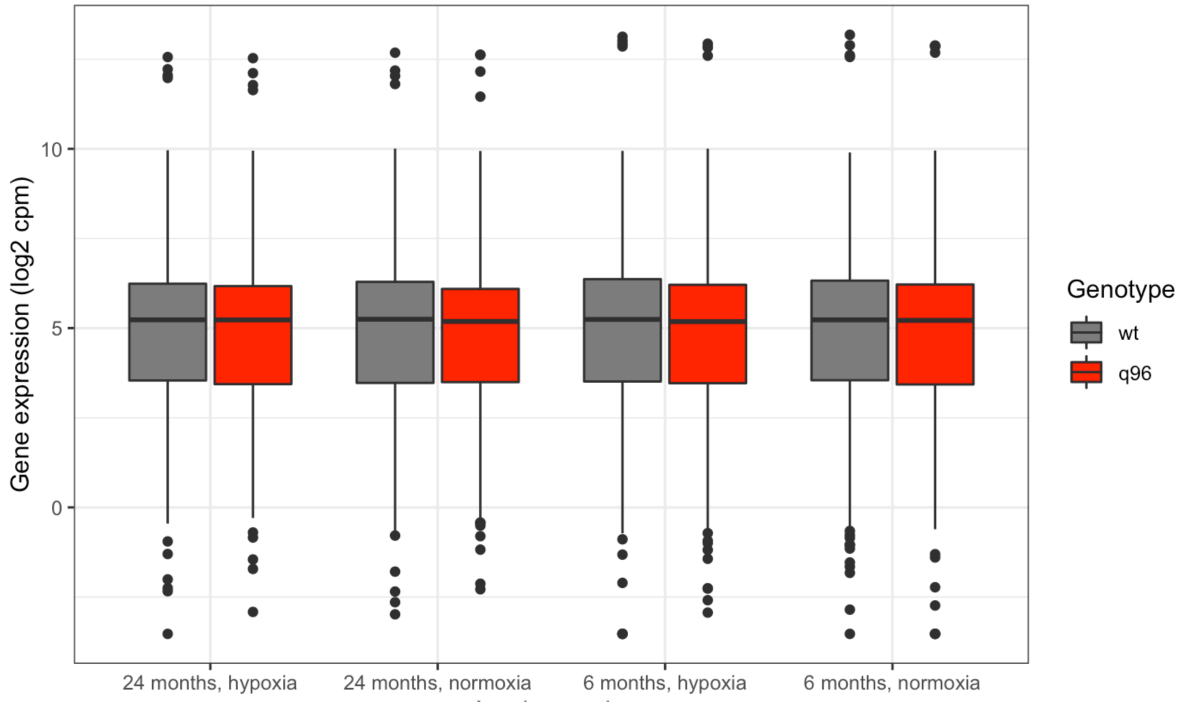


**Age- and hypoxia- dependent expression of neural marker genes (microglia, astrocyte, neuron, oligodendrocyte) in fAD-like zebrafish whole-brain RNA-seq dataset analysed.**

There is a possibility that changes in gene expression between may reflect changes in the proportions of cell types rather than changes in expression in each cell. For example, less neurons might be expected during neurodegeneration. To explore the possible changes in the proportions of cell types, we looked in marker genes of three common neural cell types: astrocytes, neurons, microglia, and oligodendrocytes.

The marker genes for astrocytes, neurons, and oligodendrocytes were derived from gene sets by Lein et al. (2007) and are based on studies in mice (available on MSigDB). The marker genes for microglia were derived from Bonham et al. (2019) and based on studies in human and mouse. All gene IDs were converted to zebrafish Ensembl IDs using BioMart.

Overall, expression of these neural marker genes appears to be stable across different conditions and ages.
